# An inside look at a biofilm: Pseudomonas aeruginosa flagella bio-tracking

**DOI:** 10.1101/2020.08.26.267963

**Authors:** Eden Ozer, Karin Yaniv, Einat Chetrit, Anastasya Boyarski, Michael M. Meijler, Ronen Berkovich, Ariel Kushmaro, Lital Alfonta

## Abstract

The opportunistic pathogen, *Pseudomonas aeruginosa*, a flagellated bacterium, is one of the top model organisms for studying biofilm formation. In order to elucidate the role of the bacteria flagella in biofilm formation, we developed a new tool for flagella bio-tracking. We have site-specifically labeled the bacterial flagella by incorporating an unnatural amino acid into the flagella monomer via genetic code expansion. This enabled us to label and track the bacterial flagella during biofilm maturation. Direct, live imaging revealed for the first-time presence and synthesis of flagella throughout the biofilm lifecycle. To ascertain the possible role of the flagella in the strength of a biofilm we produced a “flagella knockout” strain and compared its biofilm to that of the wild type strain. Results showed a one order of magnitude stronger biofilm structure in the wild type in comparison to the flagella knockout strain. This suggests a newly discovered structural role for bacterial flagella in biofilm structure, possibly acting as a scaffold. Based on our findings we suggest a new model for biofilm maturation dynamic and underscore the importance of direct evidence from within the biofilm.

## Main

*Pseudomonas aeruginosa* (*P. aeruginosa*) is a well-studied opportunistic pathogen^1^. Despite the fact that quorum-sensing and biofilm formation have been studied for the past 40 years, new information is constantly being reported^2,3^. Biofilms provide a more resistant form of existence for bacteria than their planktonic forms, proffering them with protection from possible stressors. They therefore, have been intensely studied for their complexity and the mechanisms involved in their life cycle^4,5^. Indeed, any new information emerging from these studies is crucial, allowing the development of new and diverse strategies to resist infections.

The biofilm lifecycle is composed of several commonly reported steps^1,6–8^. Initially, planktonic bacteria propel themselves to a proximal surface, followed by an irreversible attachment to the surface. Once attachment is established, exo-polymeric substances are secreted from within the cells to generate a matrix of a supporting microenvironment for the dividing cells and to initiate formation of micro-colonies. Next, mushroom-like structures start to emerge. Finally, the cells secrete enzymes to digest the exo-polymeric substances at the top of the grown mushroom-like structures where the newly flagellated cells are released in a planktonic form to attach to new exposed surfaces.

The flagella, the bacterial rotor with its unique structure, is therefore an inseparable part of biofilm research^9^. Several reports have expressed a consensus regarding the importance of flagella in biofilm formation, specifically in its initiation^10^. However, despite a consensus among researchers that flagella are not present during biofilm maturation but only in the dispersion stage, reports regarding this are still somewhat constradicting^10–16^. Therefore, it is necessary to assess the presence and possible role of flagella in biofilms. Direct imaging of these organelles inside a developing biofilm may thus shed a better light on their role in biofilm formation and maintenance as well as provide an improved understanding of what occurs during the biofilm lifecycle.

To date, live-cell imaging can be obtained through different approaches, however genetic code expansion, the reassignment of codons and incorporation of an unnatural amino acid (Uaa) into proteins^17^, displays advantages over other methodologies, and is gaining increasing exposure and momentum^18–21^. Genetic code expansion systems are being constantly improved, expanded and adapted to a growing number of organisms^22–25^. This technique aids in improving imaging. For example, the incorporation of a Uaa may alleviate the need for large and bulky labeling agents, such as fluorescent proteins or antibodies. Moreover, a protein with a site-specifically incorporated Uaa can be labeled using bio-orthogonal chemistry and serve as a specific reporter inside cells. Incorporating Uaa into *P. aeruginosa* flagellum enables live-cell imaging of flagella inside the complex environment of a biofilm. Herein, we present a robust and orthogonal genetic code expansion system in *P. aeruginosa* designed for flagella labeling for *in-vivo* flagella bio-tracking in a live and growing biofilm that revealed novel information regarding the biofilm lifecycle.

## Results

### Genetic code expansion of *P. aeruginosa*

For a Uaa incorporation into *P. aeruginosa* proteins, a new plasmid was constructed (Fig. 1a). The plasmid harbors an orthogonal translation system (OTS), as well as a reporter gene for system validation. An OTS, composed of a tRNA and tRNA-synthetase pair, could be considered orthogonal if it does not interact with native translational components^26^. *Methanosarcina mazei* pyrrolysyl orthogonal translation system^27^ (*Mm*Pyl OTS) has been previously found to be orthogonal in several organisms, including gram-negative bacteria, and was therefore our choice. Combined with *Mm*Pyl OTS, a GFP reporter gene encoded for Uaa incorporation.

**Figure 1.**
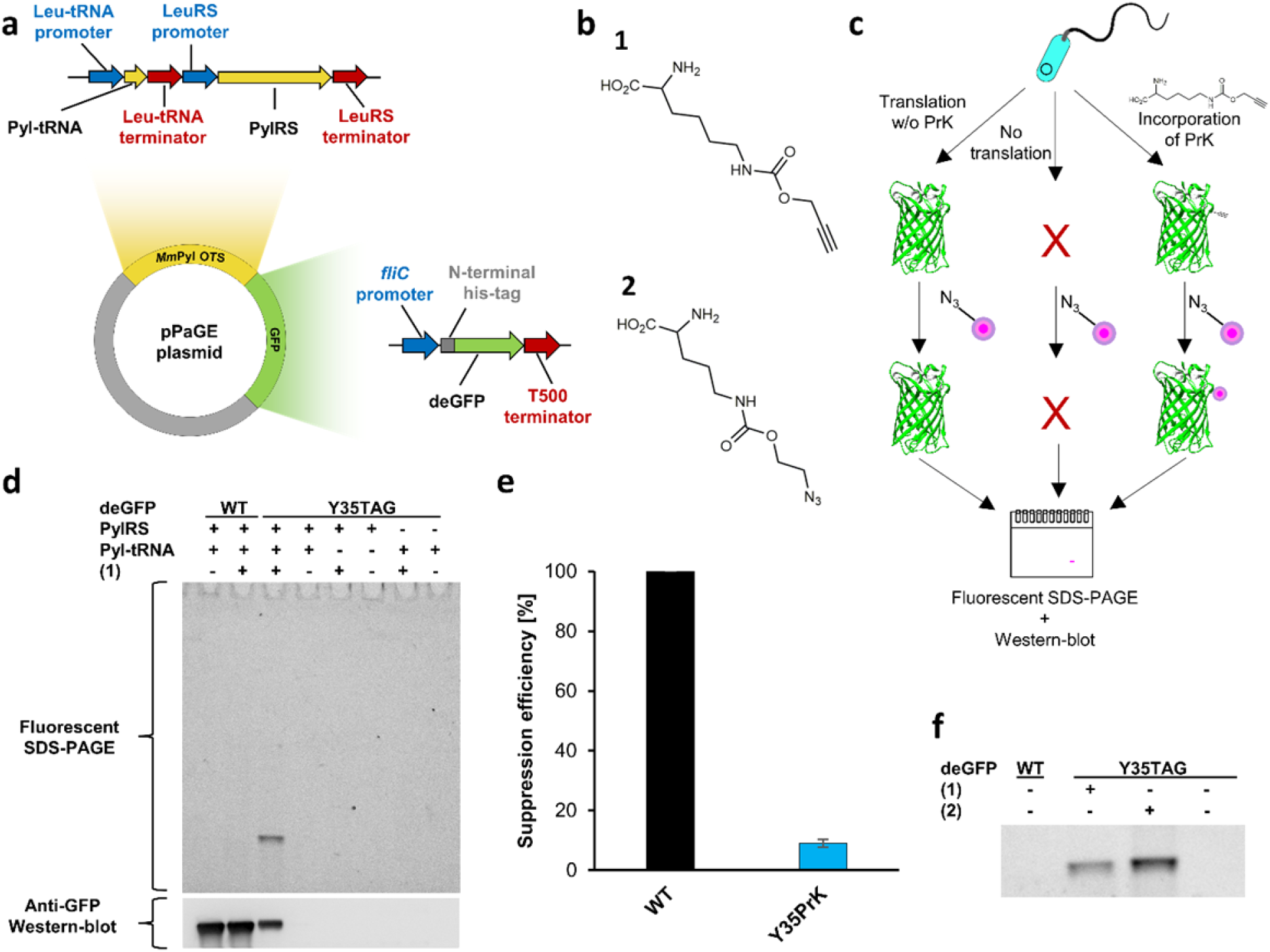
Genetic code expansion system of *P. aeruginosa* using GFP. (a) Plasmid map of pPaGE. (b) Uaas used in this study, PrK (**1**) and AzCK (**2**). (c) Schematic representation of possible translation outcomes for mid-gene TAG mutants. (d) Lysed samples following a click reaction to an azide-containing fluorophore analyzed through fluorescent SDS-PAGE and anti-GFP Western-blot analyses. (e) Suppression efficiency of GFP Y35PrK mutant. (f) Lysed samples following a click reaction to either an azide or an alkyne containing fluorophore analyzed through fluorescent SDS-PAGE.

The *P. aeruginosa* genetic code expansion plasmid (pPaGE) was assembled using *P. aeruginosa* endogenous promoters and terminators. Our rationale was that the bacterium will benefit from an attempt to maintain physiologically relevant expression levels of an exogenous OTS. We thus identified the most abundant codon in *P. aeruginosa* PAO1 genome^28^, which was found to be the CUG codon encoding for leucine (Leu). The native promoters and terminators of Leu-tRNA and Leucil-tRNA-synthetase were assigned as the upstream and downstream regions of Pyl-tRNA and Pyl-tRNA-synthetase respectively. Planning a future expression of flagella protein, the *fliC* endogenous promoter for flagellin expression, was chosen to serve as a promoter for the GFP reporter in the pPaGE plasmid. Toxicity tests for the Uaas: propargyl-L-lysine (PrK (**1**)) and azido-carboxy-lysine (AzCK (**2**)) (Fig. 1b.1 and Fig. 1b.2, respectively) were performed and have shown normal growth rates (Fig. S1).

We reassigned the TAG stop-codon for site-specific incorporation of Uaas and introduced a TAG mutation to the GFP reporter gene at its 35^th^ site. *Mm*Pyl OTS orthogonality in *P. aeruginosa* was tested next. Several protein elongation scenarios for a premature TAG introduction to GFP were tested, as well as the possibility of non-specific incorporation of a Uaa into the host organism proteome (Fig. 1c, Fig. 1d). For that purpose, pPaGE variants containing partial OTSs were generated by the removal of Pyl-tRNA or Pyl-tRNA-synthetase (PylRS). Cells were grown, lysed and analyzed using fluorescent chemical conjugation and anti-GFP Western-blot analyses (Fig. 1d). In order to incorporate (**1**) into GFP, a click reaction using Cu(I)-catalyzed azide-alkyne cycloaddition (CuAAC)^29^ to an azide-bearing fluorophore was conducted. Since there are no endogenous alkynes in bacteria, this experiment was meant to reveal whether there exists any (**1**) that was misincorporated in response to the TAG stop-codon in the bacterial proteome. Another possibility that needed to be ruled out was if PylRS can aminoacylate endogenous tRNAs, where (**1**) would have been incorporated into random locations in the genome. Both scenarios, resulting in non-specific fluorescent labeling into endogenous proteins. On the other hand, if Pyl-tRNA is not orthogonal and is aminoacylated by the host organism’s tRNA-synthetases by natural amino acids, full-length GFP may still be synthesized and observed in the Western-blot.

When examining all the protein expression options as seen in Figure 1d, we did not observe fluorescent labeling or GFP expression in the presence of the partial OTS variants. This directly proved *Mm*Pyl OTS’s orthogonality for the first time. Following the establishment that our system is indeed orthogonal, we were then able to analyze proper Uaa incorporation into GFP. Wild type (WT) GFP expression was observed either in the presence or absence of (**1**) and could be seen only in the Western-blot analysis and not in the fluorescent gel. This indicated that there was no incorporation of (**1**) into WT GFP. When the whole OTS was present together with (**1**), a fully elongated GFP was detected in the Western-blot analysis. In addition, a clear fluorescent labeling corresponding to GFP in size was visible. This not only indicated that the reporter gene was able to be synthesized but also verified the presence of (**1**) inside the protein. Indeed, when those cells were grown in the absence of (**1**), no expression was observed, signifying that the expressed protein was not a result of a read-through event by any natural amino acid. Final validation was performed using electrospray mass spectrometry (ESI-MS) (Fig. S2), where the deconvoluted mass corresponded to GFP molecular weight with the desired (**1**) at position 35 instead of a tyrosine in the WT protein. Thus, it was concluded that Uaa was successfully incorporated and encoded for, in a protein expressed in *P. aeruginosa*.

Suppression efficiency for Uaa incorporation into GFP was calculated as just under 10% (Fig. 1e). While offering information on the incorporation efficiency, it is important to note the context effects^30^ at play in this methodology which may alter incorporation yields as well as suppression efficiency. For example, having the *fliC* promoter may impair incorporation into GFP but may be better for Uaa incorporation into a flagellin. In order to further establish the generated system following genetic code expansion with (**1**), (**2**) was also incorporated into GFP to test system’s flexibility, moreover (**2**) has the ability to undergo a copper-free click chemistry^29^ which makes it useful for future *in-vivo* applications. Here, (**2**) was also detected through copper-catalyzed click chemistry, this time to an alkyne-containing fluorophore, and resulted in positive incorporation (Fig. 1f). Thus, genetic code expansion was successfully achieved and the system’s orthogonality was established in *P. aeruginosa*, using two different Uaas.

### Flagella “bio-tracking” in a biofilm

Next, we incorporated a Uaa into the endogenous gene of flagella. The flagellum is a complex machinery composed of several genes and proteins that generates filaments extending up to 15 μm in length^31,32^. Although assembled by a consortium of proteins, the flagellin protein encoded by the *fliC* gene is the repeating monomer subunit giving the flagellum its high aspect ratio. We decided to introduce modifications mainly in the D3 domain of flagellin, facing outwards in the cylindrical structure of flagellum, thus minimizing possible interference with filament assembly^32^. We modulated the *P. aeruginosa* flagellum that is composed of 41 subunits (based on PDB accession number 5wk5) with (**1**) incorporated at flagellin’s 248^th^ position (Fig. 2a). This illustrates our ability to label the micro-structure by using a Uaa as a reactive bioorthogonal chemical handle, holding vast potential for bio-labeling and other applications.

**Figure 2.**
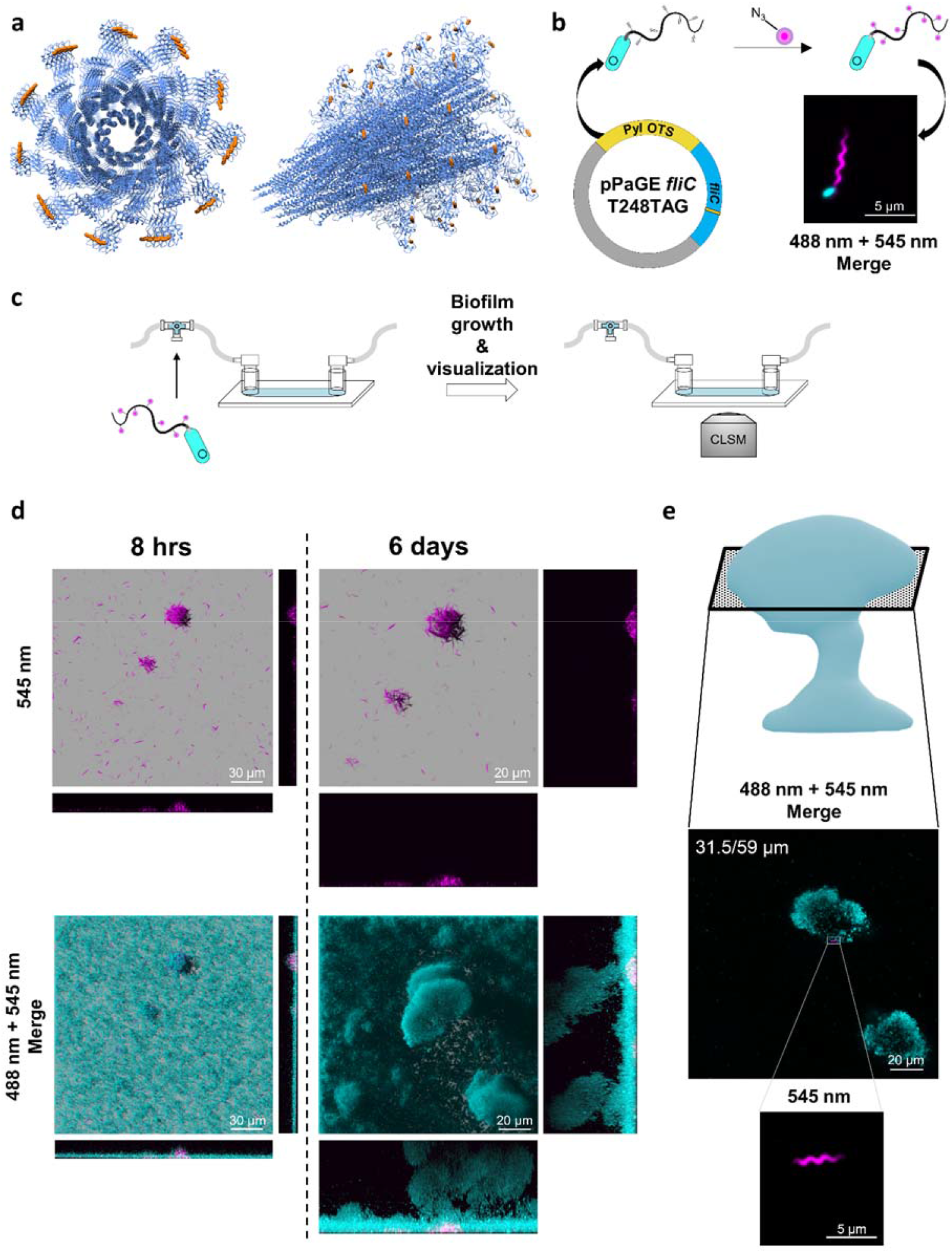
Flagellin Uaa incorporation and inoculated flagella survival in a biofilm. (a) Theoretical model of *P. aeruginosa* flagella filament with incorporated (**1**) at the D3 domain (based on PDB accession number: 5WK5) left: a top view; right: a side view, Uaa is an orange sphere. (b) Predicted system functionality and resulting fluorescent imaging using confocal laser scanning microscopy (CLSM). Flagella with an incorporated Uaa were labeled fluorescently (magenta) using PAO1 encoding genomic GFP reporter (gGFP) (cyan). Planktonic bacteria were imaged with incorporated (**1**) in the flagellum. (c) Schematic representation of the flow-cell and experimental setup for bio-tracking of flagella used for biofilm inoculation throughout biofilm maturation. Flagella with a Uaa were fluorescently labeled and used as inoculation cells for continuous biofilm growth for up to 6 days. Biofilm development and flagella were followed using CLSM. (d) Inoculated bacteria in a biofilm, pre-labeled prior to inoculation, monitored inside biofilm’s 3D structure development using CLSM. (e) Pre-labeled flagella located in a mature biofilm at mid-height after 6 days of growth. Full scale and resolution images are available in the supplementary information (SI) file Figs. S6-S11.

We reasoned that replacing the native flagellin copy into the genome of *P. aeruginosa* with an incorporated Uaa may result in a short flagellum, leading to artifacts, as Uaa incorporation is a slower process than native amino acids incorporation^33^. For that reason, we chose an integrative approach: while encoding for a Uaa incorporation into the *fliC gene* harbored in the pPaGE plasmid, we still retained native flagellin in the genome for a hybrid assembly into a single unified flagellum. The integrated flagellum was thus predicted to have fewer chemical anchor points for bio-labeling. However, despite not having every monomer subunit carry a Uaa, this technique still allowed the micro-fiber to be labeled. Using the *fliC* promoter, the downstream sequence from the inserted *fliC gene* was also chosen as the native flagellin terminator, hence the sequence was completely identical to the native genomic sequence apart from the introduced premature TAG stop-codon mutation (Fig. 2b). The new plasmid was assembled and served for flagella genetic code expansion from that point onwards.

pPaGE *fliC* was transformed into *P. aeruginosa* and resulted in (**1**) incorporation into flagellin monomers (Fig. S3). It was also established that exogenous flagellin expression does not affect flagellum synthesis in each individual cell (Fig. S4). Next, the system was used for live-cell imaging using a click reaction. pPaGE *fliC* was inserted into *P. aeruginosa* PAO1 gGFP strain, carrying a genomic GFP reporter gene for the convenience of whole-cell detection. Using a confocal laser scanning microscope (CLSM), successfully labeled flagella were observed including its unique wave-like morphology features indicating a repetitive occurrence of the incorporated label (Fig. 2b, magenta). No signal was observed when expression was attempted in the absence of (**1**) (Fig. S5). Despite the fact that not every monomer was labeled, the generated filament was successfully visualized, and the unique wave-like feature of flagella was recorded.

With this new labeling tool, it was now possible to “bio-track” flagella inside a biofilm. This technique enabled us to track the labeled inoculated flagella, attached to the cells that were used to initiate biofilm formation. Hence, the following experimental setup was pursued (Fig. 2c): *P. aeruginosa* expressing genomic GFP, harboring the pPaGE *fliC* plasmid were grown in the presence of (**1**). Planktonic bacteria containing (**1**) were fluorescently labeled by a click reaction and inserted into a flow-cell. Labeled bacteria were grown for up to 6 days, while inoculated flagella were monitored inside the biofilm every few hours. Labeled flagella, with their distinct morphology, were clearly observed after initial surface attachment and growth in the flow-cell either as singles or as clusters of different sizes (Fig. 2d). The emergence of flagella clusters was a notable observation that was not reported before.

Flagella bio-tracking for a time period of up to 6 days of biofilm growth, enabled us to provide evidence regarding possible changes in the state of the flagella in a biofilm with time. For example, we could determine whether inoculated flagella are maintained or degraded within the biofilm. We could demonstrate in a direct manner that despite continued biofilm growth to heights of above 59 μm in total, the majority of flagella remained in approximately at the bottom of the biofilm in the first 13 μm of the biofilm (Fig. 2d). While most flagella seemed to remain at the bottom of the biofilm, rarely we observed inoculated flagella in higher sections of a grown biofilm, found at around 31.5 μm (Fig. 2e). That means that not only inoculated labeled flagella are still present and are not metabolized within the time frame of the experiment but also that occasionally, they might reach higher regions in the biofilm through an unknown mechanism. The notion of flagella movement inside the biofilm has been previously speculated in the literature^13,14^, however this is the first direct evidence for its occurrence. Indeed, the images presented here (Fig. 2e) serve as the first direct evidence to this hypothesis. It is important to note that due to imaging limitations, it is hard to determine if the observed flagella are still attached to a cell or not. However, individual flagella that are observed in the biofilm’s mid-height were most likely positioned there due to a response to bacterial signaling. Hence, this has led us to believe that most of the observed flagella are indeed attached to bacterial cells.

### Flagella synthesis in a biofilm

After locating the labeled inoculated flagella in the biofilm, we wanted to ascertain whether we could track newly synthesized flagella in the biofilm and if so, where are they located in the biofilm. The biofilm lifecycle is divided into several steps where planktonic cells transform and grow together into mushroom-like structures. Interestingly, throughout the biofilm’s growth, flagella synthesis is halted and is only re-initiated during the final step of dispersion from the mature biofilm^5,7,8,34–36^. To date, the most accepted concept of flagella synthesis in a biofilm is that it occurs in a compartment located at the upper section of the mushroom-like structure in a grown biofilm. Therefore, using our system, we set out to label the compartments at the top of the mushroom-like structures in order to directly visualize flagella inside them for the first time. Pre-labeled cells were used for flow-cell inoculation as before, but this time the media supply for the flow-cell contained (**1**) for continued Uaa incorporation inside the growing biofilm for 2,4 or 6 days. Following growth, the biofilm was re-labeled using the same fluorophore (Fig. 3a).

**Figure 3.**
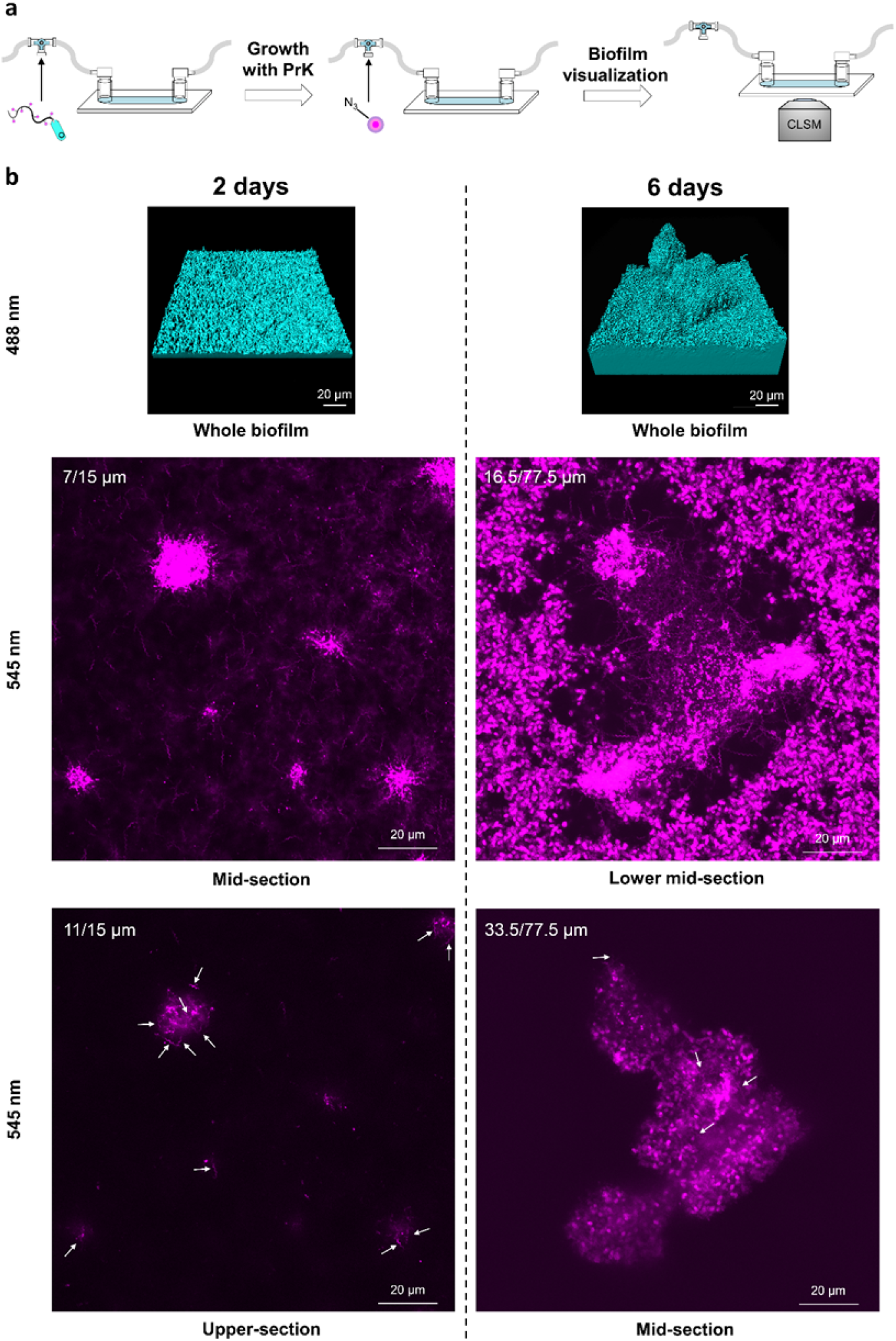
Flagella synthesis during biofilm maturation. (a) Schematic representation of the experimental setup identifying flagella synthesis within the biofilm. Flagella with incorporated Uaa were fluorescently labeled and used as inoculation for continuous biofilm growth in the presence of Uaa for up to 6 days. Flagella were fluorescently labeled inside the biofilm at different biofilm’s growth time points of 2 and 6 days and visualized using CLSM. (b) CLSM imaging from within the biofilm at different growth stages. For each time point during biofilm maturation, biofilm was imaged, and flagella were documented at different heights of the biofilm. Bacteria labeled in cyan and flagella in magenta. Enlarged images, as well as four-day old biofilm images are available in Figs. S12-S26.

Large quantities of fluorophore were trapped inside the viscous exo-polymeric matrix of the biofilm^6^ and appeared as a bulk cloud-like signal. Flagella did not appear at the top of the mature biofilm, while many flagella were observed at the lower and middle sections of the biofilm throughout its growth (Fig. 3b, images of 4 days old biofilm and Figs. S17-S21). In the two-day old biofilm, flagella were located throughout the entire biofilm up to the height of 15 μm on average. A four-day old biofilm also had flagella at the base of the biofilm as well at the height of approximately 22 μm out of 30.5 μm in total. Finally, a mature six-day old biofilm had abundant and noticeable flagella at mid-height (33.5 μm) of the 77.5 μm maximal height. These results not only confirmed our prediction of being able to detect flagella in the biofilm throughout its unique shape, but also contradicted the accepted consensus that flagella are located only at the top of mushroom-like structures in a mature biofilm.

Consistently seeing numerous flagella inside the biofilm throughout its whole lifecycle was unexpected. This is because according to the literature^5,7,8,34–36^, it appears that there is no flagella synthesis during biofilm growth. This notion was based mainly on indirect bulk-population analysis of DNA/RNA microarrays, total biomass calculations and proteomic analysis^15,37–39^. This discrepancy provides important evidence for the importance of direct flagella visualization from within the biofilm, at an appropriate spatio-temporal resolution. When new flagella were observed from the biofilm’s base up to approximately mid-height, we could only assume that previously indirect analysis had an averaging effect on the population. Another accepted aspect was that flagella exist at the top of micro-colonies in young biofilms, to allow flagella mediated movement in between micro-colonies^13,14^. A compartment containing flagella at the top of a mushroom-like structure in a grown biofilm or any flagella in higher regions were not observed even though they were easily detectable everywhere else in the biofilm.

Cells may detach from the mature biofilm by several ways; through erosion, bulk detachment or by planktonic release^8^. In order to ascertain the possible changes occurring in detaching cells, the Flow-cells’ effluent was collected prior to whole biofilm labeling and was labeled using a second fluorophore (with different excitation and emission wavelengths) to distinguish between inoculated flagella and newly synthesized flagella (Fig. 4a). We collected the biomass into chilled tubes immediately upon their exit from the flow-cells. Thus, we could assume that any flagellum that was observed was synthesized inside the biofilm. Looking at the double-labeled effluent of a young biofilm, new flagella were clear and abundant. This meant that cells are frequently flagellated before leaving the biofilm (Fig. 4b). While correlating well with known dispersion mechanisms from a mature biofilm, we expected to observe flagellated cells leaving only the mature biofilm and were surprised by the continued flow of flagellated cells in the effluent of our flow-cell set-up. (Fig. 4b, Figs. S27-S38).

**Figure 4.**
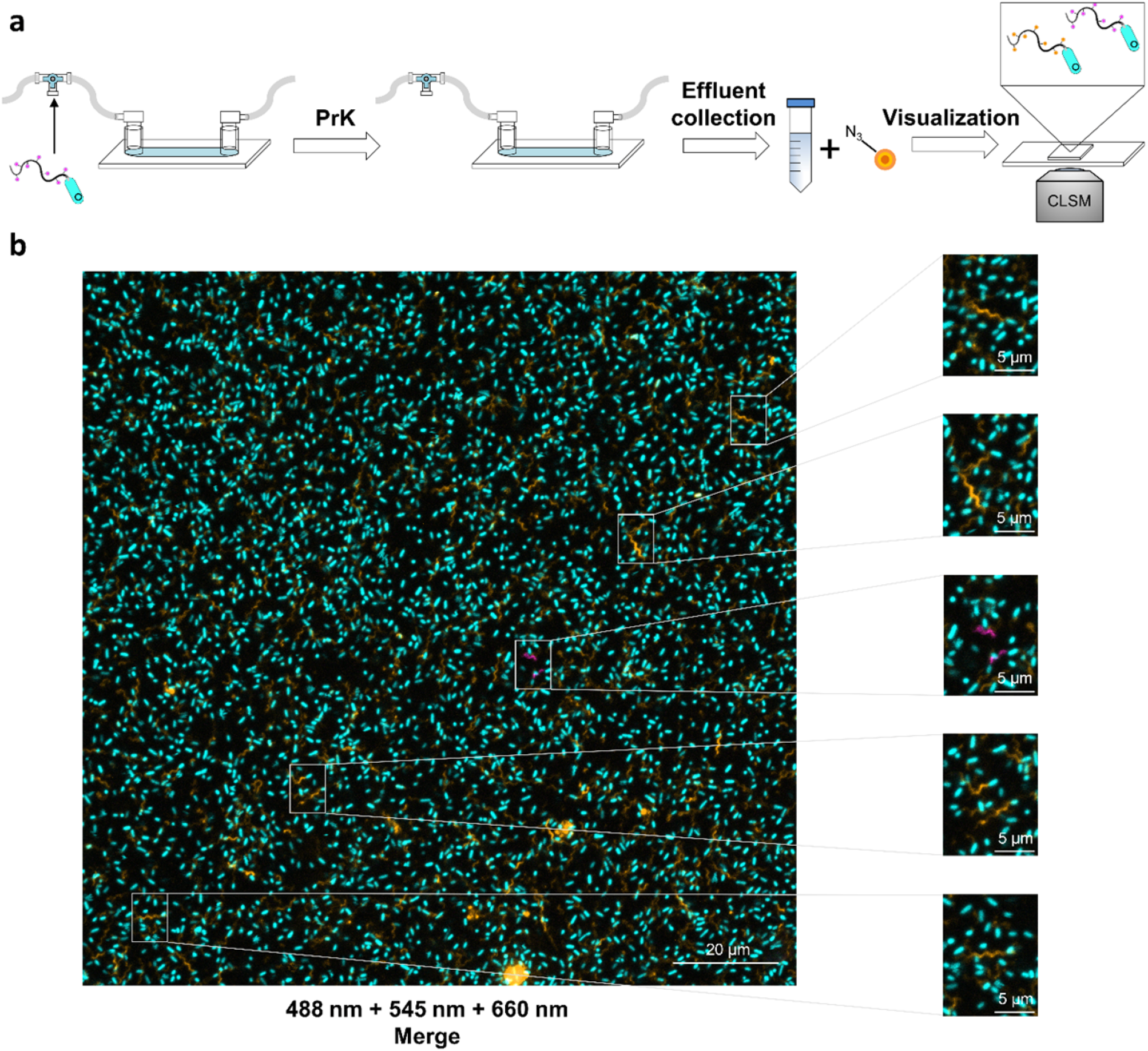
Dispersed flagellated cells. (a) Schematic representation of the experimental setup identifying dispersed cells from within the biofilm carrying synthesized flagella from before their release. Flagella with incorporated Uaa were fluorescently labeled and used as inoculation for continuous biofilm growth in the presence of Uaa for up to 6 days. Effluent was collected at different time points, labeled with a second, different fluorescent dye and visualized using CLSM. (b) CLSM imaging of dispersed flagellated cells from a two-day old biofilm. Bacteria labeled in cyan, inoculated flagella in magenta and new flagella in orange. Enlarged images, as well as dispersion images for 4-/6-day old biofilms, are available as Figs. S27-S38.

Examining many flagellated cells dispersed from the biofilm, we also detected very rare cases of labeled inoculated flagella (Fig. 4b). Although these observations were rare, indicating that this may be a statistical error when analyzed quantitatively, the image was clear and reproducible. As mentioned previously, it is difficult to detect if the flagella that were observed in the biofilm were connected to cells or not. Despite this, we showed that labeled inoculated flagella were still attached to cells. This served as further support of our initial hypothesis, suggesting most of the observed flagella were indeed still connected to a cell and that they displayed some level of movement inside the biofilm.

Discovering continued flagellated dispersion is novel information regarding the biofilm lifecycle model and reflects the observation that the biofilm is filled with flagella as was discovered in this study. From a clinical aspect, mature biofilms are prone to planktonic dispersion at times, causing exacerbations in chronic infections and afflicting new environment within the host^40^. Usually, these exacerbations are in need of antibiotics treatment but are only taken under consideration in case of a mature biofilm. Hence, continuous planktonic cells release could affect the clinical view of possible treatments for chronic “biofilm infected” patients. All the information gathered in this work, enabled us to portray an updated and more accurate model for the biofilm lifecycle (Fig. 5).

**Figure 5.**
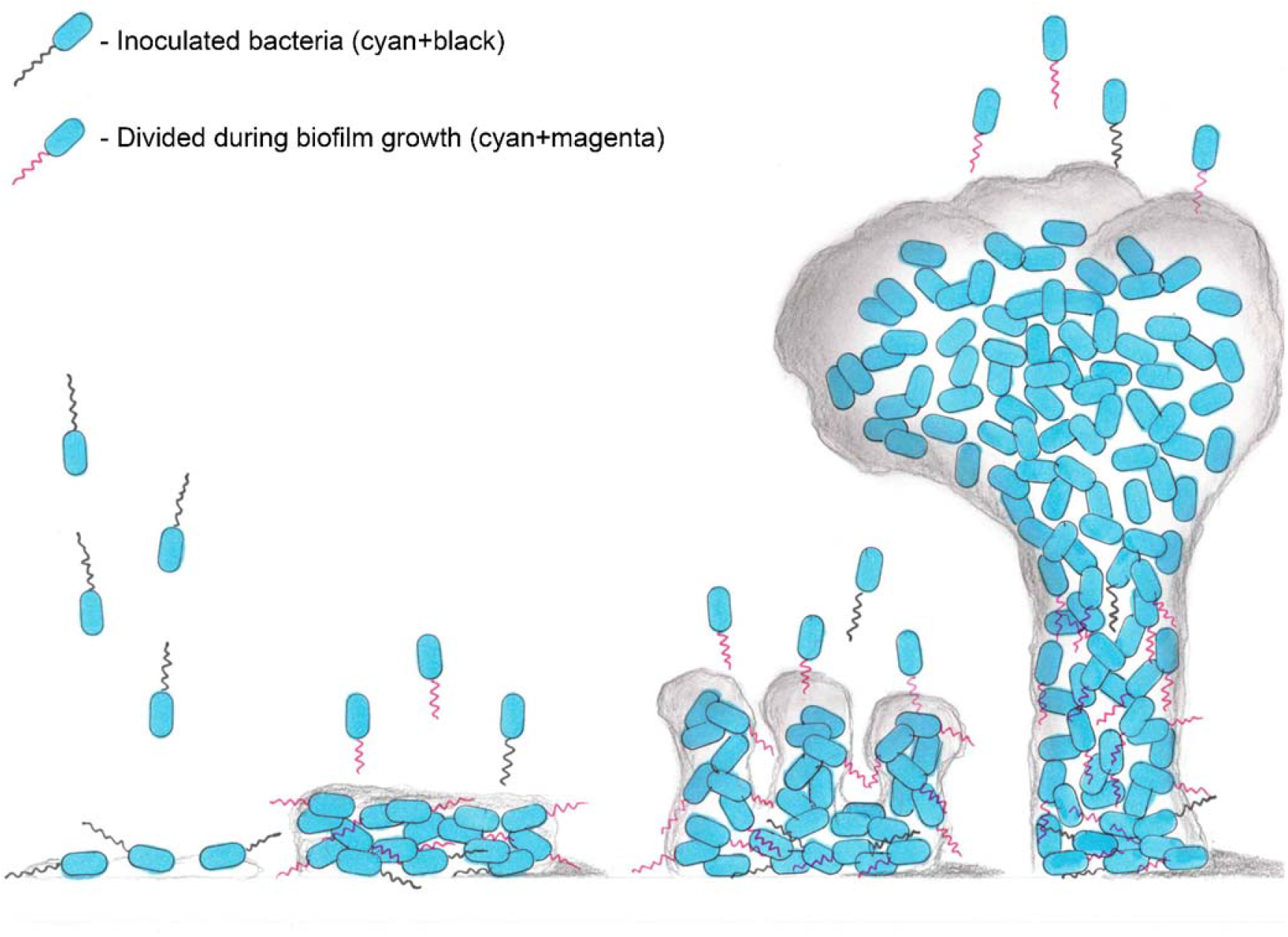
Updated biofilm lifecycle model based on the presented discoveries.

### Flagella’s structural importance in a biofilm

Learning that flagella are continuously being synthesized within a growing biofilm, we asked ourselves whether they might play a possible role in a biofilm mechanical structure. Considering that flagella synthesis is energetically costly^41^ and that cells within the biofilm are mainly non-motile, there must be a reasonable explanation as to why the flagella are formed in cells inside the biofilm. We observed a visible grid-like appearance of the flagella in the lower part of the biofilm (Fig. 6a). In addition, we observed several bacteria assembling on a single flagellum as shown in Figure 6b. These observations seemed of importance as the structure resembled a structural scaffold that supports a substantial architecture. The idea of the role of flagella as a mechanical support in a biofilm has been previously suggested in *E. coli* macrocolony biofilms but has yet to be demonstrated^42^. An additional observation to support this hypothesis is the observation of *Geobacter sulfurreducens* electrochemical-chamber biofilm in their cytochromes spatial arrangement on the flagella^12^. To investigate such a possible role of flagella for physical support, we constructed a *P. aeruginosa* strain that allows us to examine this notion.

**Figure 6.**
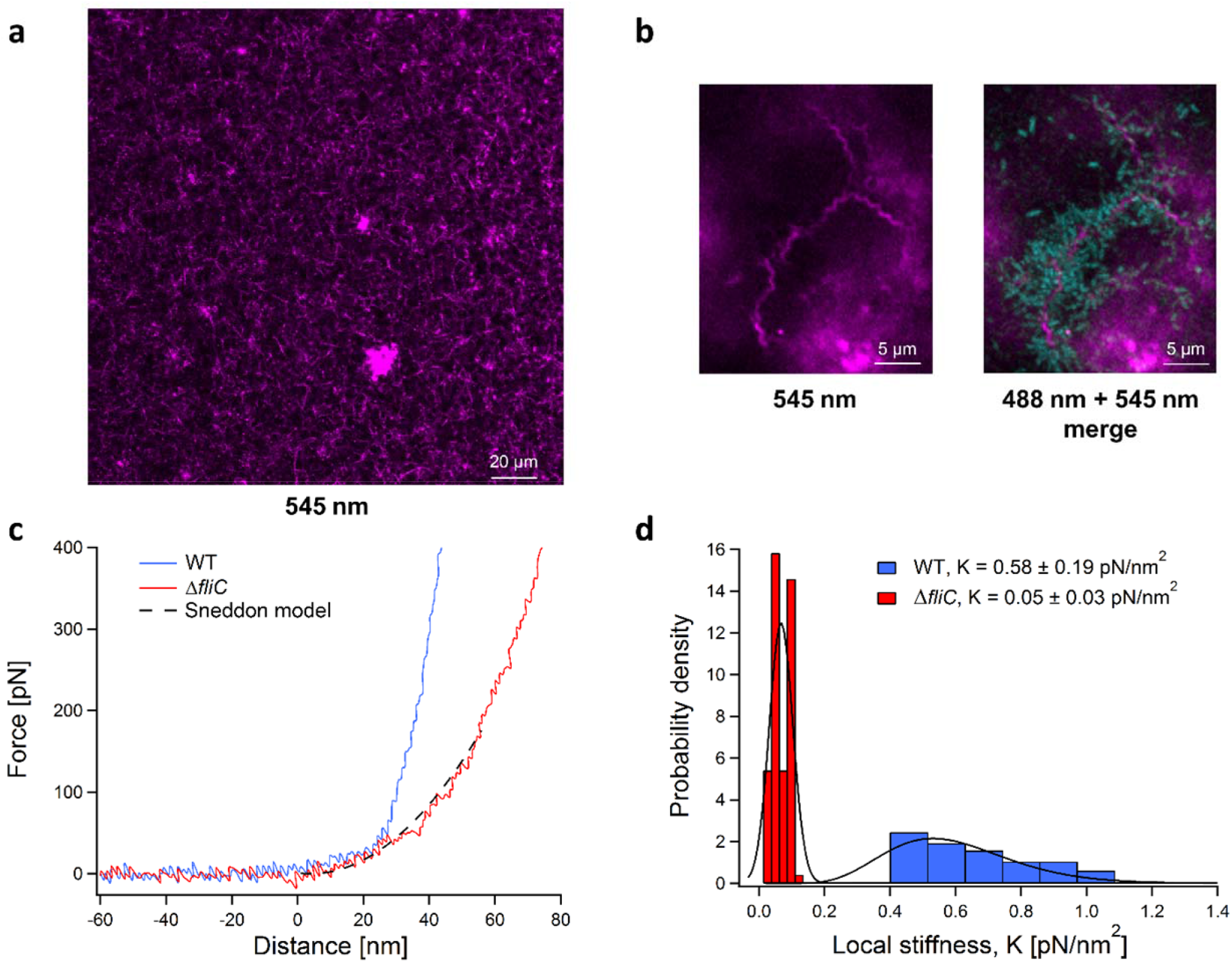
Flagella as physical support in a biofilm microenvironment. (a) CLSM imaging of the lower section of a four-day old biofilm with numerous flagella in a grid-like structure. (b) CLSM imaging of bacterial assembly on flagella in a growing biofilm. Bacteria labeled in cyan and flagella in magenta. Enlarged images are available as Figs. S39-S40. (c) Nanoindentation force-distance curves of the WT biofilm (blue) and a Δ*fliC* biofilm (red). The Δ*fliC* curve is fitted with the Sneddon model (dashed black line), from which K, the local stiffness modulus was estimated. (d) Local stiffness probability density distributions of WT biofilm (blue) and Δ*fliC* biofilm (red), The distributions were fitted by relevant statistical models (black line), from which their means and standard deviations were calculated.

Using CRISPR Cas9 for genome editing, based on *Streptococcus pyogenes* Cas9 which was adapted for *P. aeruginosa* genome editing^43^, we generated a sub-strain lacking the ability to produce flagella. This was carried out by deleting the flagellin gene (Δ*fliC*) from the *P. aeruginosa* genome. Transmission electron microscope (TEM) imaging and the loss of its swarming abilities validated that the new strain does not express flagella (Fig. S41). We then attempted to identify differences between WT and Δ*fliC* based biofilms. We hypothesized that mushroom-like structures will be affected by the absence of flagella. Despite this, we could not detect any distinct gross morphological differences between the two biofilms (Fig. S42). However, high-resolution scanning electron microscopy (HR-SEM) revealed noticeable visual difference between the two biofilms (Fig. S43). WT biofilm possessed fibers forming a web between the cells, while in the Δ*fliC* biofilm cells were much more crowded and there were significantly less fibers. Since multiple string-like fibers were present in the biofilm and represented the complex environment surrounding the cells (sugars, proteins, DNA, etc.) it was not possible to use regular microscopic methods to determine morphologically which elements are flagella and which are not. Therefore, further comparison was performed using atomic force microscopy (AFM) on WT and Δ*fliC* biofilms (Fig. S44). Though we did not find an indication in these images for the structural importance of the flagella to the biofilm, flagella were easily recognized within the WT biofilm. This serves as further validation of the consistent presence of flagella in the biofilm.

Hypothesizing that flagella contribute to the strength of the biofilm’s structure, we decided to inspect the biofilm’s mechanical strength and stiffness. A biofilm’s mechanical strength is important to its survival. This characteristic is being actively studied in order to better understand the contribution of different elements to the exo-polymeric environment or to measure biofilm’s resilience following different treatments^44–46^. For that purpose, we used AFM nanoindentation measurements on the two variants’ mature biofilms. If the hypothesis is correct and the flagella take part in the biofilms’ structural strength, then a biofilm lacking flagella should be mechanically weaker. Figure 6c shows examples of WT and Δ*fliC* (marked as blue and red, respectively) biofilm force-distance traces obtained from the nanoindentation measurements. Each indentation curve demonstrates specific elastic response. There is a clear difference in the elastic response curvatures for the two biofilms indicating lower local stiffness for the Δ*fliC* biofilm. Fitting the curves with the Sneddon model allowed the estimation of local elastic stiffness, K, as illustrated for the Δ*fliC* biofilm (Fig. 6c black dashed line).

We analyzed 100 force-distance curves collected for each of the biofilms, enabling the estimation of local stiffness and verification that the biofilms did not undergo irreversible deformation. The local stiffness, K, was collated into probability density functions that interestingly displayed two spreading behaviors. While the stiffness of the Δ*fliC* displayed a narrow normal distribution, the stiffness of the WT biofilm showed a wide heavy tail distribution. This means that the stiffness values of the WT biofilm spread over a wider range compared to those of the Δ*fliC* biofilm, with stiffness values that can get considerably high. For this reason, the Δ*fliC* K probability distribution was fitted with a normal (Gaussian) distribution, and the WT K was fitted with Gamma distribution, in order to properly assess their collective values, i.e., means and standard deviations (Fig. 6d). Local stiffness value estimated for WT biofilm was one order of magnitude higher than Δ*fliC* biofilm’s local stiffness. This proved that the Δ*fliC* biofilm is weaker in terms of physical strength (K_WT_ = 0.58 ± 0.19 pN/nm^2^ vs. K_Δ*fliC*_ = 0.05 ± 0.03 pN/nm^2^). This result implied that, indeed, flagella take a crucial part in the biofilm mechanical strength, much like scaffolds in a construction.

In order to exemplify the importance of these findings we returned to the literature and found that in some cases, *P. aeruginosa* strains isolated from cystic fibrosis patients lack flagella due to various mutations in flagella synthesis involved genes^47^. It was found that cystic fibrosis isolates occasionally develop differently, in order to evade the human immune system that mainly targets bacterial flagella. Recently, Harrison et. al.^47^ revealed the connection between the loss of flagellum and overexpression of exopolysaccharides in biofilms created from cystic fibrosis isolates. The nanoindentation results presented here can elucidate further details on this fascinating mechanism, as it is possible that bacteria produce more exopolysaccharides in the attempt to compensate for the loss of flagella and its mechanical support. Also, if flagella indeed contribute to the biofilm’s strength, future biofilm treatment approaches may potentially target flagella, to weaken the biofilm’s structure and improve antibiotics penetration. Such a strategy may then be followed by an effective antibiotic treatment that would otherwise be less efficient in a strong and fully functional biofilm, and give way to future studies in clinical/environmental biofilms.

## Discussion

The field of biofilm studies is of great importance for basic microbiological understanding as well as in medical applications. Being studied intensely for over 40 years there are still many pieces missing in the complex biofilm puzzle. Despite the vast interest, part of the reason for this great mystery is its complexity and the challenges in its manipulation. For the most part, biofilm research is performed through bulk population analysis^15,37–39^, resulting in the possibility of overlooking details that seem to be minor. It is likely that these details are those that are actually the crucial details for understanding the true nature and dynamics of the biofilm population. Direct evidence of factors from within the biofilm provides an important tool to tackle this challenge. The work presented here has focused on spatial and temporal localization of flagella in the biofilm, providing direct visualization for the first time.

A designated genetic code expansion system was developed for direct flagella labeling, enabling its direct identification and characterization in a mature biofilm. Following orthogonality establishment for the first time, this system was utilized for bio-tracking flagella inside biofilms. Direct imaging revealed inoculated flagella persistence during initial biofilm growth and maturation. It also revealed inoculated flagellar movement in the biofilm. Furthermore, as opposed to what was previously reported regarding flagella synthesis in a biofilm, we demonstrated that flagella are present in the biofilm throughout its entire lifecycle. While the presence of flagellated cells in compartments at the top of mushroom-like structures found in a grown biofilm are a common concept for dispersion of cells, we did not detect flagella any higher than the mid-section of these structures. Furthermore, we found that flagellated cells constantly left the biofilm, either carrying newly synthesized flagella or on rare occasions inoculated flagella. A combination of these findings led us to create an updated biofilm lifecycle model that includes the dynamics of flagella within these biofilms.

The unexpected discoveries regarding the presence of flagella inside the biofilm, led to a possible explanation for the reason behind this bacterial behavior with regards to the analysis of biofilm’s physical strength. The detection of scaffold-like structures of flagella inside the biofilm resembled mechanical support needed for architectural buildings. This was strengthened when native biofilm displayed higher physical strength compared to a biofilm lacking flagella. Accordingly, we concluded that flagella play an important role in providing mechanical and physical support of the biofilm. Further research though is needed to clarify the exact details behind this newly suggested role of flagella in a biofilm setting. We do recognize the existing obstacles at play, however, the work presented herein strongly emphasizes the need for additional direct evidence of occurrences within the biofilm environment. Direct imaging can thus serve as a window for new research venues. We posit that the new knowledge, afforded by this novel model, and approach that uses genetically code expanded strains as presented here will serve in tackling clinical/environmental biofilm research and envision multiple other new studies in regards to *P. aeruginosa*.

## Acknowledgments

Prof. Ehud Banin is greatly acknowledged for useful discussions. We would like to thank the Ilse Katz Institute for Nanoscale Science and Technology Shred Resource Facility for their technical contribution in image acquisition with Zeiss LSM880 Airyscan (Dr. Uzi Hadad), Verios 460L Thermo Fisher Scientific HR-SEM (Einat Nativ-Roth), Helios G4 UC Thermo Fisher Scientific dual-beam HR-SEM (Nitzan Maman) and MFP-3D-Bio AFM (Juergen Jopp). Dr. Anna Bakhrat’s assistance in genetic engineering is thankfully acknowledged. Yoni Ozer and Itay Algov’s graphical assistance is thankfully acknowledged. Esti Kramarsky Winter is acknowledged for writing assistance. We thank the Kreitmann School for graduate students for a Ph.D. fellowship (E.O., A.B., K.Y.) and the Ben-Gurion University for a continued support of our research (L.A., A.K.).

## Contributions

E.O. and K.Y. share equal contribution to this paper, E.O. conceived, performed and analyzed experimentations, established genetic code expansion system and authored the manuscript, K.Y. conceived, performed and analyzed all experimentations and co-authored the manuscript, E.C. performed nanoindentation experiments, A.B. synthesized unnatural amino acids used in this research, M.M.M., perceived and advised with P. Aeruginosa biofilm experiments, R.B. supervised, analyzed and wrote manuscript regarding AFM measurements, A.K. conceived experiments, supervised, provided research facilities and edited the manuscript, L.A. conceived experiments, supervised the research, provided facilities, written and edited the manuscript.

## Online Methods

### Reagents

PrK ((S)-2-Amino-6-((prop-2-ynyloxy)carbonylamino)hexanoic acid) and AzCK ((S)-2-Amino-6-((2-azidoethoxy)carbonylamino)hexanoic acid) were both synthesized according to a protocol reported by Nguyen et al.^48^. Tris(3-hydroxypropyltriazolylmethyl)amine (THPTA), Tetramethylrhodamine-azide and Tetramethylrhodamine-alkyne were purchased from Sigma-Aldrich (Rehovot, Israel). MB 660R Azide was a kind donation from Click Chemistry Tools (Scottsdale, AZ, USA). All restriction enzymes were purchased from Thermo Fisher Scientific (Waltham, MA, USA), while all DNA oligonucleotides were obtained from Syntezza Bioscience (Jerusalem, Israel).

### Plasmids construction

All plasmids for initial method establishment were constructed by a standard yeast assembly protocol^22^. The upstream and downstream regions from *P. aeruginosa*’s leucyl translational system were chosen for OTS expression. Upstream and downstream regions of leucyl tRNA “flanked” pyrrolysyl tRNA, as well as for the upstream and downstream regions of leucyl tRNA-synthetase that “flanked” pyrrolysyl tRNA-synthetase. The deGFP reporter gene (a variant designed for better expression in in-vitro systems, but also works well in in-vivo systems) was also chosen to have a PAO1 endogenous promoter, the flagellin native promoter (*fliC* promoter). It was chosen considering *fliC* gene was designed for Uaa incorporation after deGFP.

The initial construct (pPaGE Pyl TAG *fliC* prom deGFP WT NHis) was assembled in two stages. First, pMRP9-1 backbone was amplified using primers 1 and 2 (table S1), without the deGFP gene, and was assembled with the 2μ origin and URA3 selectivity gene for yeast. Second, the new vector after restriction with BamHI was assembled with seven other PCR amplified parts, containing the OTS, deGFP expression gene and necessary promoters and terminators (primers 3-16, table S1). Endogenous regions of *P. aeruginosa* were amplified from PAO1 genome. N-terminal his-tag deGFP and its T500 terminator were amplified from the pBEST plasmid^27^. Pyrrolysyl tRNA-synthetase was amplified from pEVOL-Pyl plasmid described in previous work^27^. Pyrrolysyl tRNA was assembled through primers 4 and 5 homology (table S1) during the yeast assembly. The pPaGE Pyl TAG *fliC* prom deGFP Y35TAG NHis construct was assembled in the same manner, only with a mutant deGFP amplified from the pBEST plasmid.

Variant for orthogonality testing of Pyl tRNA was generated through standard DNA collapse using HindIII. HindIII restriction, followed by self-ligation of the plasmid without the PylRS gene. During the initial construct generation, a deletion construct without Pyl tRNA was also created. This variant was used for orthogonality testing of PylRS.

A construct containing the *fliC* gene with a TAG mutation (pPaGE Pyl TAG *fliC* T248TAG) was assembled through Gibson assembly. pPaGE was restricted with NcoI for vector generation without the deGFP gene and T500 terminator. *fliC* gene had the TAG mutation installation as part of the assembly, by two pieces amplification. The gene, together with its downstream sequence, was amplified from the PAO1 genome using primers 17+20 and 19+18 (table S1).

All plasmids inserted into *E. coli* underwent standard heat-shock transformation protocol. All plasmids inserted into *P. aeruginosa* underwent standard electroporation protocol.

### Viability assay

Bacterial liquid culture in LB-Miller, after 24 hrs of growth at 37°C, was diluted 1:100. The diluted culture was placed at 37°C. Every 1 hr, a duplicate was measured for OD at a wavelength of 600 nm using a Synergy HT plate reader (Biotek, Winooski, VT, USA). After 17 hrs of measurements, when the cells reached growth plateau, the culture was left for incubation for another 10 hrs, when a final measurement was taken. When needed, the culture was supplemented with final concentration of 1 mM Uaa (optimal concentration found is shown in Fig S36). In case of plasmid containing bacteria, growth medium was supplemented with 300 μg/mL carbenicillin.

### Suppression efficiency

Five separate sets of liquid cultures were grown at 37°C for 24 hrs. Each set was tested for GFP expression and was composed of a native strain of *P. aeruginosa*, WT GFP and Y35TAG GFP with PrK. Following growth, each sample, from each set, was tested in triplicates for OD at a wavelength of 600 nm and GFP fluorescence by using a Synergy HT plate reader (Biotek, Winooski, VT, USA). Each sample’s fluorescence was divided by the OD_[600]_ value. Values of WT and Y35TAG with PrK were normalized to the native strain’s value of fluorescence/OD_[600]_. Finally, the suppression efficiency of each set was determined through value of normalized fluorescence\OD_[600]_ of mutant divided by the value of normalized fluorescence\OD_[600]_ of WT.

Error bar in figure 1e could only be calculated for mutant expression (as WT is always 100% by definition), representing the standard deviation between 5 different suppression efficiencies values calculated.

### Protein expression in PAO1 and cells lysis

Culture growth: PAO1 harboring pPaGE variants, were grown in LB-Miller with 300 μg/mL carbenicillin at 37°C for 24 hrs. After growth, the cultures were diluted 1:100 for another 24 hrs of growth. In case of needed Uaa addition, the culture was supplemented with final concentration of 1 mM Uaa.

Lysis: Following 24 hrs of growth, 1 mL of culture was sedimented and resuspended with 1 mL of phosphate buffer 100 mM pH 7. The cells were once again sedimented and were resuspended with 100 μL lysis solution composed of: 90% phosphate buffer, 10% BugBuster^®^ 10X protein extraction reagent (Merck, Billerica, MA, USA), Turbonuclease (Sigma, St. Louis, MO, USA), Lysozyme (Sigma-Aldrich, Rehovot, Israel) and protease inhibitor (Merck, Darmstadt, Germany). Cells were incubated with the lysis solution for 30 min at room temperature while shaking, followed by 4°C centrifugation at 10000 g for 10 min. Supernatant containing soluble proteins fraction was taken for further analysis.

### Click reaction followed by SDS-PAGE, fluorescent imaging and Western-blot analysis

Cu(I)-catalyzed azide-alkyne cycloaddition (CuAAC) click reaction^29^ was performed on lysates at a final volume of 50 μL. A fluorophore (with either an azide moiety or an alkyne moiety, according to necessity) was added to a concentration of 50 μM, while THPTA, sodium ascorbate and CuCl_2_ were added to final concentration of 1.2 mM, 2.5 mM and 200 μM, respectively. A volume of 20 μL of cell lysate was added to the reaction, followed by 1 hr incubation at room temperature with shaking. Clicked samples were examined through SDS-PAGE 4-20% Expressplus protein gel (GenScript, Nanjing, China). Fluorescent SDS-PAGE images were obtained through ImageQuant LAS 4000 imager (Fujifilm, Tokyo, Japan), using green light (520 nm Epi) and a 575 nm Cy3 detection filter. Next, when Western-blot was needed, SDS-PAGE was transferred to a membrane (Bio-Rad, Hercules, CA, USA) through eBlot^®^ protein transfer system (GenScript, Nanjing, China). Using goat T-19 anti GFP antibody^49^ (sc-5384, Santa Cruz, CA, USA) as a primary antibody and donkey anti goat IgG-HRP^50^ (sc-2020, Santa Cruz, CA, USA) as a secondary antibody, standard Western-blot protocol was performed. Chemiluminescence imaging was done using ImageQuant LAS 4000 imager (Fujifilm, Tokyo, Japan).

### Protein purification and mass spectrometry

A 200 mL culture of the P. aeruginosa GFP Y35TAG mutant, in the presence of PrK, was grown at 37°C for 24 hrs. Using standard needle sonication, culture was lysed and purified using IMAC (Novagene, Madison, WI, USA) according to manufacturer guidelines. Elution fraction was concentrated using Vivaspin 6, 10000 MWCO PES (Sartorius, Goettingen, Germany) and concentrated fraction was analyzed by liquid chromatography mass spectrometry (LCMS) (Finnigan Surveyor Autosampler Plus/LCQ Fleet, Thermo Scientific, Waltham, MA, USA).

### Theoretical model of P. aeruginosa flagella

A monomer model of *P. aeruginosa* flagellin was obtained through SWISS-MODEL^51^ and a PrK residue was incorporated in position 248 (PyMOL Molecular Graphics System, Version 2.0 Schrӧdinger, LLC). PDB file 5WK5, containing 41 subunits of the *P. aeruginosa*’s filament core (no outer protein structure of D2 and/or D3), was used as alignment reference for 41 SWISS-MODEL generated monomers with PrK (PyMOL Molecular Graphics System, Version 2.0 Schrӧdinger, LLC). The final product is a theoretical model of a flagella filament, consisting of 41 *P. aeruginosa* flagellin proteins, all of which contain PrK in the 248^th^ position.

### Live-cell click reaction and flow-cells construction

pPaGE harboring *fliC* with TAG mutation at the 248^th^ site, was electroporated into PAO1 gGFP strain (a strain containing GFP expression gene in the genome, gGFP = genome GFP). Liquid culture in the presence of PrK, was grown at 37°C for 24 hrs. Grown culture at the volume of 200 μL was centrifuged and resuspended with 20 μL phosphate buffer 100 mM pH 7. Click reagents were added to a final reaction volume of 50 μL. Azide-containing 545 fluorophore was added to a concentration of 50 μM, while THPTA, sodium ascorbate and CuCl_2_ were added to a final concentration of 1.2 mM, 2.5 mM and 200 μM, respectively. Following 40 min incubation at room temperature with shaking, cells were washed three times with 1 mL of phosphate buffer 100 mM pH 7 and were brought to a value of OD_600_ of 0.1. A flow-cell system was constructed as described before^52^, using the labeled bacteria as inoculation for growth. Following 1 hr of static attachment in the flow-cell at 30°C, AB minimal growth media at temperature of 37°C was supplied at a rate of 4 mL/hr. Flow-cells were grown for up to 6 days while imaging for different experimental procedures was done using confocal laser scanning microscopy (CLSM).

For pre-inoculation labeled flagella tracking –flow-cell was constructed as described and was imaged using CLSM at 2 different channels (488 at cyan for bacteria and 545 at magenta for pre-inoculation labeled flagella) at specific locations every several hours for up to 6 days.

For effluent flagella tracking –flow-cell was constructed as described while AB minimal media was supplemented with PrK for continuous incorporation. Effluent of ~15 mL was collected into ice following 2/4/6 days of growth. Effluent total volume was pelleted down and resuspended with 20 μL phosphate buffer 100 mM pH 7. Click reagents were added to a final reaction volume of 50 μL. Azide-containing 660 fluorophore was added to a concentration of 50 μM, while THPTA, sodium ascorbate and CuCl_2_ were added to a final concentration of 1.2 mM,

2.5 mM and 200 μM, respectively. Following 40 min incubation at room temperature with shaking, cells were washed two times with 1 mL of phosphate buffer 100 mM pH 7 and were finally resuspended with ~50 μL PB 100 mM pH 7. Labeled effluent (5 μL onto a glass slide) was imaged using CLSM at 3 different channels (488 at cyan for bacteria, 545 at magenta for pre-inoculation labeled flagella, 660 at orange for newly synthesized labeled flagella).

For newly synthesized labeled flagella tracking within a biofilm –flow-cell was constructed as described and grown for 2/4/6 days while AB minimal media was supplemented with PrK for continuous incorporation. Click reagents solution was prepared for a final reaction volume of 500 μL. Azide-containing 545 fluorophore was added to a concentration of 50 μM, while THPTA, sodium ascorbate and CuCl_2_ were added to a final concentration of 1.2 mM, 2.5 mM and 200 μM, respectively. The click mixture was slowly injected into the flow-cell chamber and remained stationary for 20 min at room temperature, followed by intensive wash with growth media at a rate of 16 mL/hr at room temperature. Flow-cell was imaged using CLSM at 2 different channels (488 at cyan for bacteria, 545 at magenta for labeled flagella).

All mentioned experimental procedures were repeated for a minimum number of 5 different flow-cells.

### Confocal microscope settings and image analysis

Confocal images were acquired using Zeiss LSM880 system (Zena, Germany). Plan-Apochromat 63x/1.4 Oil DIC M27 or Plan-Apochromat 40x/1.3 Oil DIC M27 objective were used. Cyan channel was imaged using 488 nm Argon laser (usually in the range of 5-15%) with emission filter BP 493-556. Magenta channel was imaged using 561 nm DPSS laser (usually in the range of 2-4%) with emission filter BP 570-624. Orange channel was imaged using 633 nm HeNe laser (usually in the range of 8-15%) with emission filter BP 638-755.

Scanning resolution for all flow-cells was 1024×1024. Scanning resolution for effluent planktonic cells was 2048×2048.

Image analysis was performed using either ImageJ software (National institutes of health, USA) or IMARIS software (Bitplane AG, Zurich, Switzerland). Despite channel modifications in image analysis, a gamma value of 1.00 was strictly conserved in all images and analysis.

Figure 2b –all 9 Z plains were stacked and channels were merged (488+545) and a smooth filter was used. (ImageJ)

Figures 2d – 3D digital visualization of 545 alone or merged channels (488+545) were at 6000×6000 dimensions and 900 dpi using “shadow” function. (IMARIS)

Figure 2e – Out of 118 Z plains, 61-65 were stacked and 488+545 channels were merged. (ImageJ)

Figures 3b – Digital bio-volume was saved as 6000×6000 and 900 dpi. 2 days mid-section was stacked from 13-16 Z plains. Upper-section was stacked from 20-25 Z plains. Lower mid-section was stacked from 27-39 Z plains. Mid-section was stacked from 66-68 Z plains. (IMARIS)

Figures 4b – Out of 13 Z plains, image was stacked using 4-9 Z plains and saved as individual channels (488/545/660) or a merged image of all channels. (ImageJ)

Figure 6a – The 14 Z plain was presented only in the 545 channel. (IMARIS)

Figure 6b – Z plains of 23-26 were stacked and saved as individual 545 channel or a merged image of 488+545. (IMARIS)

### P. aeruginosa PAO1 gGFP ΔfliC strain generation

For the creation of a KO strain to flagella filament, a CRISPR/Cas9-based platform adapted to *P. aeruginosa*^43^ was utilized. Out of a 2-plasmids system, pCasPA plasmid was used as is, while a relevant pACRISPR plasmid had to be constructed using Gibson assembly to contain gRNA and complementary homology region to the genome. The gRNA used was chosen based on CHOPCHOP^53^ and complemented sites 168-174 in *fliC* gene. Homology region was chosen as 500 bp upstream to the ATG codon of *fliC*, 30 bp from the end of the *fliC* gene (including stop-codon) and 500 bp downstream to the stop-codon of *fliC*, meaning deletion of 1437 bp out of the *fliC* gene. Gibson assembly for pACRISPR with relevant gRNA and homology was done using primers 21-28 (table S1), following standard assembly and plasmid sequence verification. pCasPA was electroporated into PAO1 gGFP strain using standard protocol and grown on tetracycline 100 μg/mL selective plate. After first plasmid insertion, PAO1 gGFP pCasPA was grown over night at 37°C, and was added with L-arabinose to final concentration of 2 mg/mL for 2 hrs incubation at 37°C (targeted for Cas9 inductive protein expression). Cells then were prepared for electroporation using standard protocol. pACRISPR targeted for *fliC* deletion was electroporated into PAO1 gGFP pCasPA cells prepared in advance and grown on tetracycline 100 μg/mL + carbenicillin 150 μg/mL. Colonies were screened for genome segment modification and chosen colonies were cured from both plasmids by plating on sucrose 5% plates. Chosen colony was isolated, tested for positive plasmid curing, and tested by PCR for positive deletion (primers 29+30, table S1). PCR segment was purified using nucleospin gel and PCR clean-up (MACHEREY-NAGEL, Germany) and underwent Sanger sequencing with primer 31 (table S1).

### WT/KO strains flow-cells construction

WT (PAO1 gGFP) or KO (PAO1 gGFP Δ*fliC*) strain stationary grown liquid culture was diluted to OD_600_ of 0.1 and cells were inoculated in a flow-cell as described before. After 1 hr of attachment at 30°C, 37°C heated AB minimal media at a rate of 4 mL/hr was supplemented for 5 days of growth. Flow-cells were examined for biofilm structure and 3D maturation using CLSM.

### Transmission electron microscopy (TEM)

Grown bacteria cultures were diluted 1:20. A carbon type-B TEM grid was prepared using plasma cleaner PDC-32G (HARRICK PLASMA, Ithaca, NY, USA) for 30 sec. TEM grids were loaded with 2 μL from the diluted liquid culture, while excess liquid was dried using filter paper.

Prepared grids were examined in FEI Tecnai T12 G2 TWIN transmission electron microscope operating at 120 kV.

### Swarming assay

*P. aeruginosa* PA01 were grown overnight in LB-Miller medium. The next day, the bacteria were diluted 1:100 into fresh M9 medium and were grown to mid-log phase. The swarming plates were prepared using M9, solidified with 0.7% [wt/vol] Difco agar. Following bacteria refreshment, 2 μL of the inoculums were placed in the middle of the plates enabling assessment of surface coverage after 24 hrs of growth at 37°C.

### Static biofilm growth

Round glass coverslips, 15 mm diameter (Ted Pella, Inc., Redding, CA, USA) were incubated with HCl 8% for 1 hr at room temperature and underwent autoclave sterilization in advance to biofilm growth. WT (PAO1 gGFP) or KO (PAO1 gGFP Δ*fliC*) were refreshed by taking 30 μL stationary grown culture into 3 mL LB-Miller for 3 hrs at 37°C with agitation. Reaching an OD_600_ of ~0.2, 20 μL of refreshed culture were taken into 2 mL of LB-Lenox in a 12-well plate’s well. Treated glass cover slip was placed in the well as well. Plate was incubated at 37°C for 16 hrs. Wells surrounding sample, were filled with DDW to keep high moisture during growth.

### High-resolution scanning electron microscopy (HR-SEM)

Following static biofilm growth, liquid was aspirated and the glass cover slip underwent fixation (2.5% glutaraldehyde, 2% paraformaldehyde, 0.2 M phosphate buffer) for 15 min. Samples were washed twice with PBS, 10 min each time, and were dehydrated using increasing ethanol concentrations before an addition of hexamethyldisilazane/ethanol solution in increasing concentrations. Finally, all liquids were removed and samples were dried in a chemical hood.

Fixed glasses were coated in Cr using Quorum Q150T-ES sputter and examined in Verios 460L Thermo Fisher Scientific scanning electron microscope operating at 3.00 kV.

### Dual-beam HR-SEM

For an inside look at a biofilm and not just outer-rim observations, the same glasses coated with Cr that were used for HR-SEM, were also analyzed with dual-beam HR-SEM. Areas meant for examination were locally coated with Pt and were sliced using focused ion beam. Biofilm’s inside was examined in Helios G4 UC Thermo Fisher Scientific scanning electron microscope operating at 5.00 kV.

### Atomic force microscopy (AFM) imaging

Following static biofilm growth, the well was washed twice with PBS and left to dry in a hood. The dry samples were imaged on a MFP-3D-Bio (Oxford Instruments Asylum Research, Santa Barbara, CA, USA) in AC-mode ("tapping mode") in air, using an AC240BSA probe (Olympus) at room temperature. Imaging rate was of 0.5 Hz, with parameters of 512 scan lines and 1024 scan points.

### AFM nanoindentation

Local stiffness modulus, *K*, of the two different biofilms was measured with nano-indentation experiments performed on a Luigs & Neurann LTD AFM. The biofilm samples were grown on a clean glass coverslip as was mentioned before. The grown biofilm on top the coverslip was gently washed with PBS and semi-dried in open air. The indentation was performed with pyramidal silicon nitride cantilevers (V-shape MLCT, Bruker) with a measured mean spring constant of 0.01 N/m. The normal spring constants of the cantilevers were determined before each measurement using the equipartition theorem^54^. In a nano-indentation measurement, force-distance curves are collected by approaching the cantilever tip towards the surface of the biofilm sample at a rate of 400 nm/s and indentation depth amplitude of 500 nm. Corresponding to the compliance of the sample, the cantilever deflects proportionally to the compliance of the sample. The curvature of this response region was fitted using the Sneddon model^55,56^, a contact mechanics model for a conical sharp probe (as the one used), to the force distance curves:

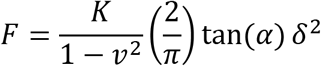

where *v* is Poisson ratio, taken as 0.5 (typical value for incompressible materials), *δ* is the indentation length coordinate, *θ* is the cone half angle (face angle) of the AFM probe, taken as the manufacturer's nominal value of 29.1°, and *K* is the local stiffness modulus. The indentation was performed at random locations across the surface of each biofilm sample. The measurement sets were repeated three times for each biofilm sample, where each set was comprised of 100 force-distance traces. All measurements were carried out under PBS buffer (150 mM NaCl, 20 mM PBS, pH 7.2) at room temperature. All data were recorded and analyzed using custom software written in Igor Pro 6.37 (Wavematrics).

From 100 collected values of *K* for each biofilm, we constructed their probability distributions, and fitted them with the relevant statistical model. The Freedman-Diaconis rule was used as criterion for setting the bin size of the distributions^57^. The stiffness distribution of the WT biofilm was fitted with a Gaussian distribution, *ϕ*(*K*) = (*σ*(2*π*)^½^)^−1^exp{–½[(*K* – *μ*)/*σ*]^2^}, and the stiffness distribution of the Δ*fliC* biofilm was fitted with the Gamma distribution, which is given by *ϕ*(*K*) = (*β*^*α*^/Γ(*α*))*K*^*α*−1^e^*−βK*^. This latter is highly useful to describe the distribution of a random variable that does not normally distribute. The mean is calculated by *μ* and *α*/*β* (for the normal and Gamma distributions respectively) and the variance (from which the standard deviation is calculated by taking its root) by *σ*^2^ and *α*/*β*^2^ (for the normal and Gamma distributions respectively).

