## supplementary information for "An inside look at a biofilm: Pseudomonas aeruginosa flagella bio-tracking"

**Table of Contents:**


| | | |
|---|---|---|
| Cover page | | S-1 |
| Table of contents | | S-2 |
| Figure S1 | Viability assay | S-3 |
| Figure S2 | ESI mass spectrometry | S-4 |
| Figure S3 | Flagellin Uaa incorporation | S-5 |
| Figure S4 | Planktonic bacteria TEM imaging with PrK incorporated flagellum | S-6 |
| Figure S5 | Click reaction in the absence of Uaa | S-7 |
| Figure S6-S11 | Figure 2 enlarged images | S-8 |
| Figures S12-S26 | Figure 3 enlarged images | S-14 |
| Figure S27-S38 | Figure 4 enlarged images | S-29 |
| Figure S39-S40 | Figure 6 enlarged images | S-41 |
| Figure S41 | Δ*fliC* validation for lacking flagella | S-43 |
| Figure S42 | Biofilm structure comparison through CLSM | S-44 |
| Figure S43 | Biofilm structure comparison through HR-SEM | S-45 |
| Figure S44 | Biofilm comparison through AFM | S-46 |
| Figure S45 | Uaa concentration determination | S-47 |
| Gene sequences | | S-48 |
| Table S1 | Primers list | S-54 |

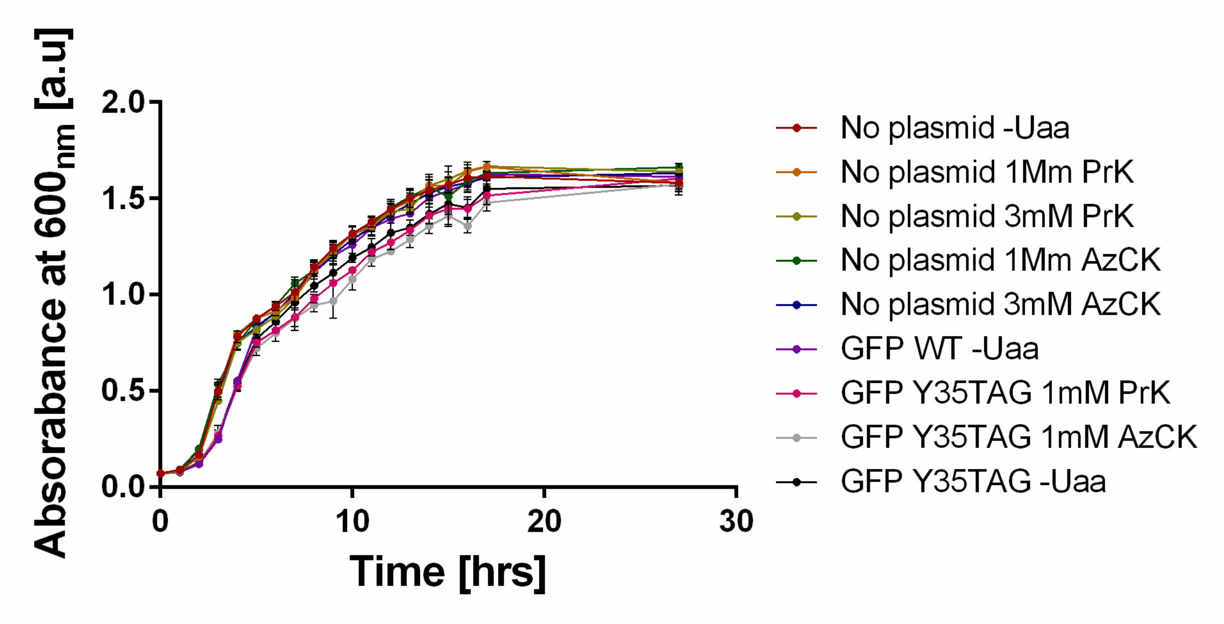


**Figure S1| Viability assay in response to the pPaGE plasmid and Uaas used in this research.** Hourly measurements of culture turbidity over a growth period of 17 hours, followed by a final measurement after 27 hours. The pPaGE plasmid and Uaas in media were tested for possible effect on *P. aeruginosa* growth.

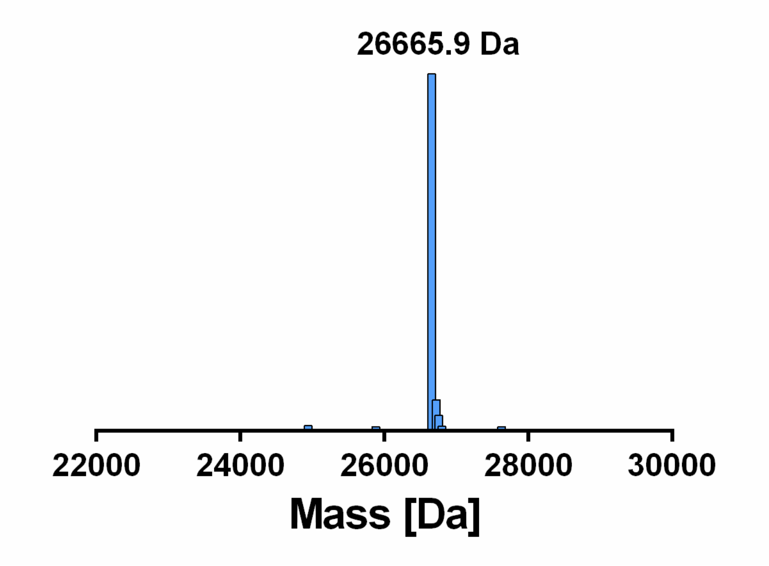


**Figure S2| ESI mass spectrometry identification of deGFP incorporating PrK.** Deconvoluted mass of 26665.9 Da for the deGFP Y35PrK purified protein (calculated mass of 26667.76 Da well within machine error).

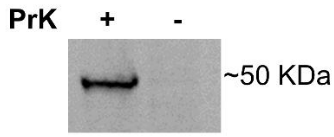


**Figure S3| Uaa incorporation into flagellin.** Fluorescent SDS-PAGE examining PrK incorporation into flagellin protein. Samples were lysed and insoluble proteins fraction underwent CuAAC click reaction to an azide-containing fluorescent dye.

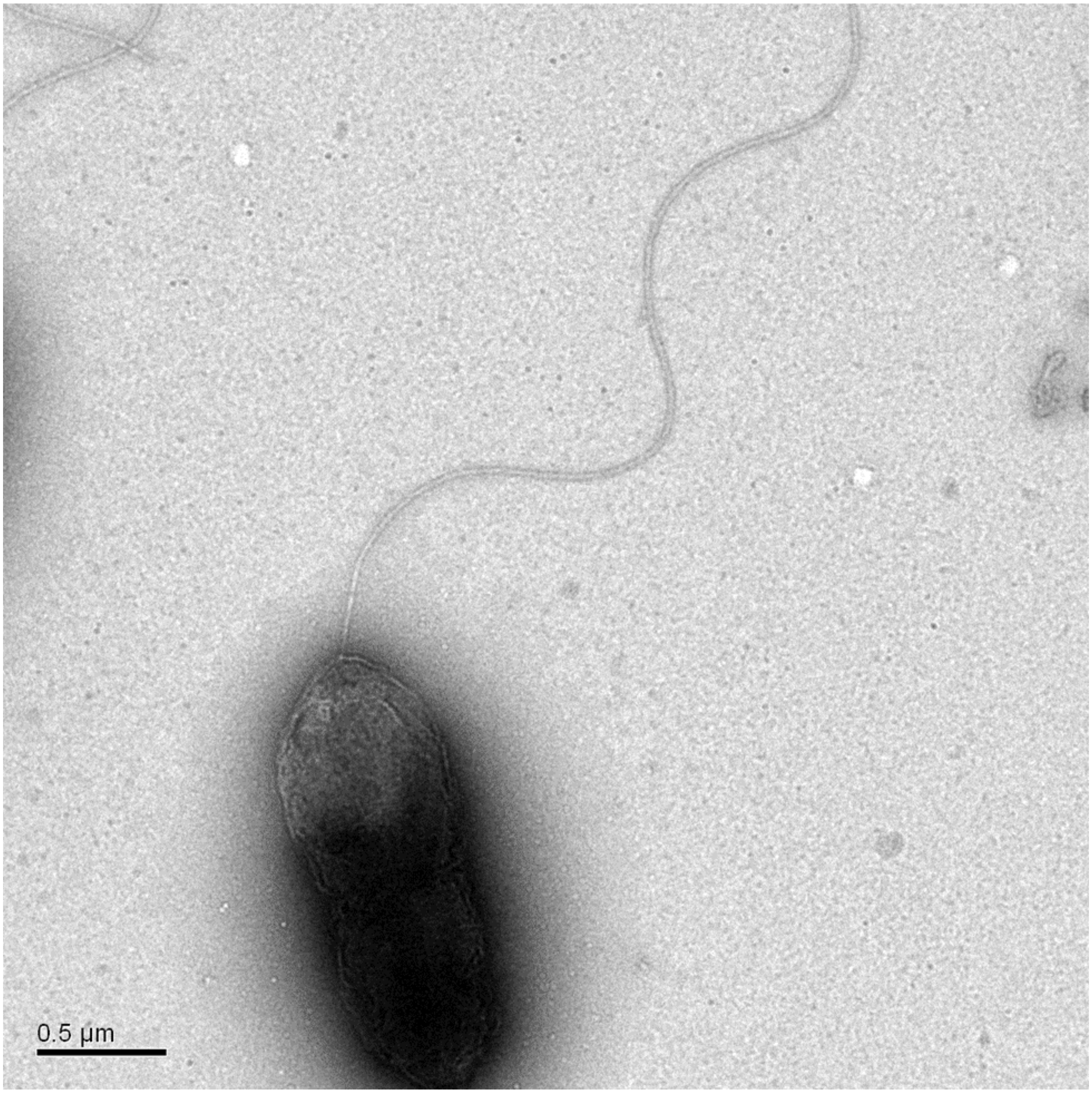


**Figure S4| Flagellin incorporated with PrK does not affect bacterial morphology.** TEM imaging of planktonic bacteria incorporating PrK into flagellin and possessing a single flagellum.

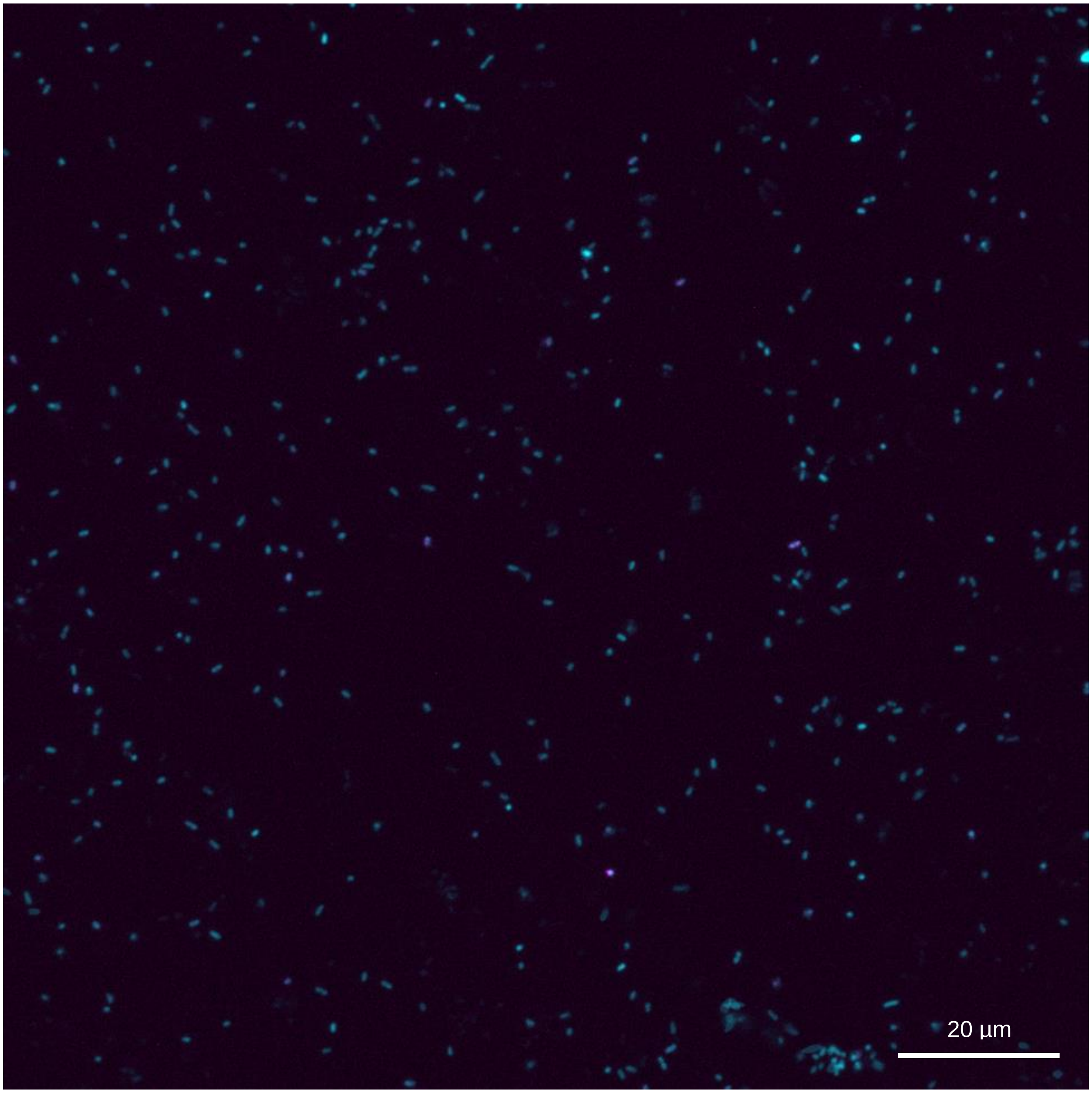


**Figure S5| Bacterial imaging in the absence of a Uaa.** CLSM of Planktonic cells expressing genomic GFP, in the absence of a Uaa. Sample was labeled with an azide-containing 545 nm fluorophore through CuAAC click reaction. Channels: λ=488 nm and λ=545 nm.

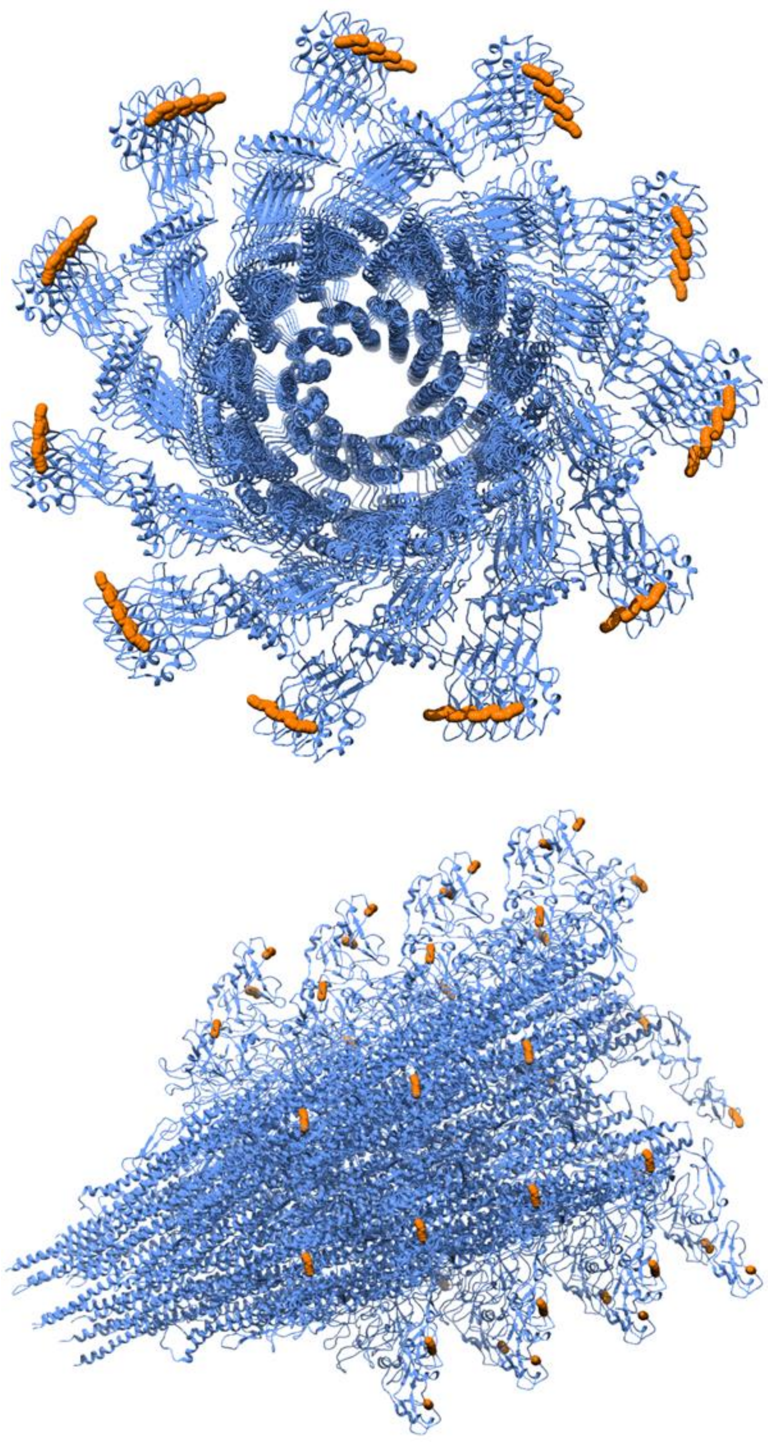

**Figure S6| Theoretical model of flagella incorporated with PrK.** Enlarged images of theoretical model generated.

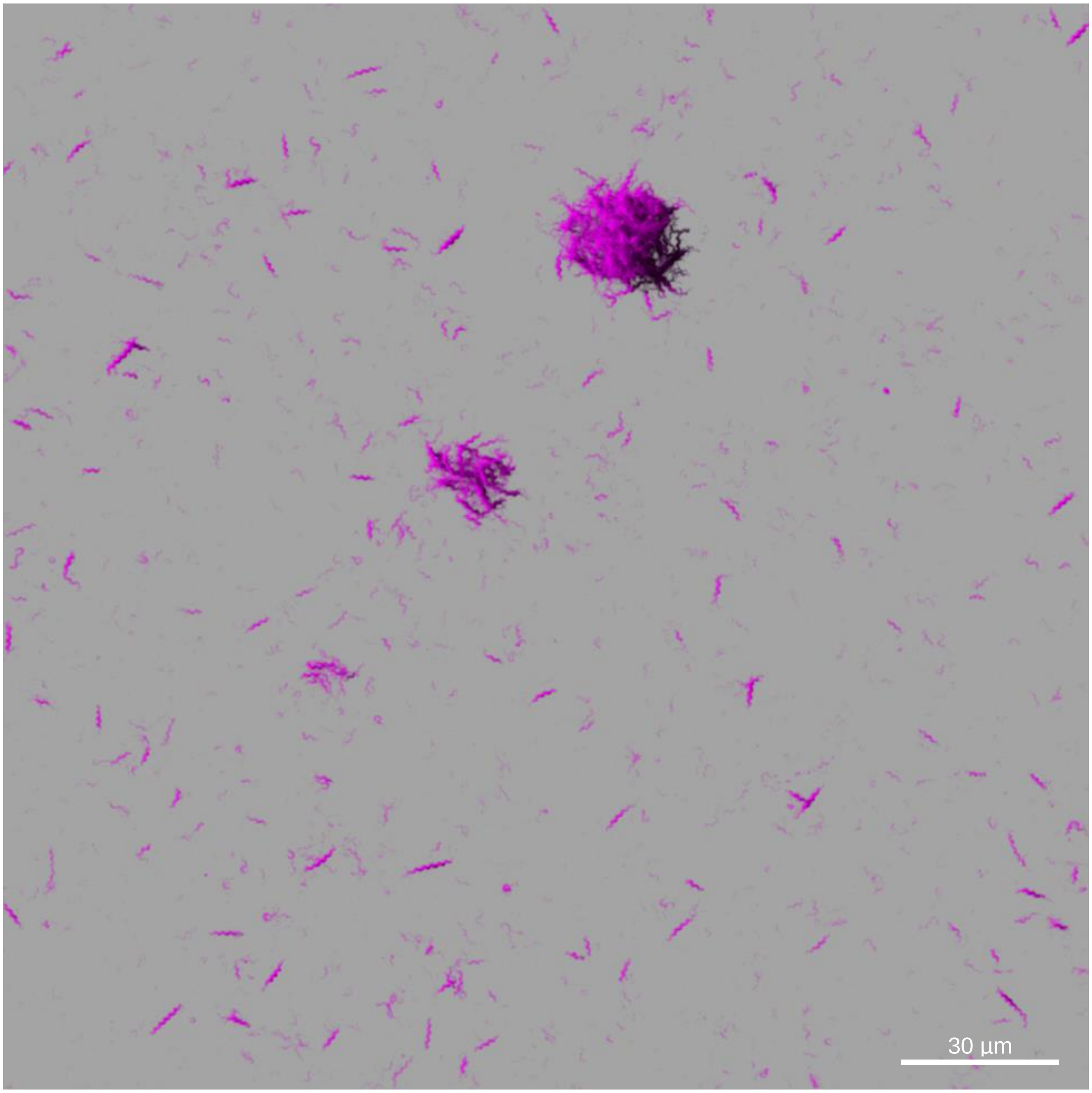


**Figure S7| Inoculated flagella visualization in a young biofilm.** Enlarged image of pre-labeled flagella in a young biofilm expressing genomic GFP (8 hrs). Channel: λ=545 nm.

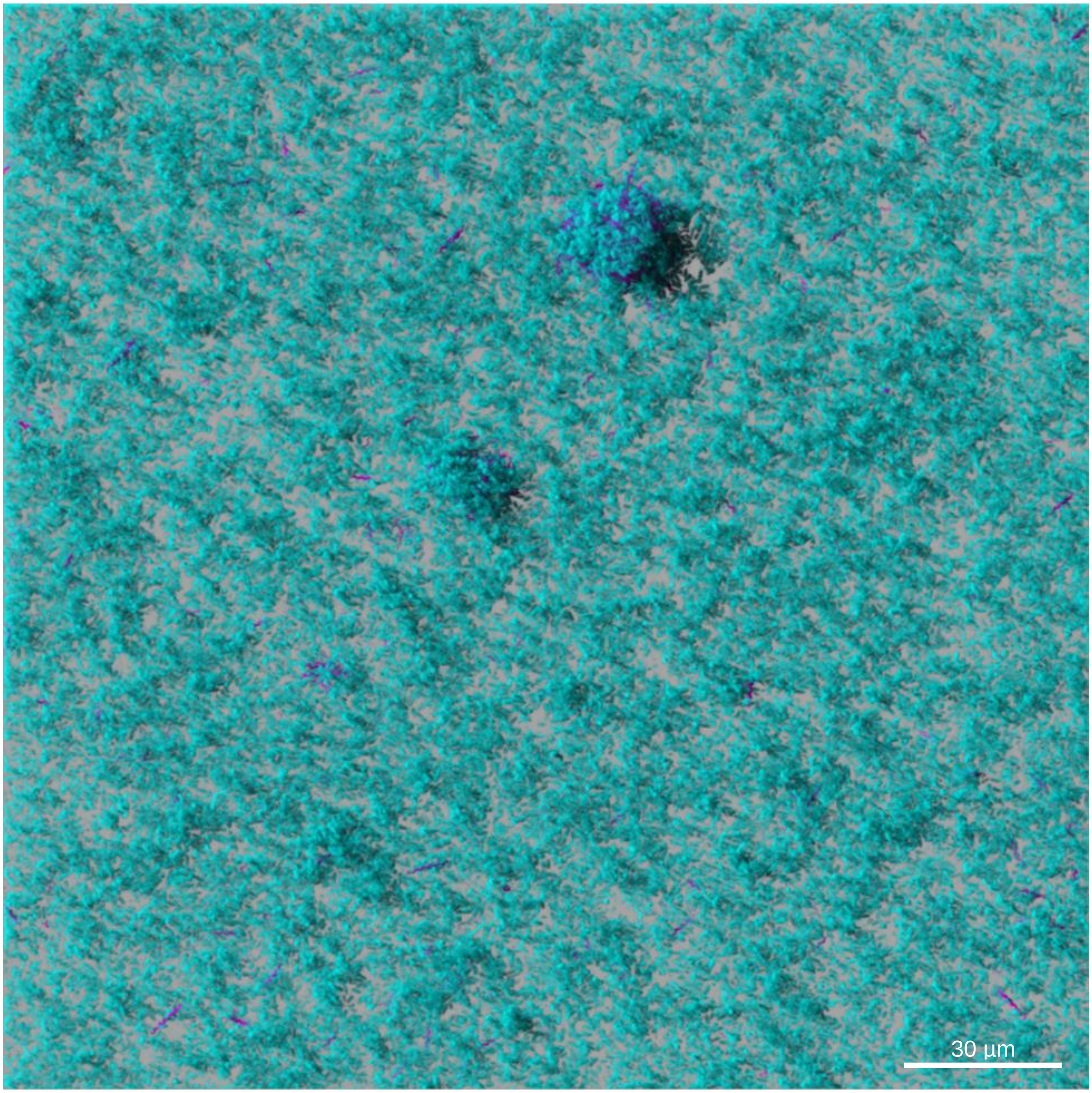


**Figure S8| Inoculated flagella and bacteria visualization in a young biofilm.** Enlarged image of pre-labeled flagella in a young biofilm expressing genomic GFP (8 hrs). Channels: λ=488 nm, λ=545 nm.

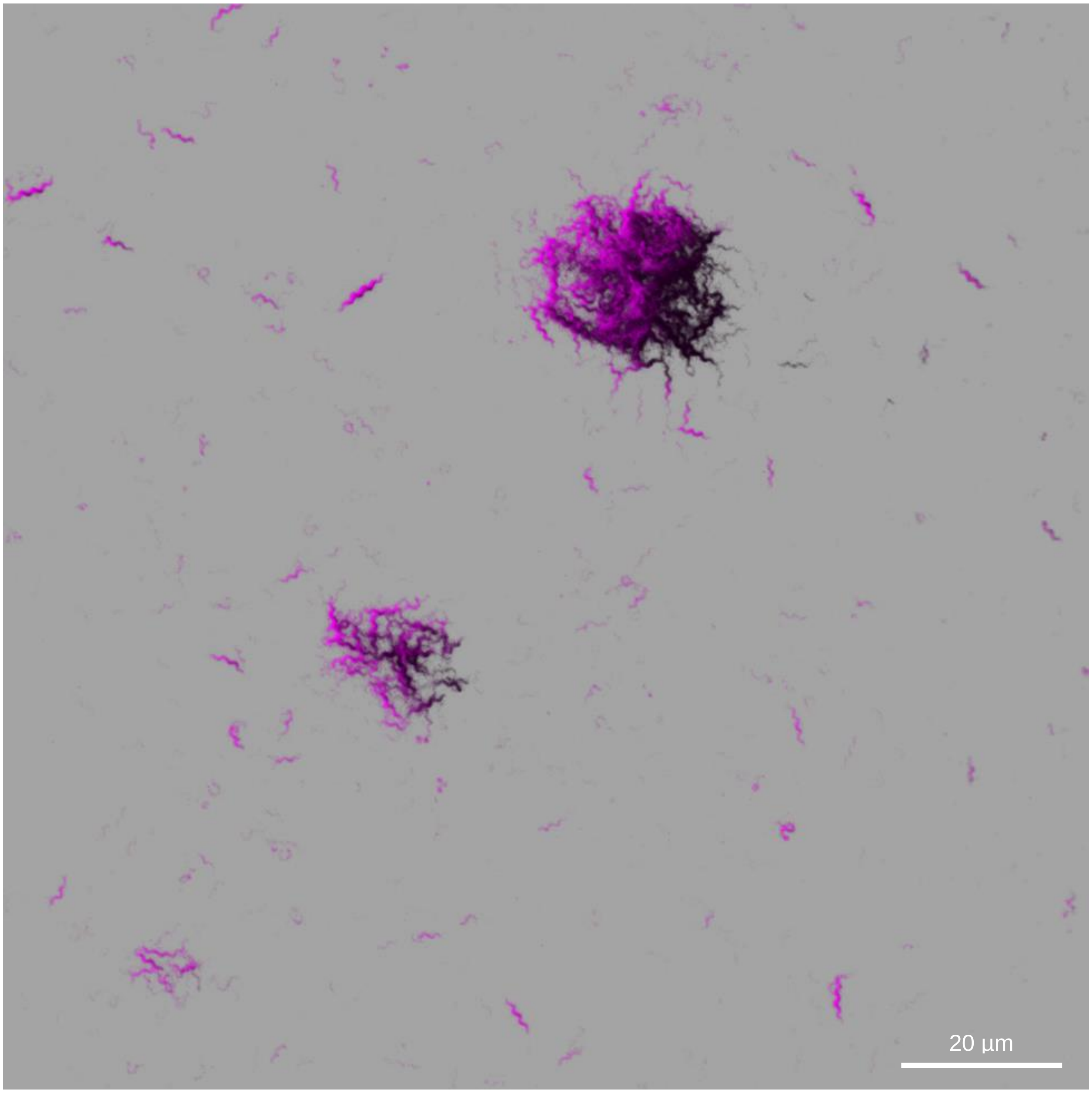


**Figure S9| Inoculated flagella visualization in a grown biofilm.** Enlarged image of pre-labeled flagella in a grown biofilm expressing genomic GFP (6 days). Channel: λ=545 nm.

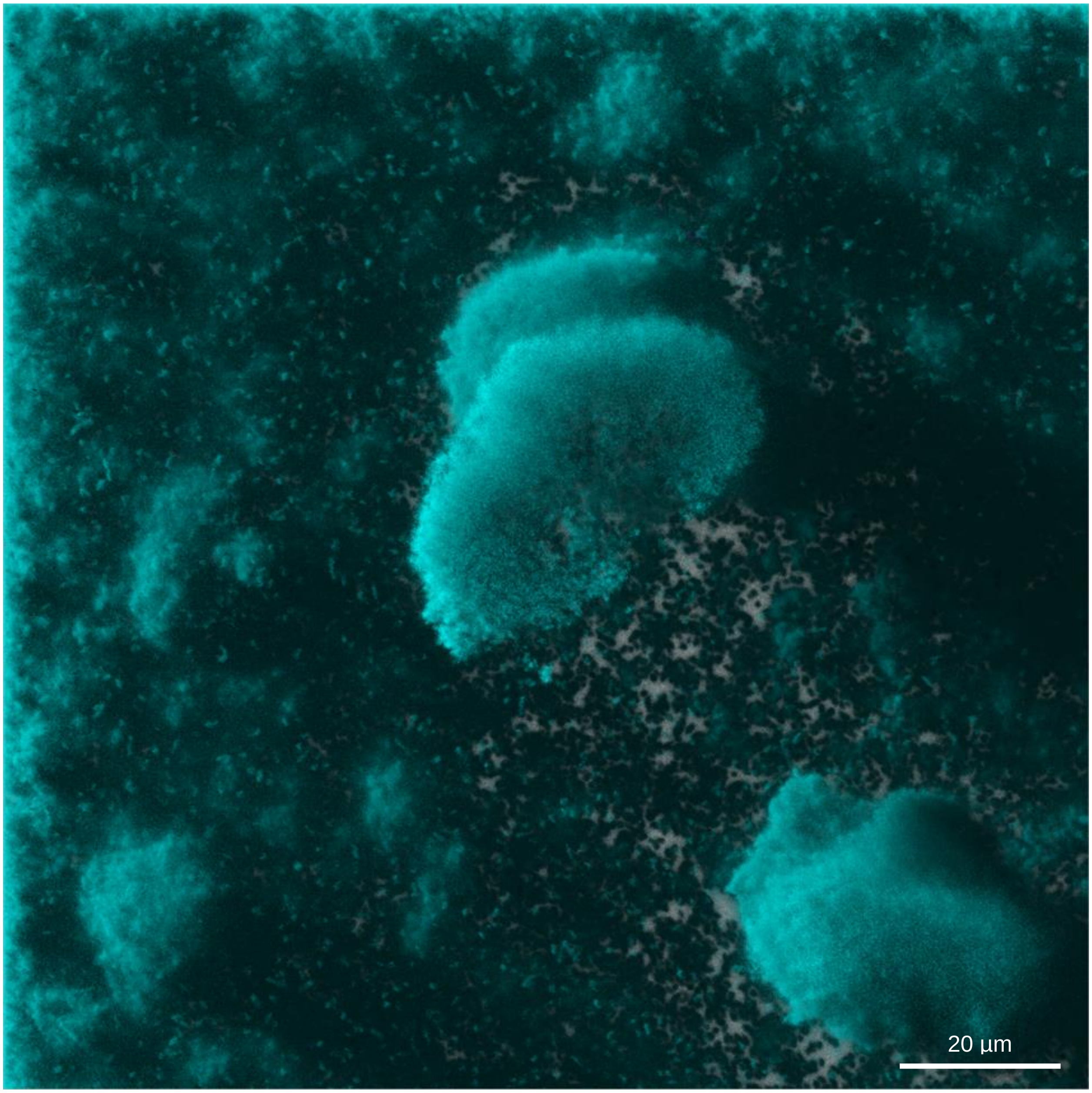


**Figure S10| Inoculated flagella and bacteria visualization in a grown biofilm.** Enlarged image of pre-labeled flagella in a grown biofilm expressing genomic GFP (6 days). Channels: λ=488 nm, λ=545 nm.

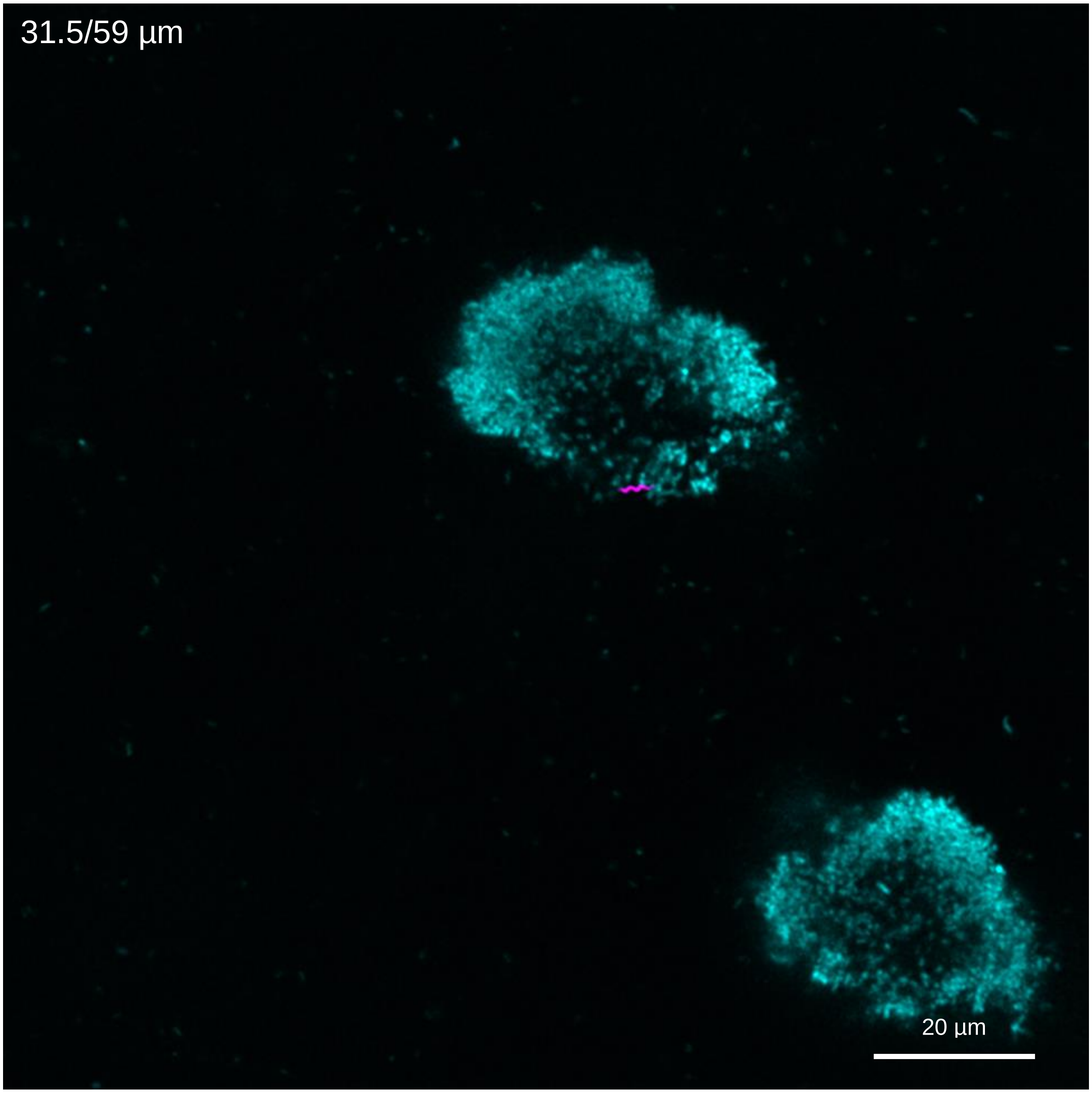


**Figure S11| Inoculated flagella movement during biofilm growth.** Enlarged image of pre-labeled flagella in a grown biofilm (6 days) expressing genomic GFP. Channels: λ=488 nm and λ=545 nm.

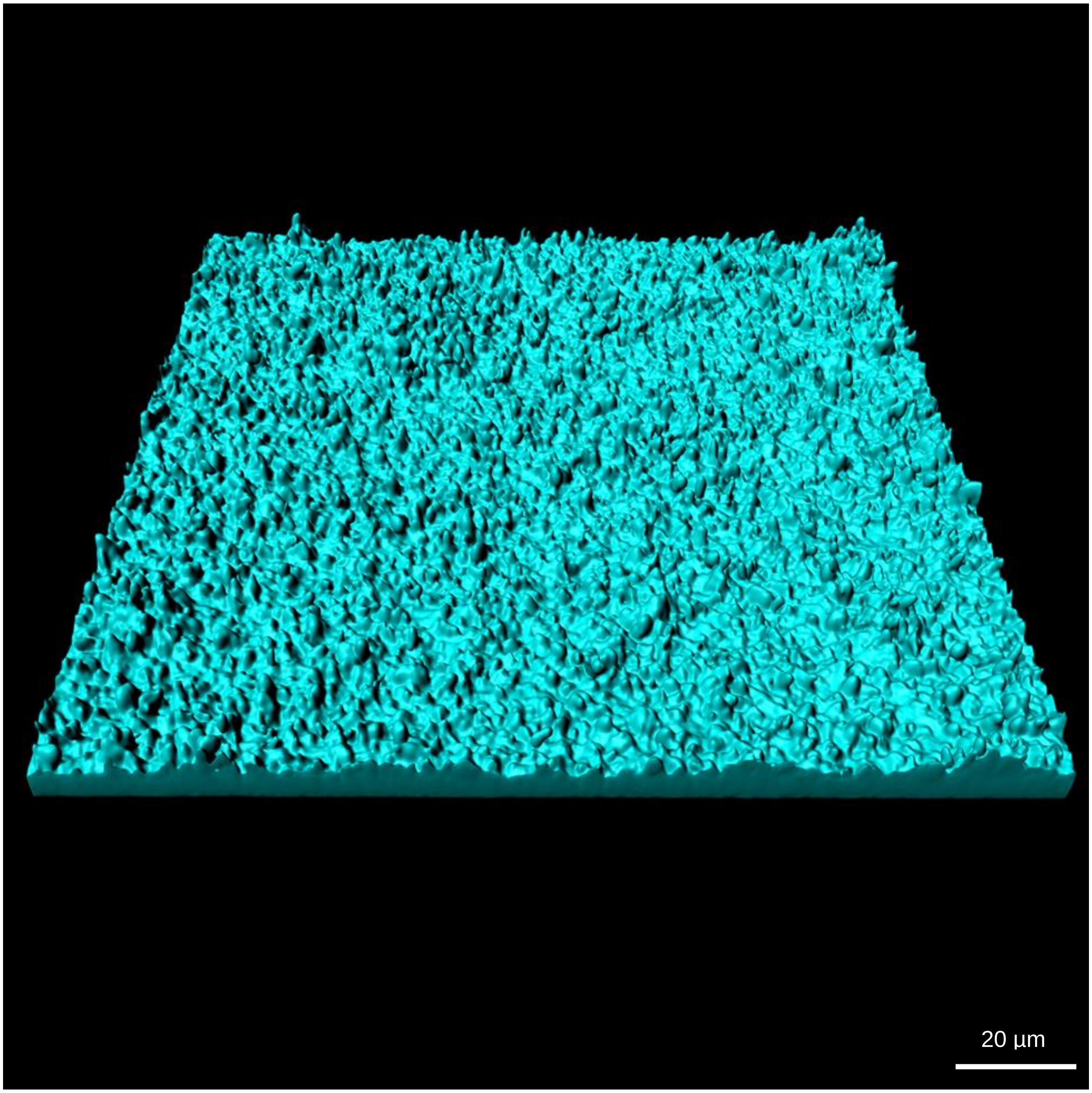


**Figure S12| Young biofilm biomass.** Enlarged image of visualized biofilm entire biomass after 2 days of growth.

Channel: λ=488 nm

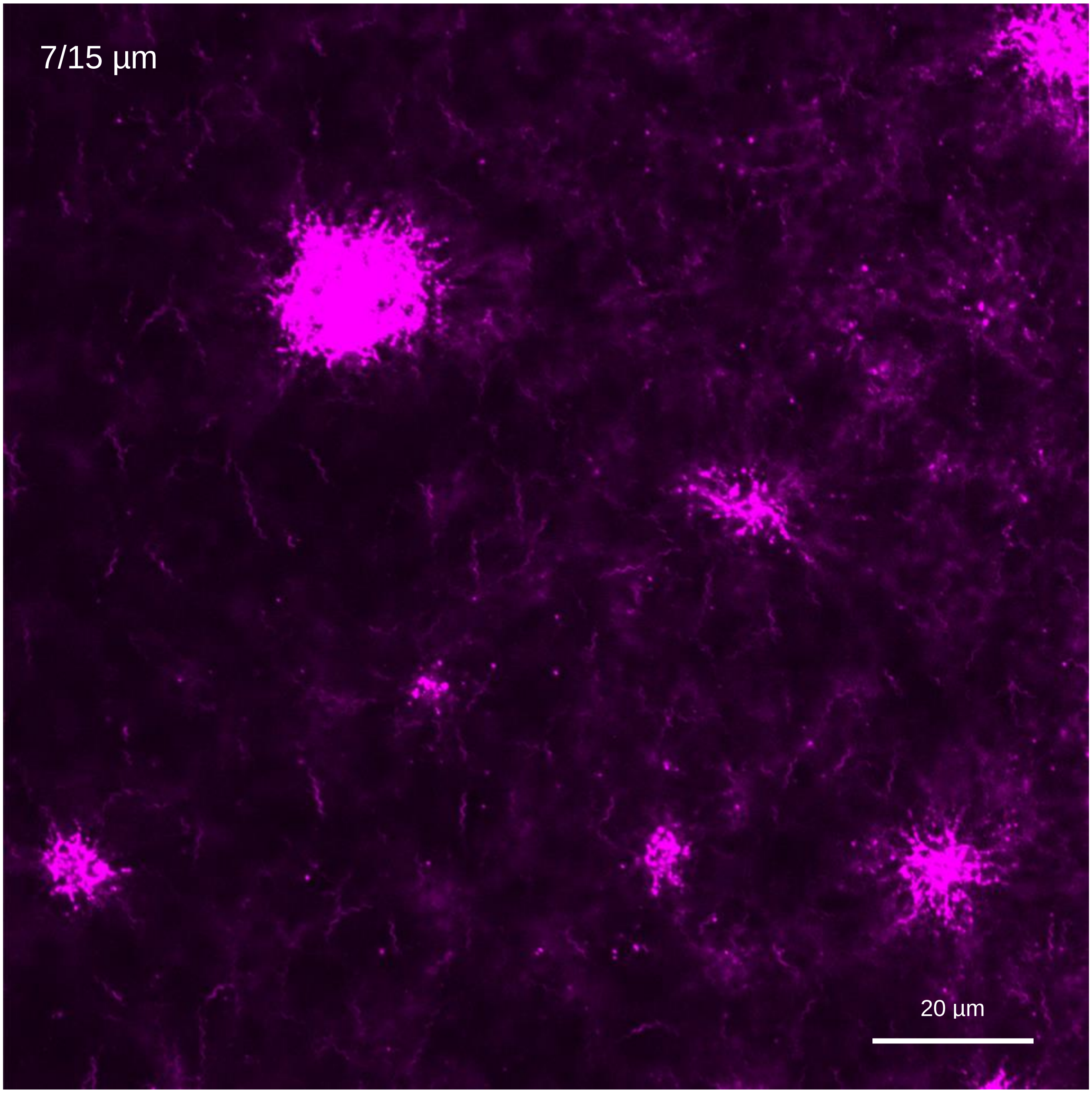


**Figure S13| Young biofilm flagella in mid-section.** Enlarged image of stacked layers from a young biofilm mid-section after 2 days of growth presenting labeled flagella. Channel: λ=545 nm

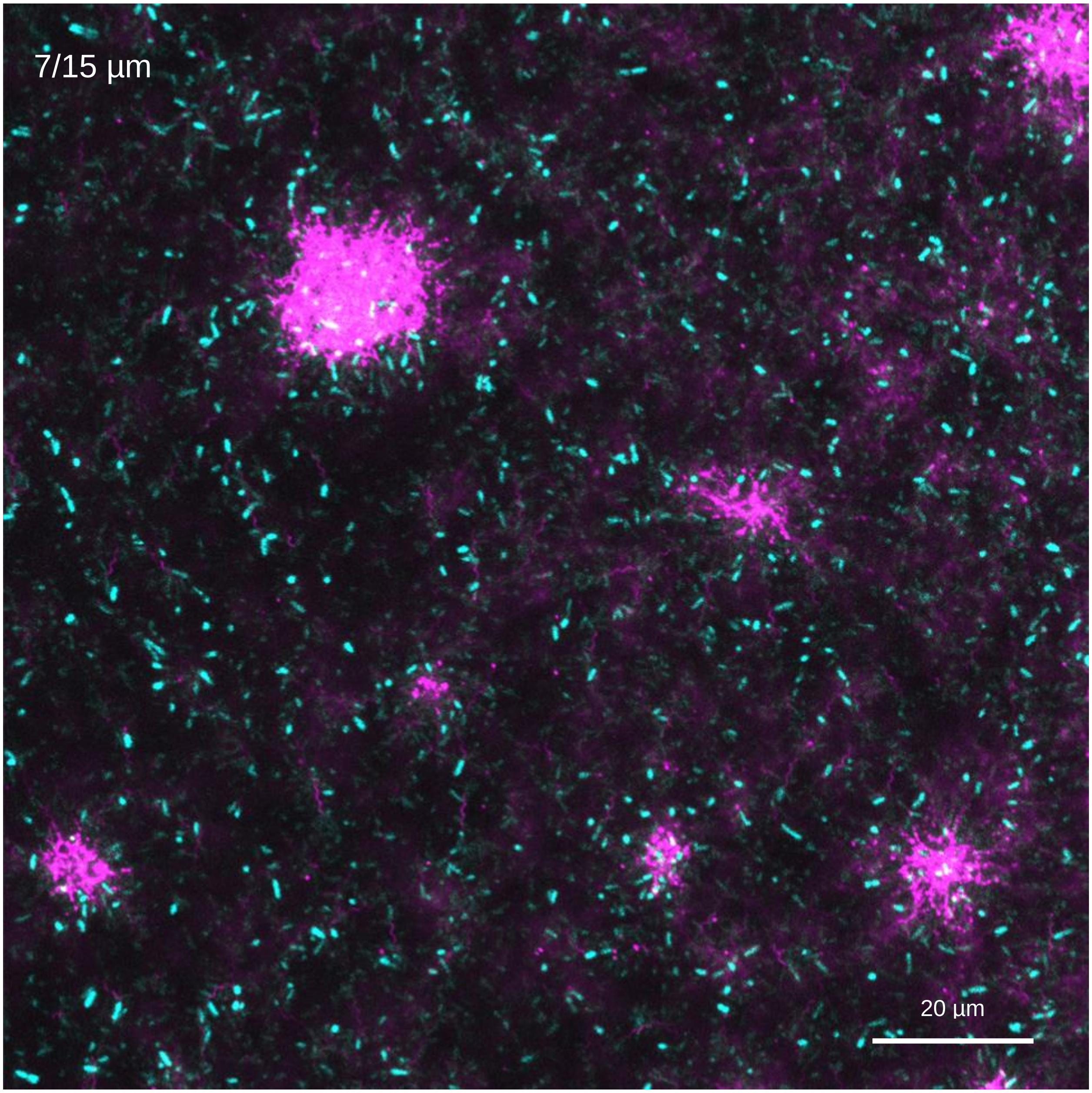


**Figure S14| Young biofilm flagella in mid-section.** Enlarged image of stacked layers from a young biofilm mid-section after 2 days of growth presenting labeled flagella and bacteria. Channels: λ=488 nm, λ=545 nm

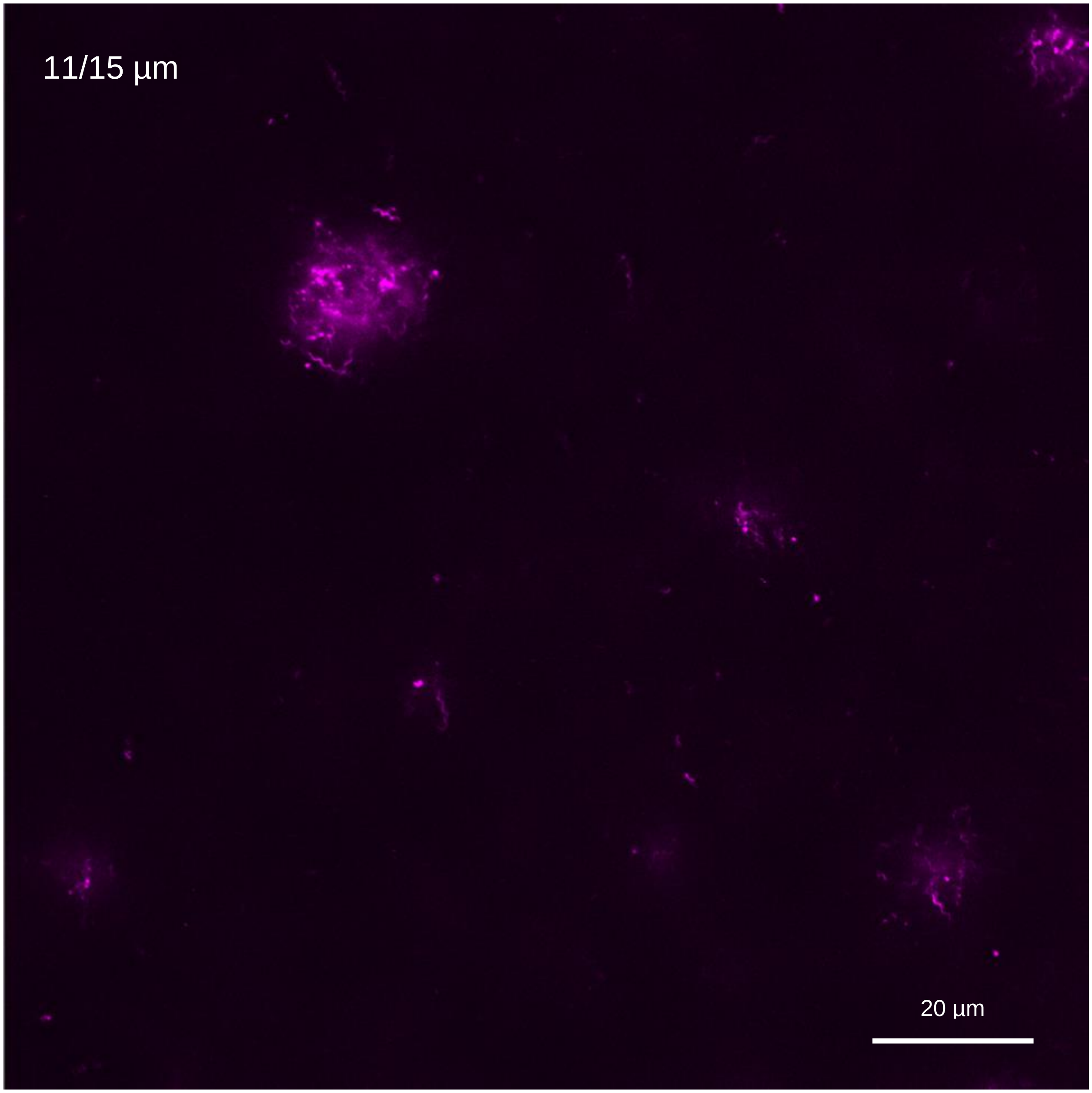


**Figure S15| Young biofilm flagella in upper section.** Enlarged image of stacked layers from a young biofilm upper section after 2 days of growth presenting labeled flagella. Channel: λ=545 nm

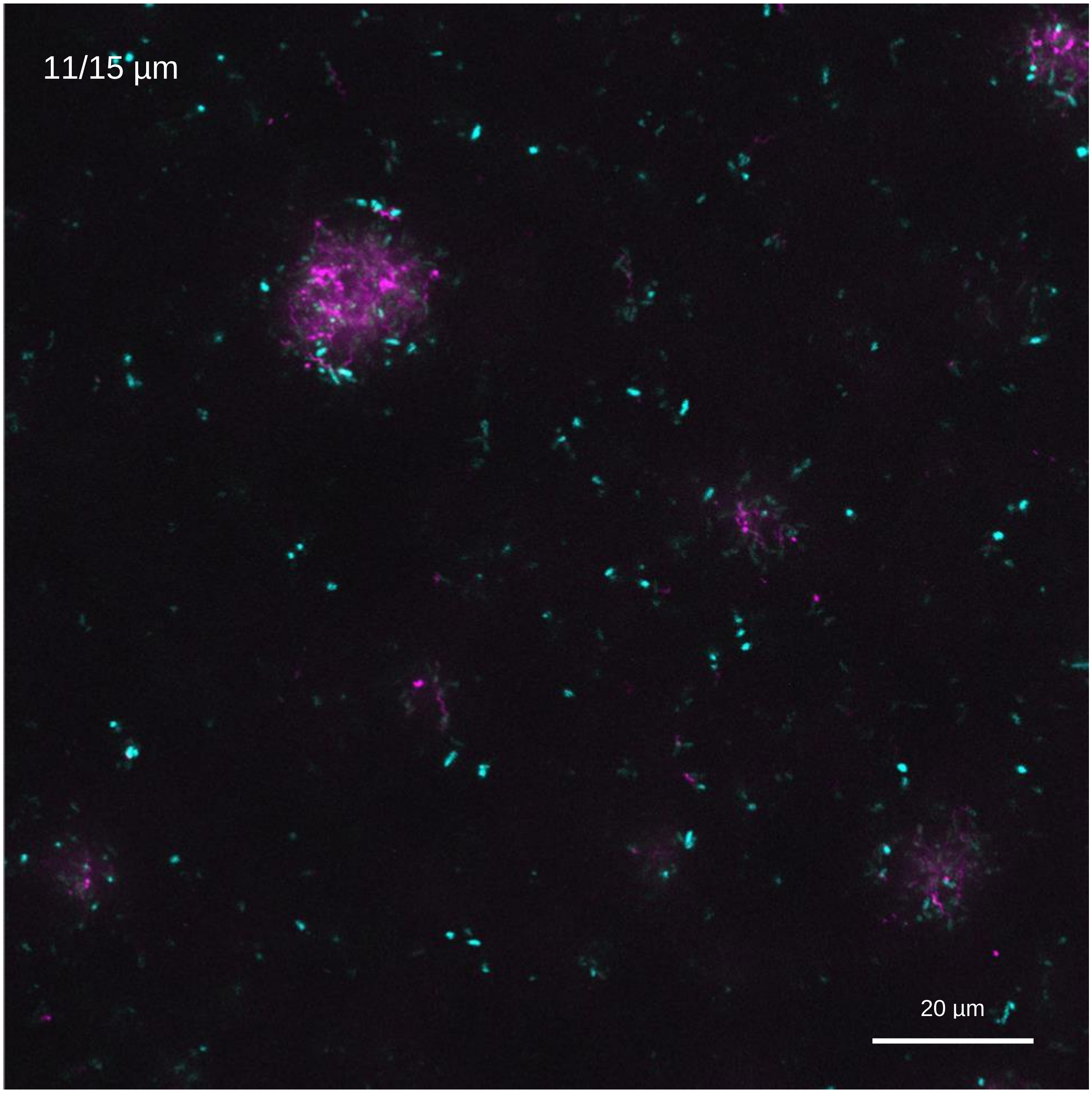


**Figure S16| Young biofilm flagella in upper section.** Enlarged image of stacked layers from a young biofilm upper section after 2 days of growth presenting labeled flagella and bacteria. Channels: λ=488 nm, λ=545 nm

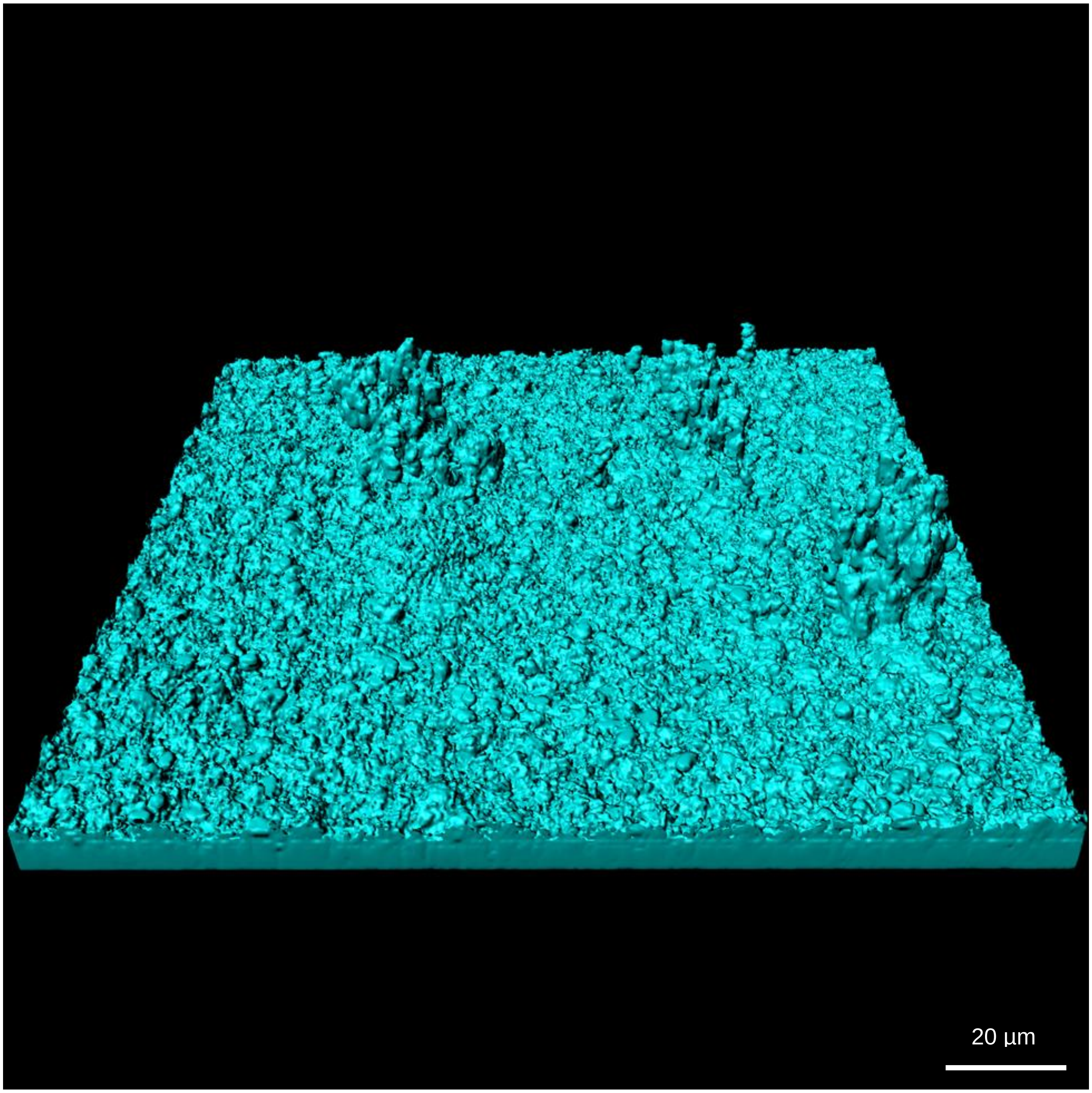


**Figure S17| Growing biofilm biomass.** Enlarged image of visualized biofilm entire biomass after 4 days of growth.

Channel: λ=488 nm

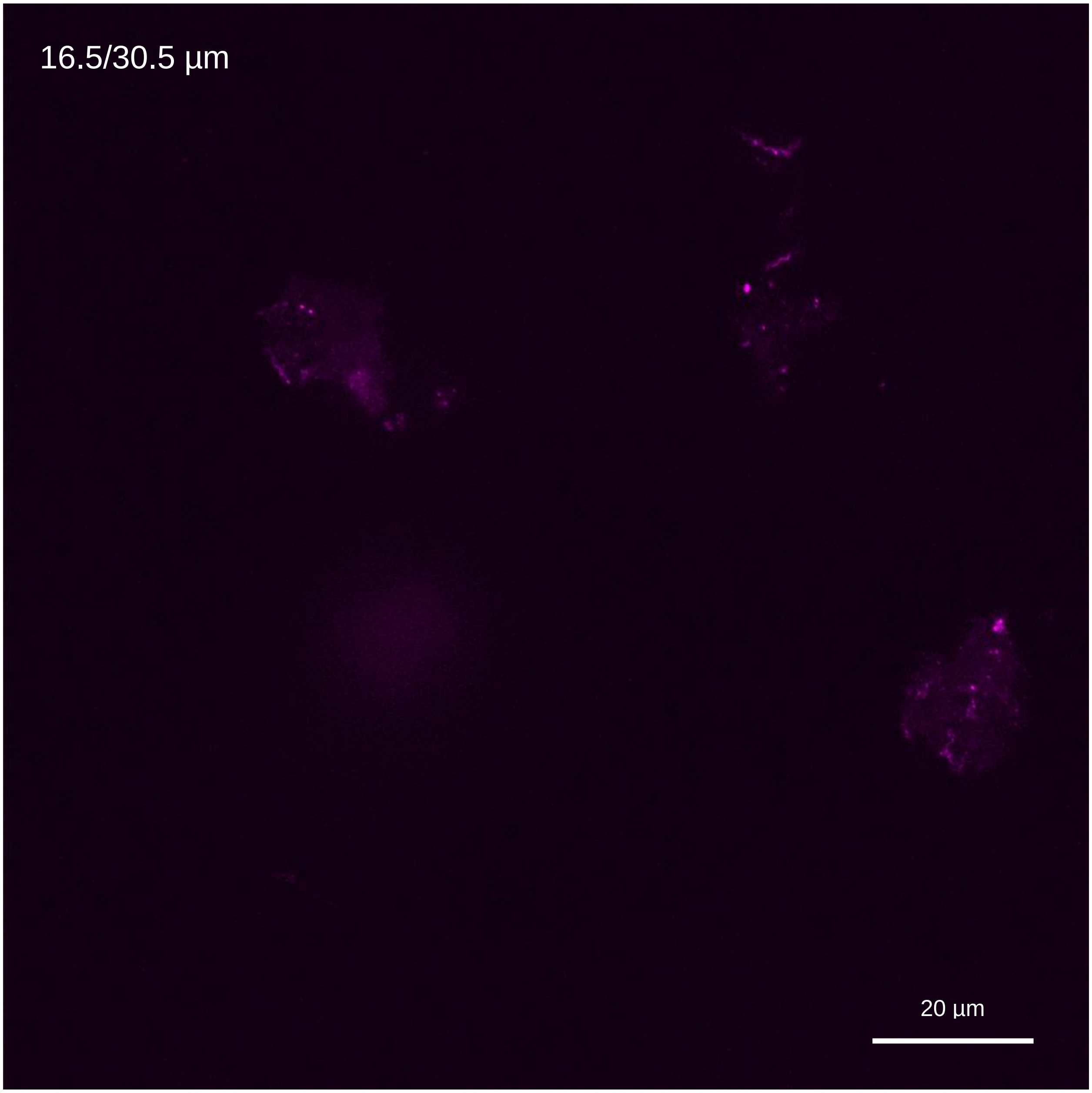


**Figure S18| Growing biofilm flagella in mid-section.** Enlarged image of stacked layers from a growing biofilm mid-section after 4 days of growth presenting labeled flagella. Channel: λ=545 nm

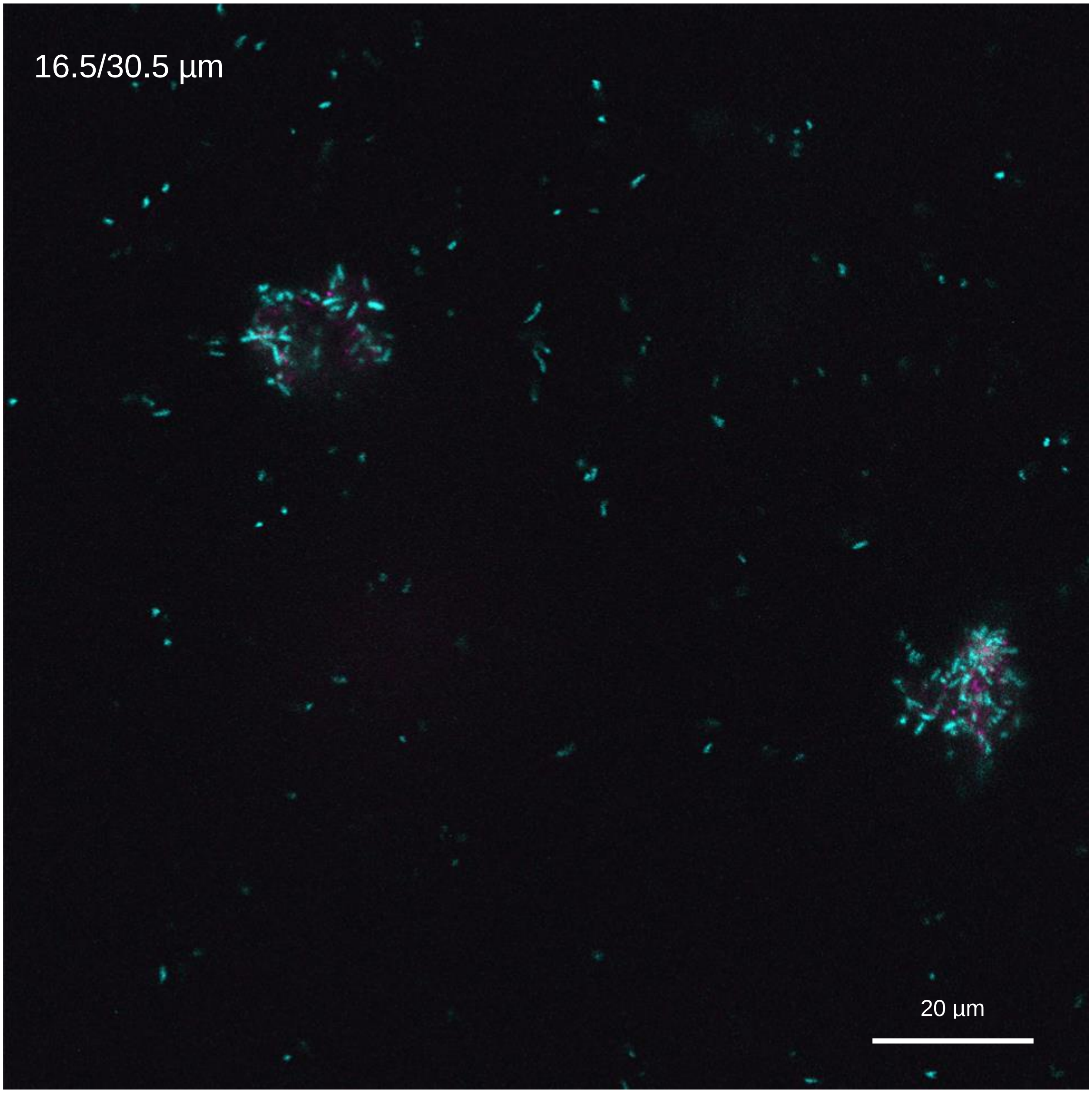


**Figure S19| Growing biofilm flagella in mid-section.** Enlarged image of stacked layers from a growing biofilm mid-section after 4 days of growth presenting labeled flagella and bacteria. Channels: λ=488 nm, λ=545 nm

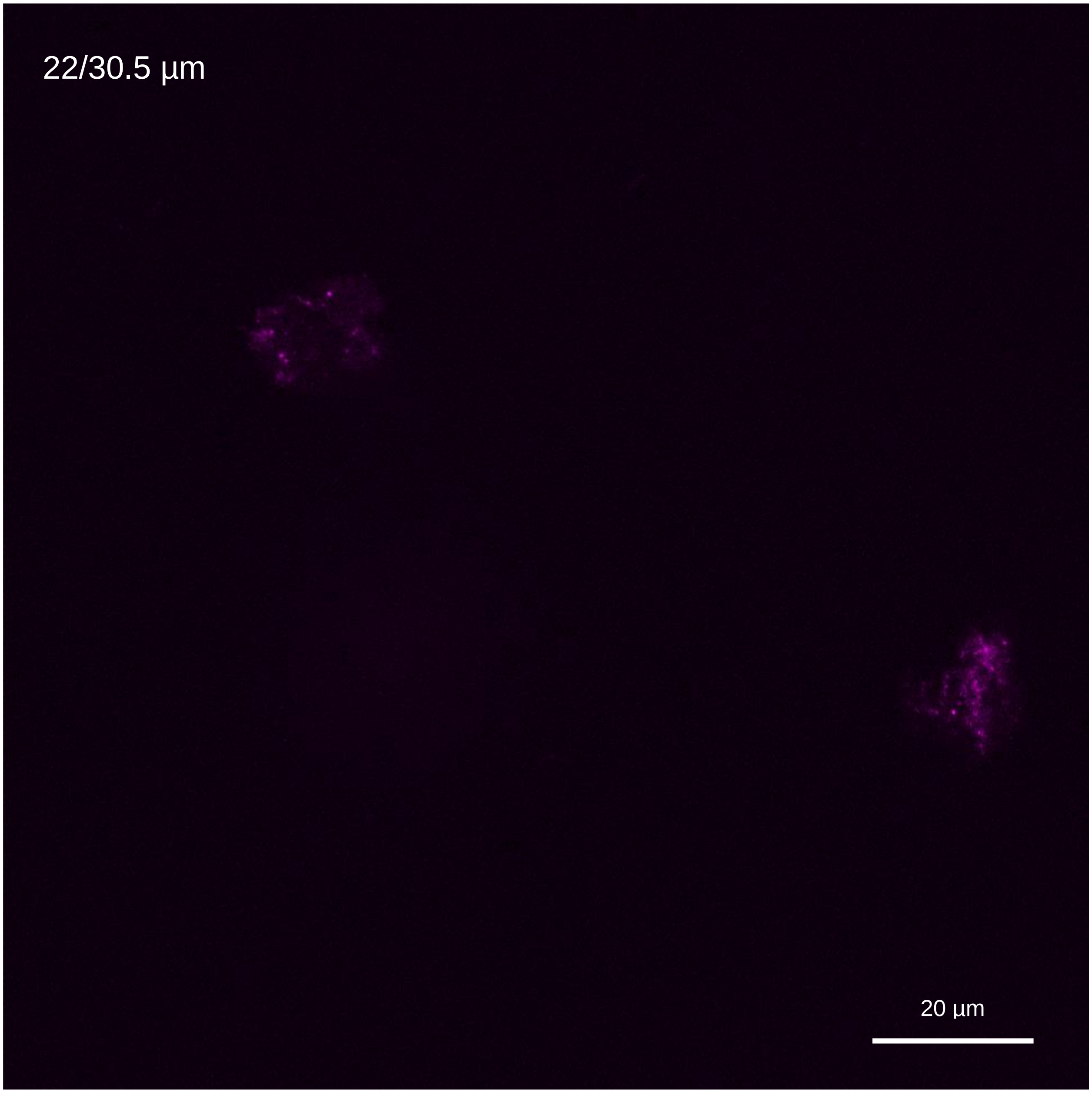


**Figure S20| Growing biofilm flagella in upper section.** Enlarged image of stacked layers from a growing biofilm upper section after 4 days of growth presenting labeled flagella. Channel: $\lambda$=545 nm

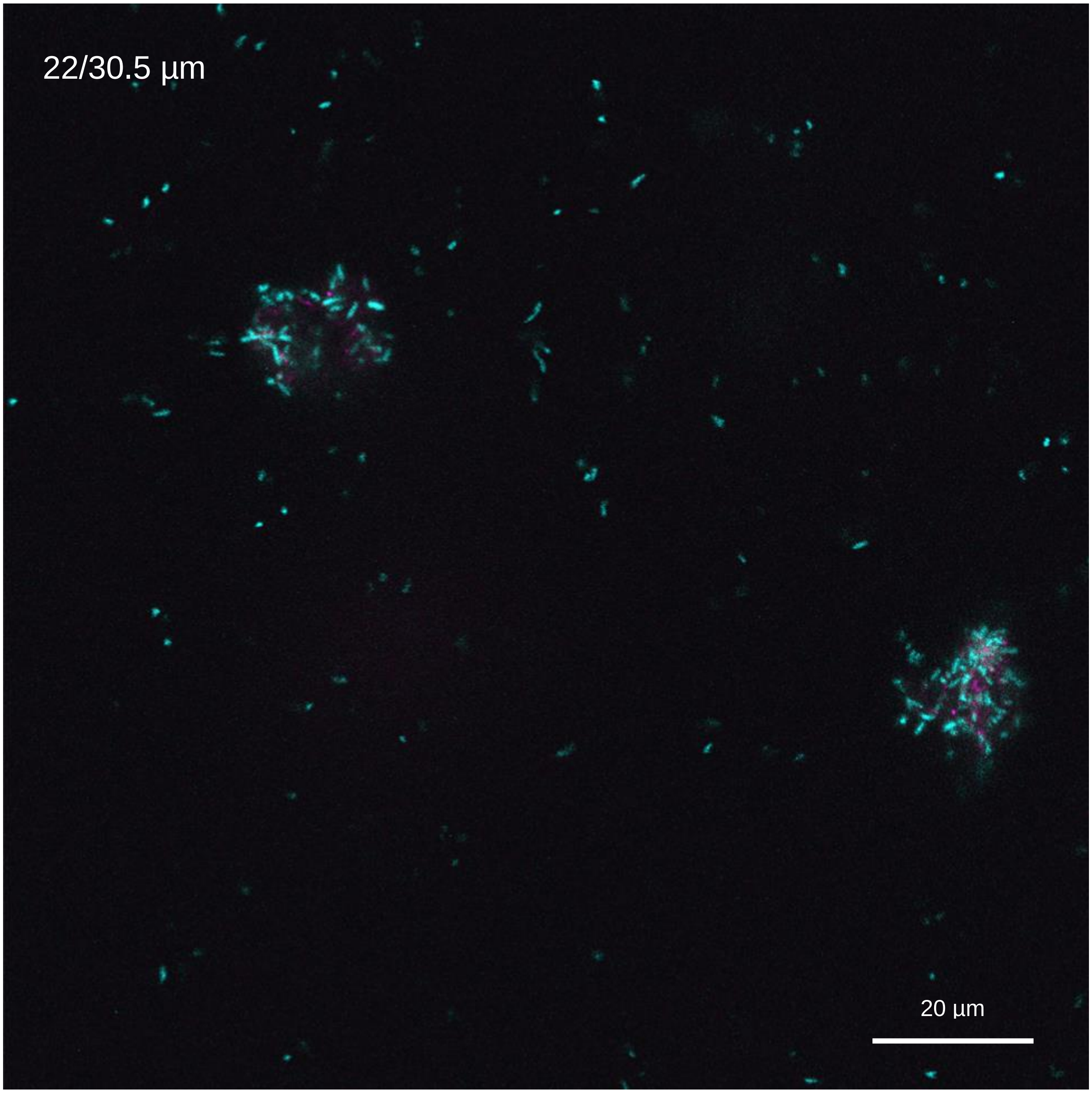


**Figure S21| Growing biofilm flagella in upper section.** Enlarged image of stacked layers from a growing biofilm upper section after 4 days of growth presenting labeled flagella and bacteria. Channels: $\lambda$=488 nm, $\lambda$=545 nm

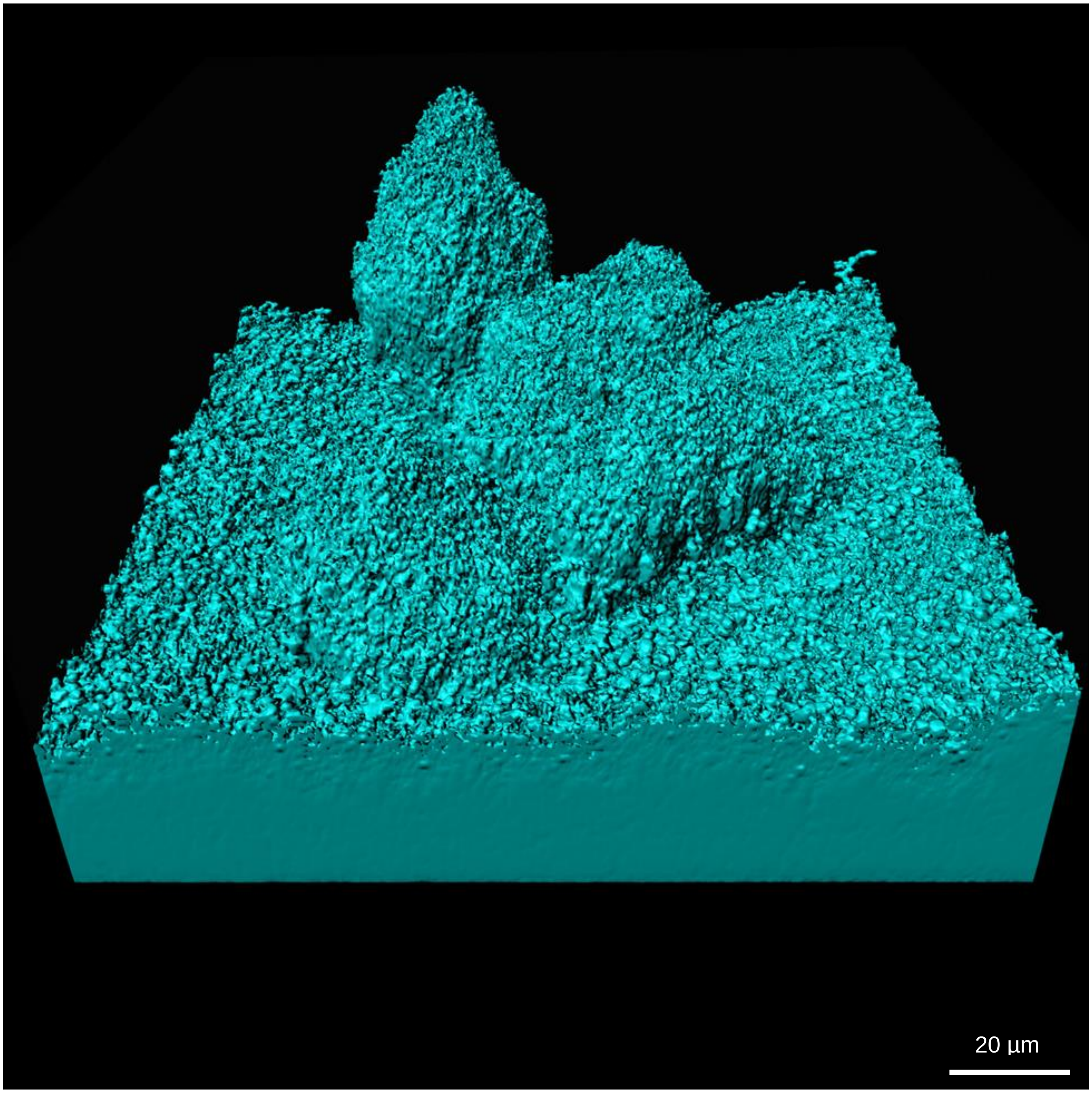


**Figure S22| Mature biofilm biomass.** Enlarged image of visualized biofilm entire biomass after 6 days of growth.

Channel: $\lambda$=488 nm

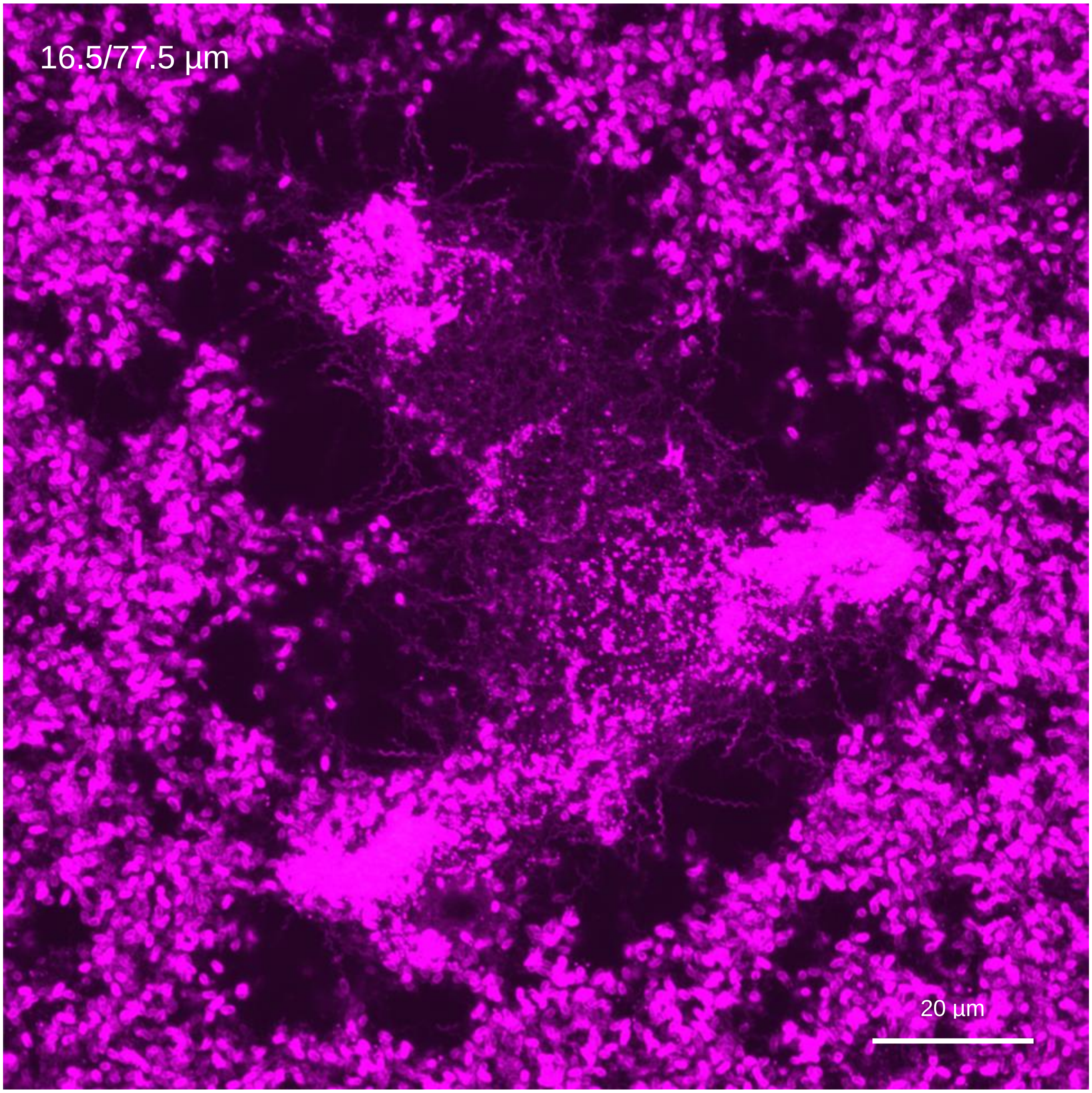


**Figure S23| Mature biofilm flagella in lower mid-section.** Enlarged image of stacked layers from a mature biofilm lower mid-section after 6 days of growth presenting labeled flagella. Channel: λ=545 nm

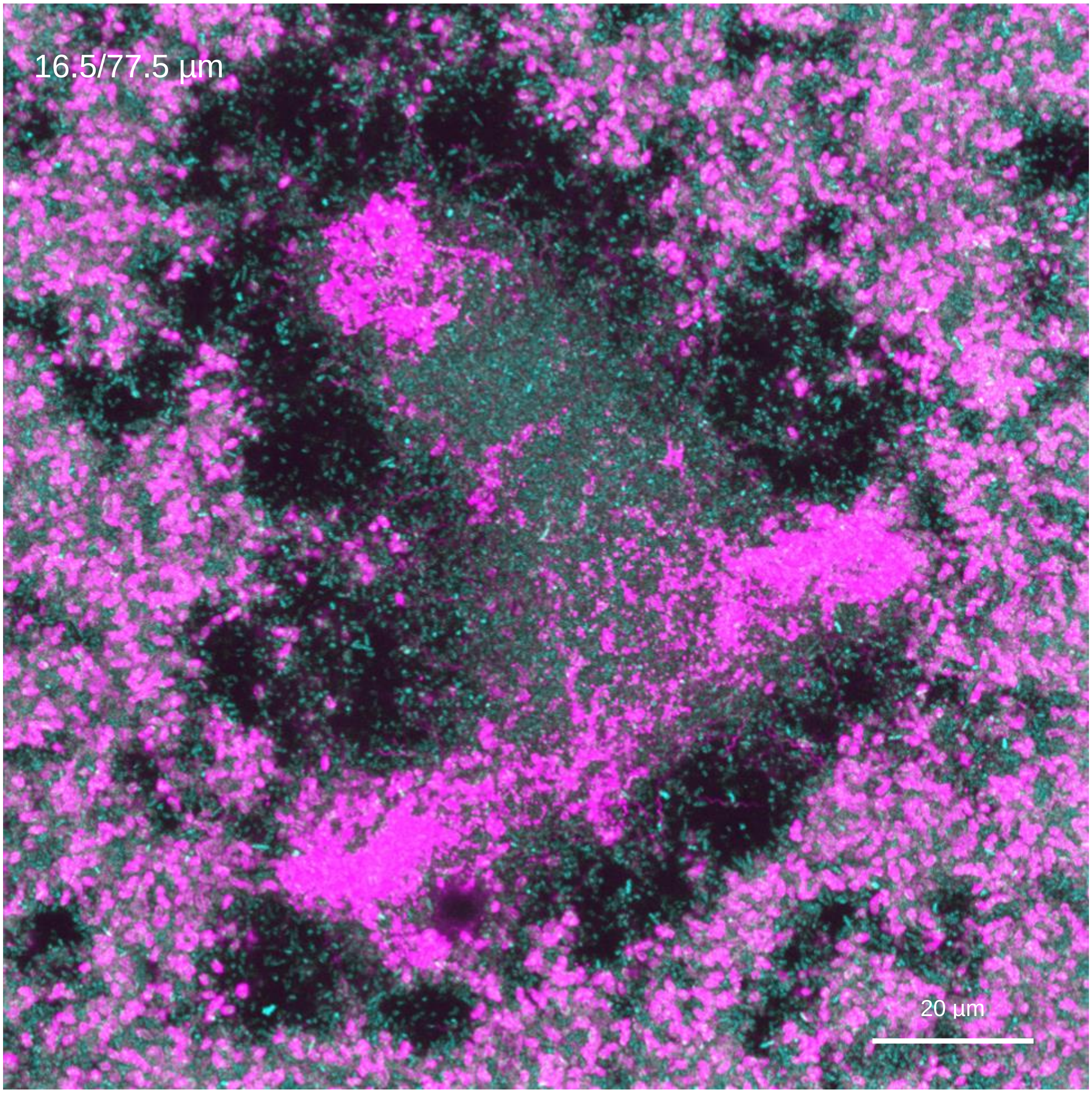


**Figure S24| Mature biofilm flagella in lower mid-section.** Enlarged image of stacked layers from a mature biofilm lower mid-section after 6 days of growth presenting labeled flagella and bacteria. Channels: λ=488 nm, λ=545 nm

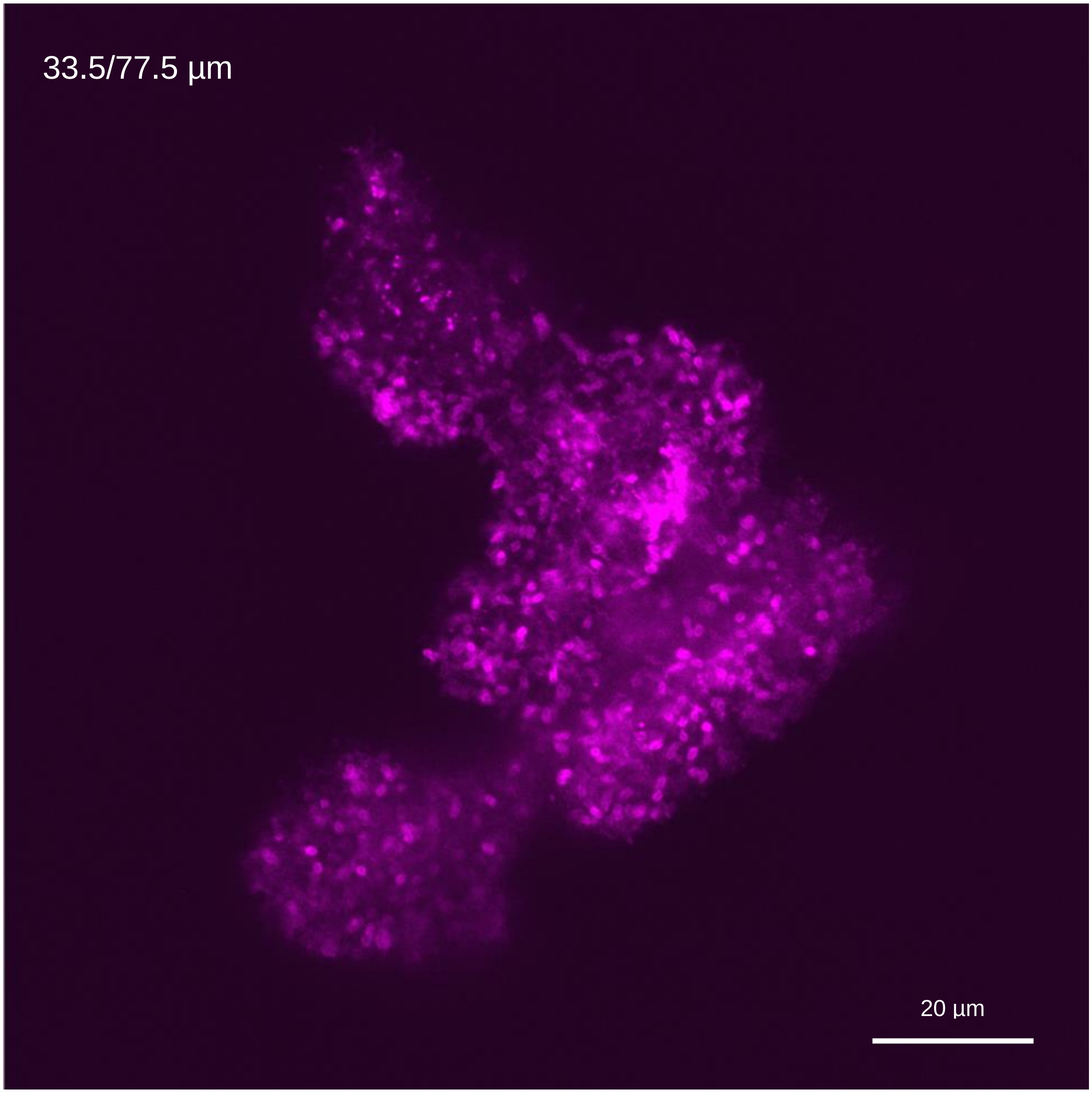


**Figure S25| Mature biofilm flagella in mid-section.** Enlarged image of stacked layers from a mature biofilm mid-section after 6 days of growth presenting labeled flagella. Channel: λ=545 nm

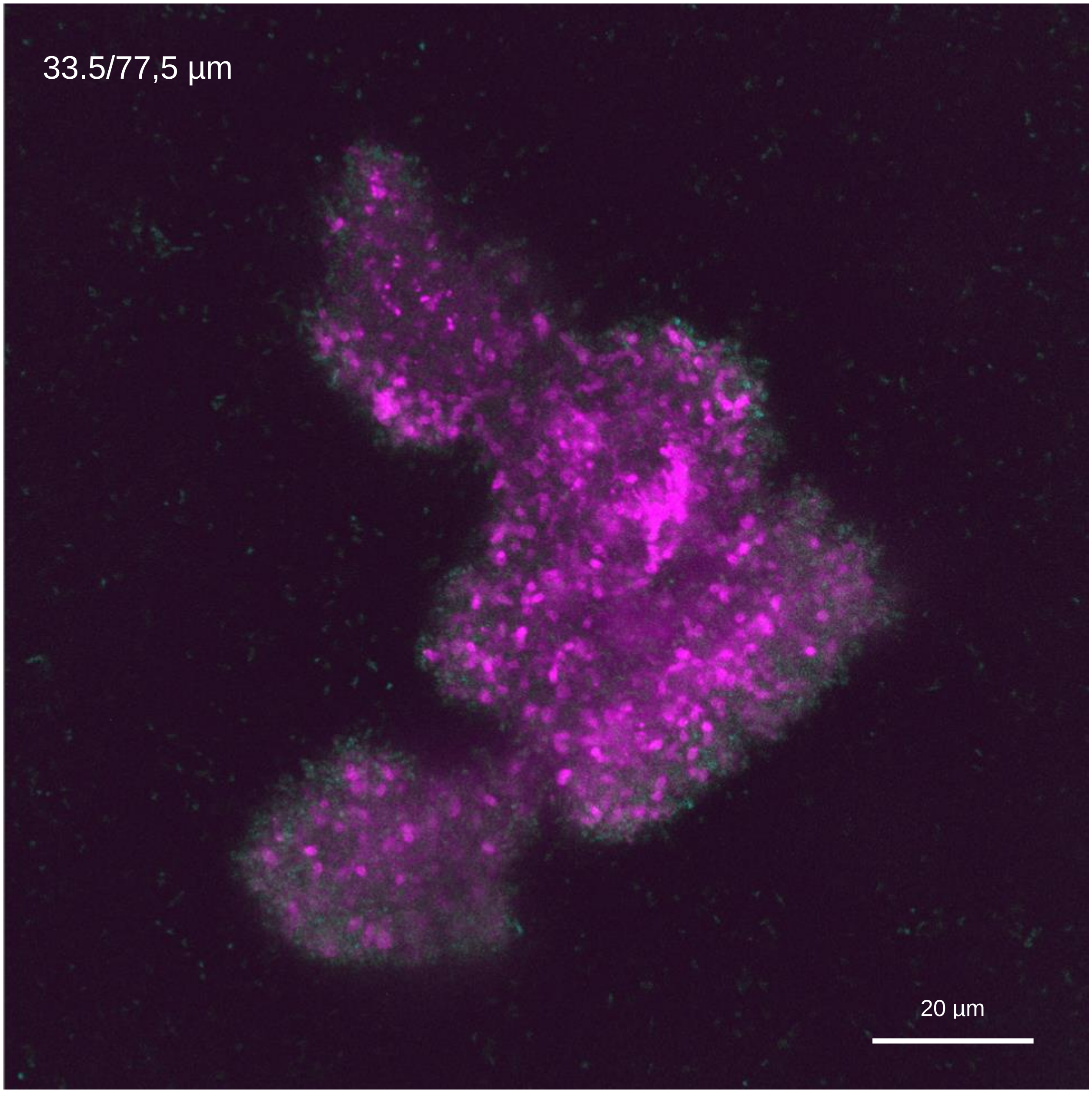


**Figure S26| Mature biofilm flagella in mid-section.** Enlarged image of stacked layers from a mature biofilm mid-section after 6 days of growth presenting labeled flagella and bacteria. Channels: λ=488 nm, λ=545 nm

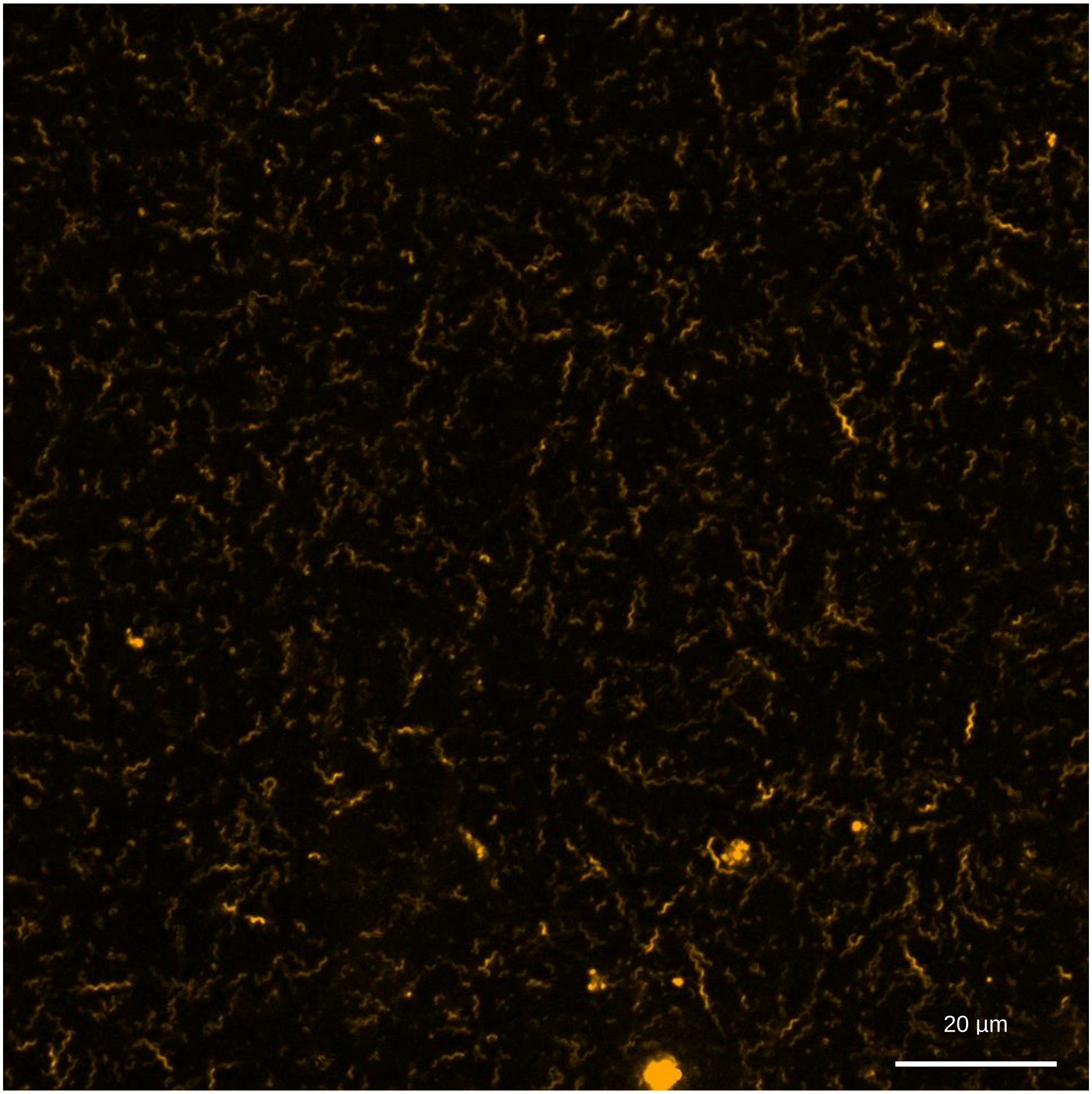


**Figure S27| Young biofilm flagellated bacteria effluent.** Enlarged image of a dispersed cells out of a young biofilm after 2 days of growth. Newly synthesized flagella labeled in orange. Channel: λ=660 nm.

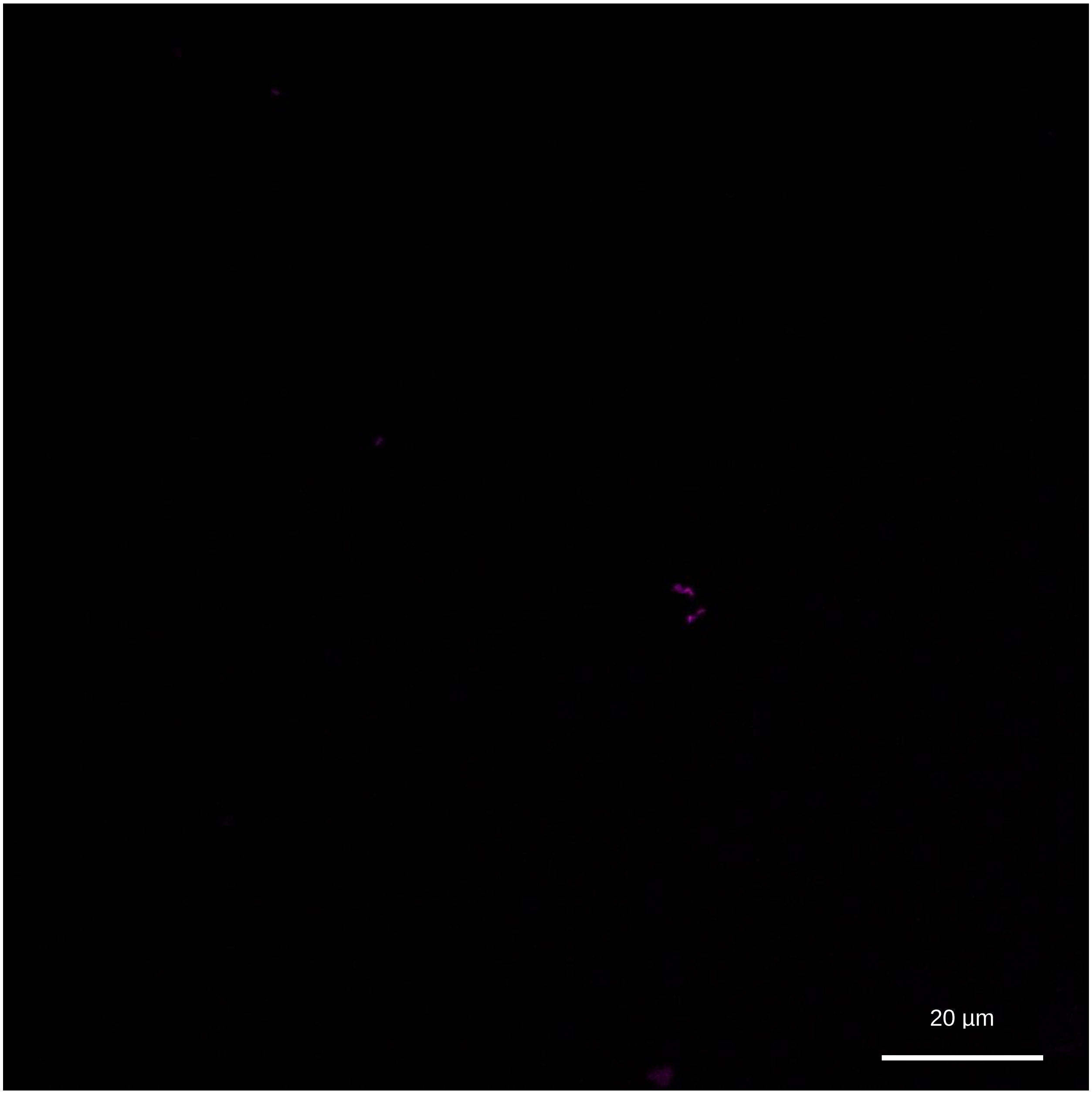


**Figure S28| Young biofilm flagellated bacteria effluent.** Enlarged image of a dispersed cells out of a young biofilm after 2 days of growth. Inoculated flagella labeled in magenta. Channel: λ=545 nm.

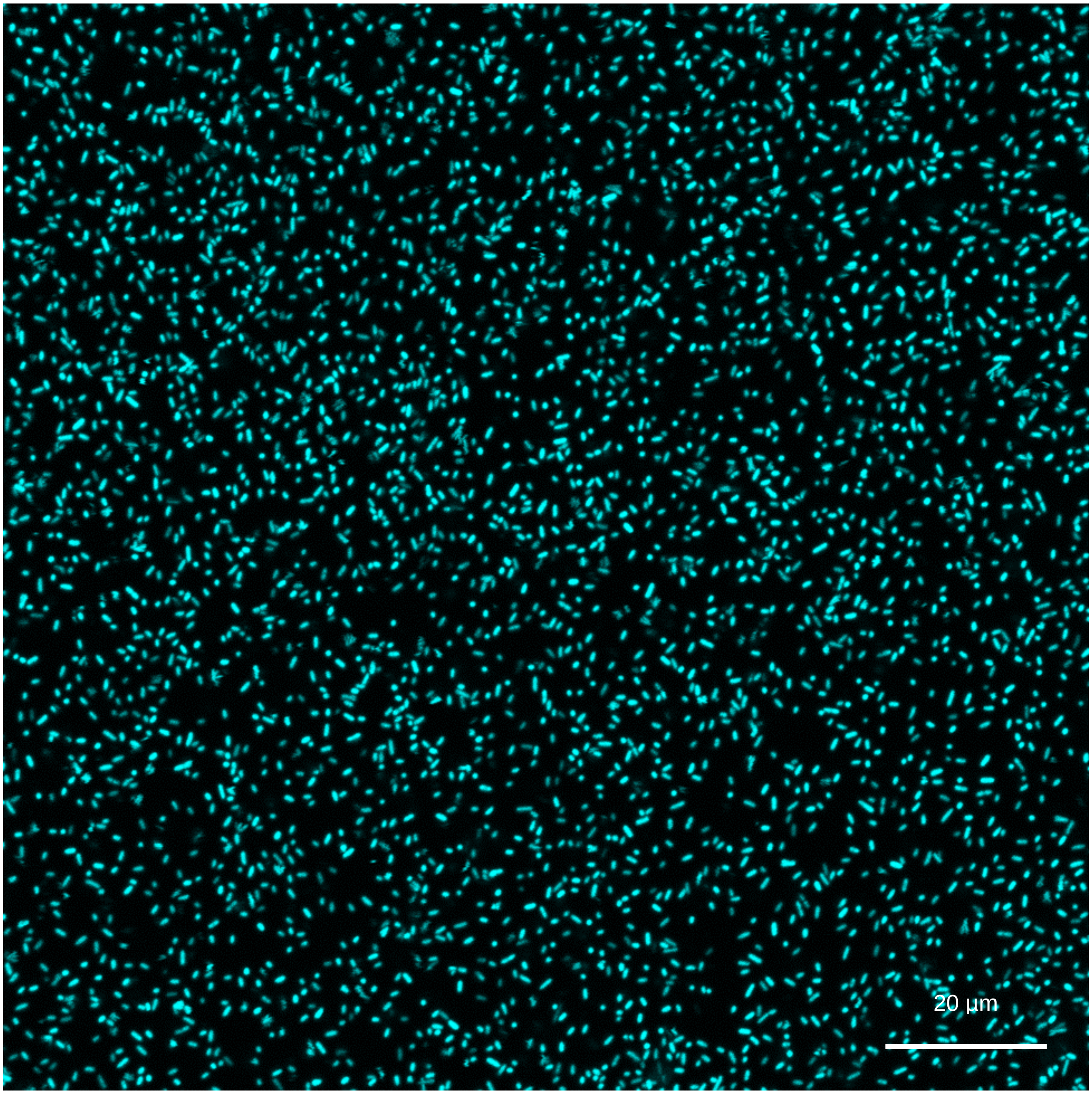


**Figure S29| Young biofilm flagellated bacteria effluent.** Enlarged image of a dispersed cells out of a young biofilm after 2 days of growth. Cells labeled in cyan. Channel: λ=488 nm.

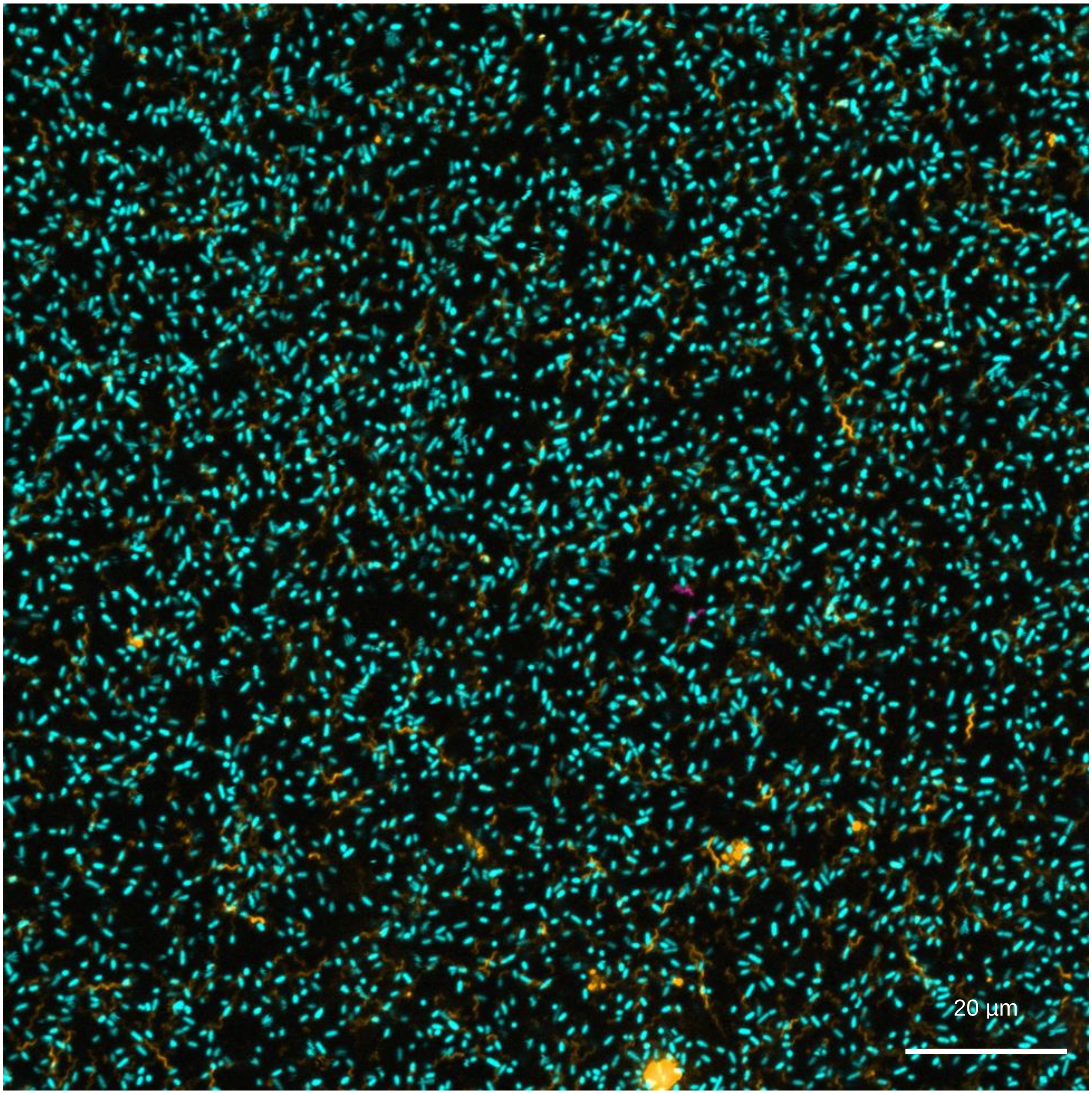


**Figure S30| Young biofilm flagellated bacteria effluent.** Enlarged image of a dispersed cells out of a young biofilm after 2 days of growth. Cells labeled in cyan, inoculated flagella labeled in magenta and newly synthesized flagella labeled in orange. Channels: λ=488 nm, λ=545 nm, λ=660 nm.

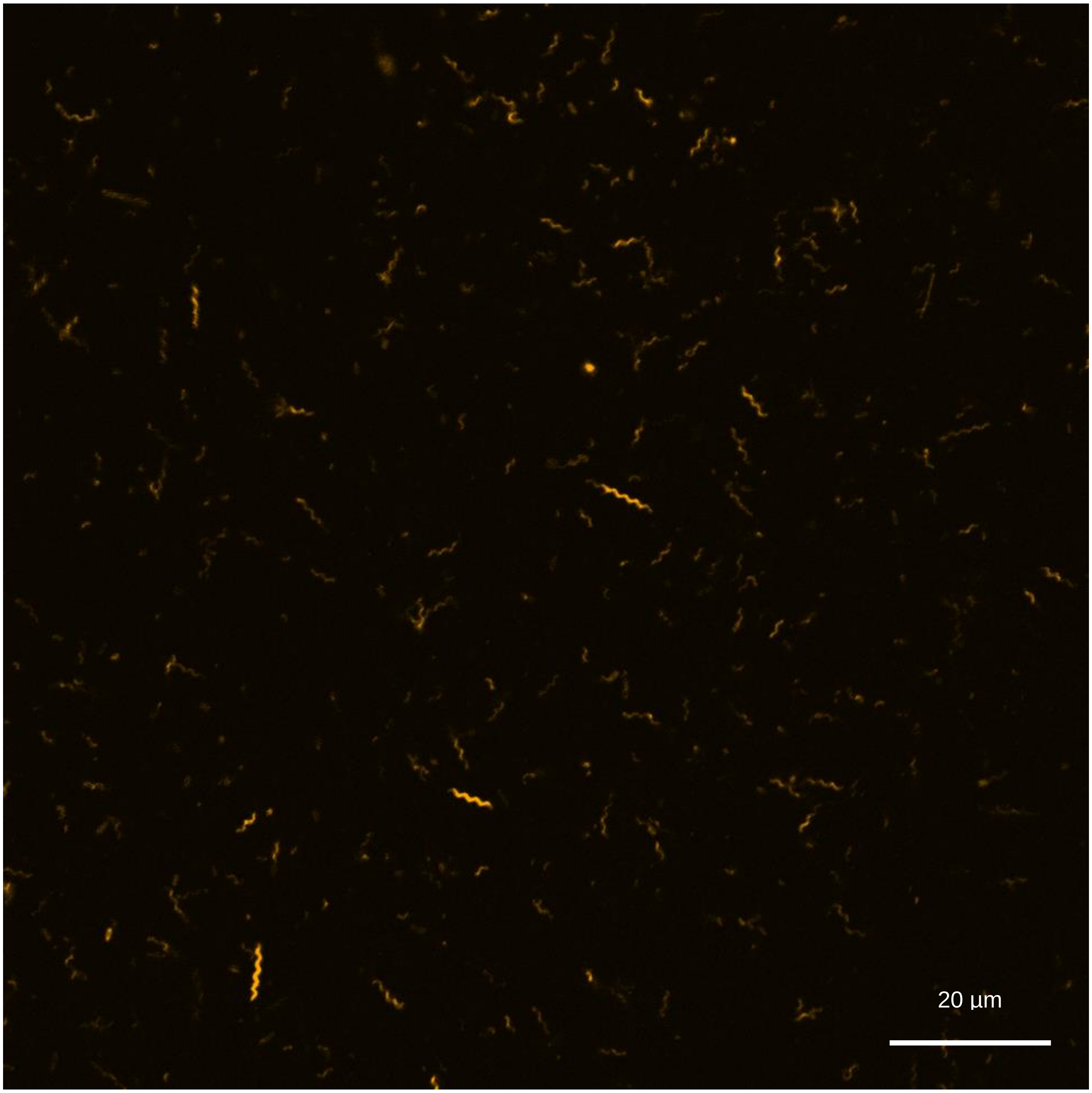


**Figure S31| Growing biofilm flagellated bacteria effluent.** Enlarged image of a dispersed cells out of a growing biofilm after 4 days of growth. Newly synthesized flagella labeled in orange. Channel: λ=660 nm.

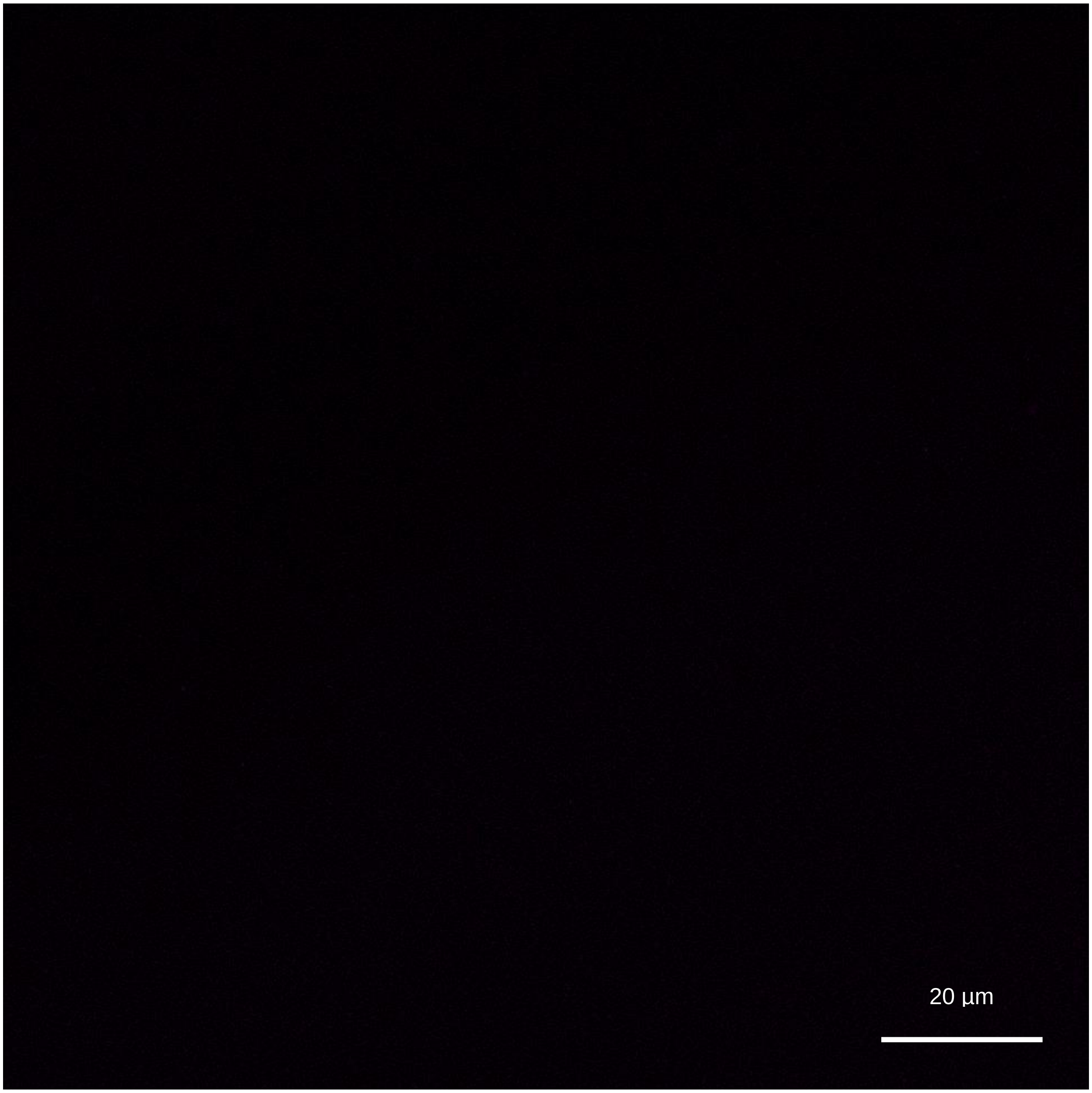


**Figure S32| Growing biofilm flagellated bacteria effluent.** Enlarged image of a dispersed cells out of a growing biofilm after 4 days of growth. Inoculated flagella labeled in magenta. Channel: λ=545 nm.

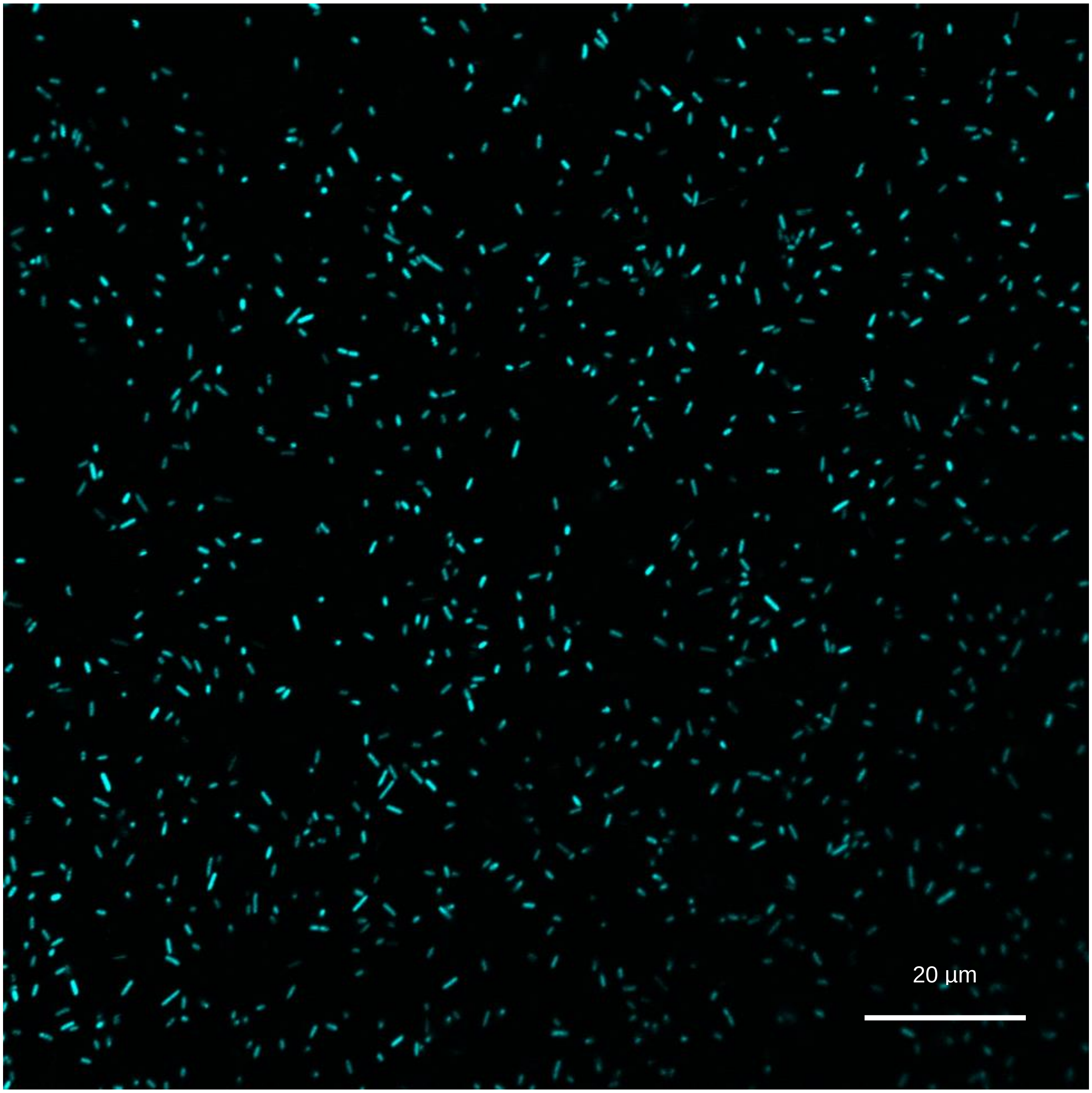


**Figure S33| Growing biofilm flagellated bacteria effluent.** Enlarged image of a dispersed cells out of a growing biofilm after 4 days of growth. Cells labeled in cyan. Channel: λ=488 nm.

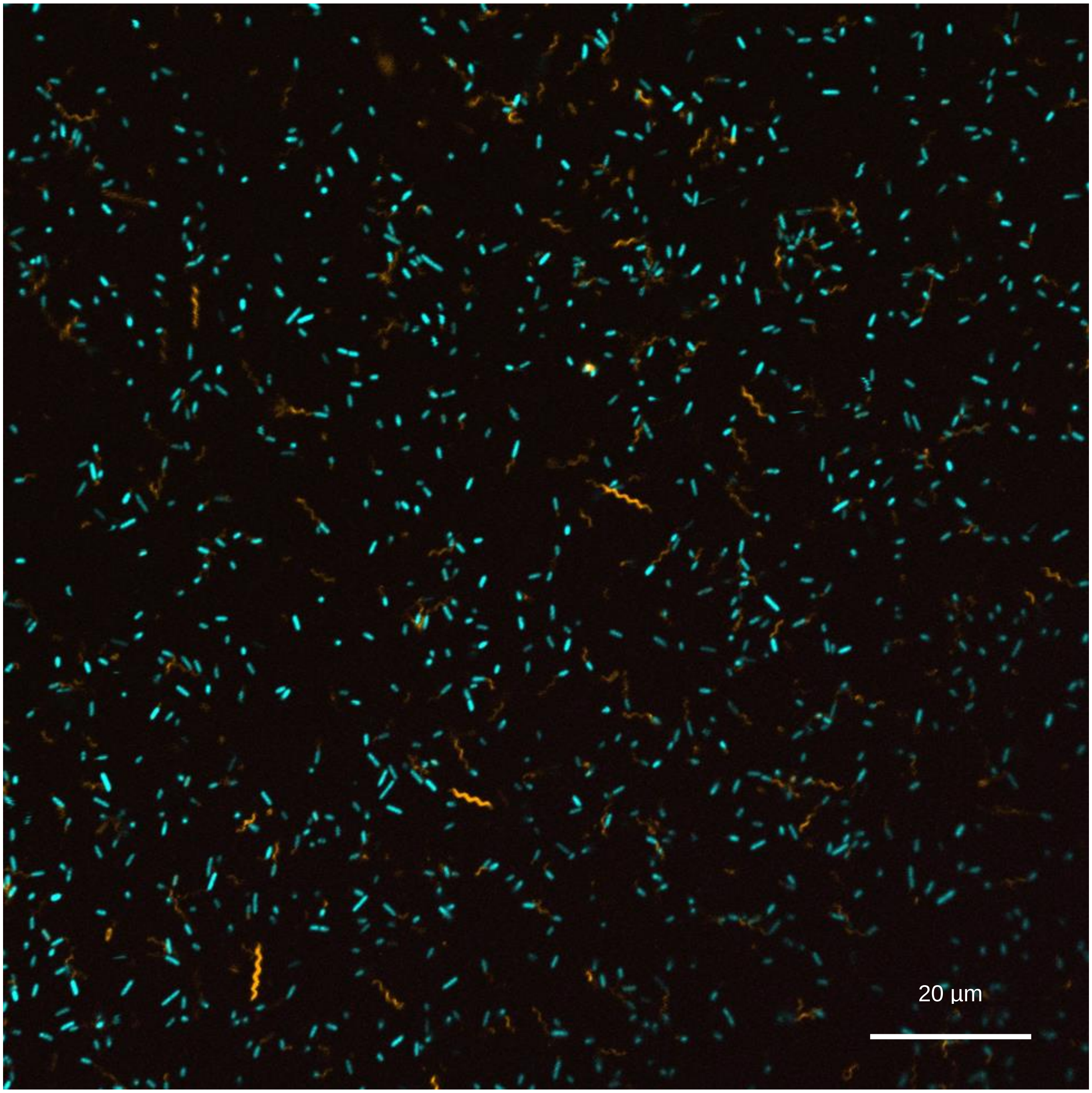


**Figure S34| Growing biofilm flagellated bacteria effluent.** Enlarged image of a dispersed cells out of a growing biofilm after 4 days of growth. Cells labeled in cyan, inoculated flagella labeled in magenta and newly synthesized flagella labeled in orange. Channels: λ=488 nm, λ=545 nm, λ=660 nm.

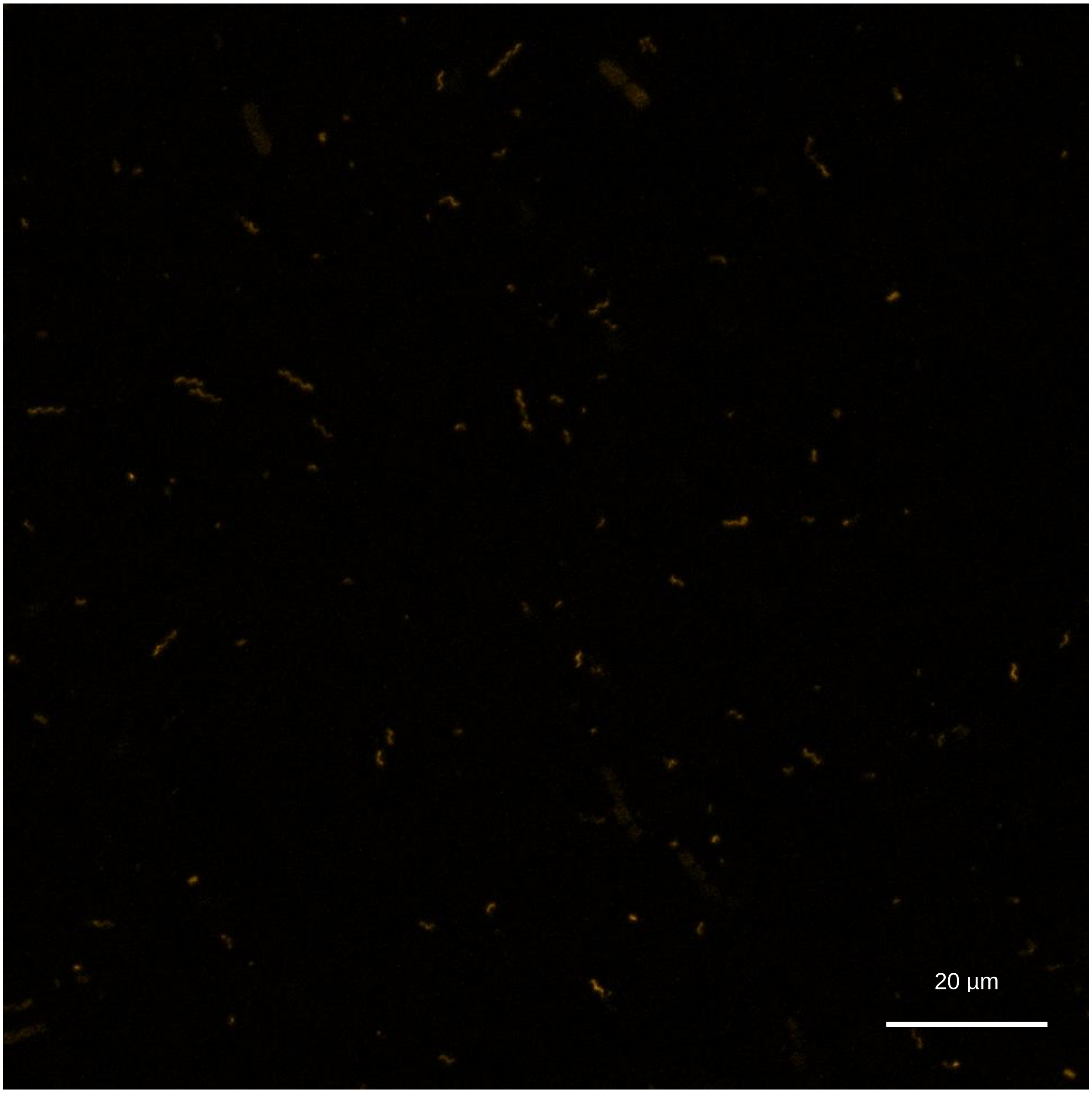


**Figure S35| Mature biofilm flagellated bacteria effluent.** Enlarged image of a dispersed cells out of a mature biofilm after 6 days of growth. Newly synthesized flagella labeled in orange. Channel: λ=660 nm.

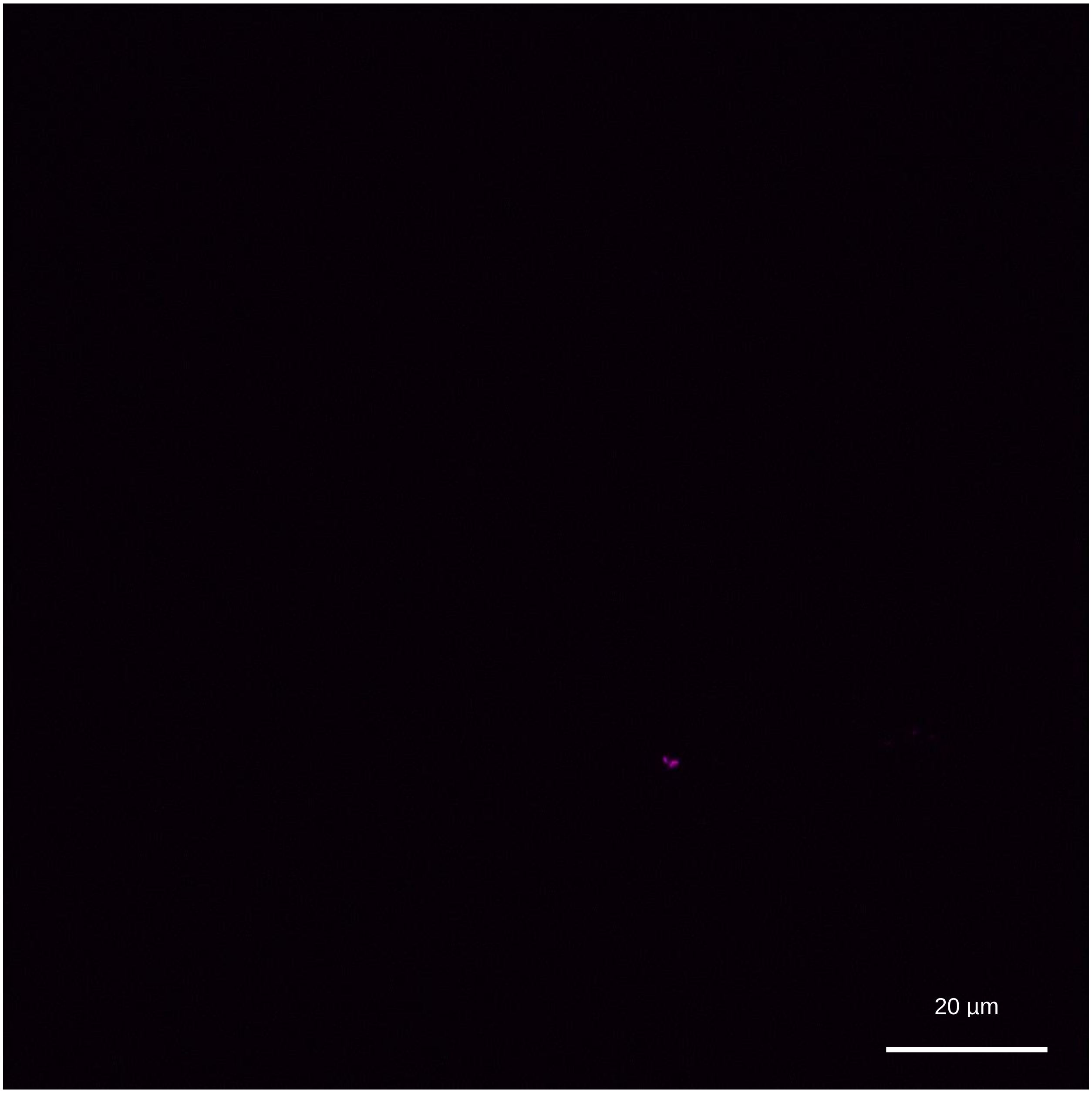


**Figure S36| Mature biofilm flagellated bacteria effluent.** Enlarged image of a dispersed cells out of a mature biofilm after 6 days of growth. Inoculated flagella labeled in magenta. Channel: λ=545 nm.

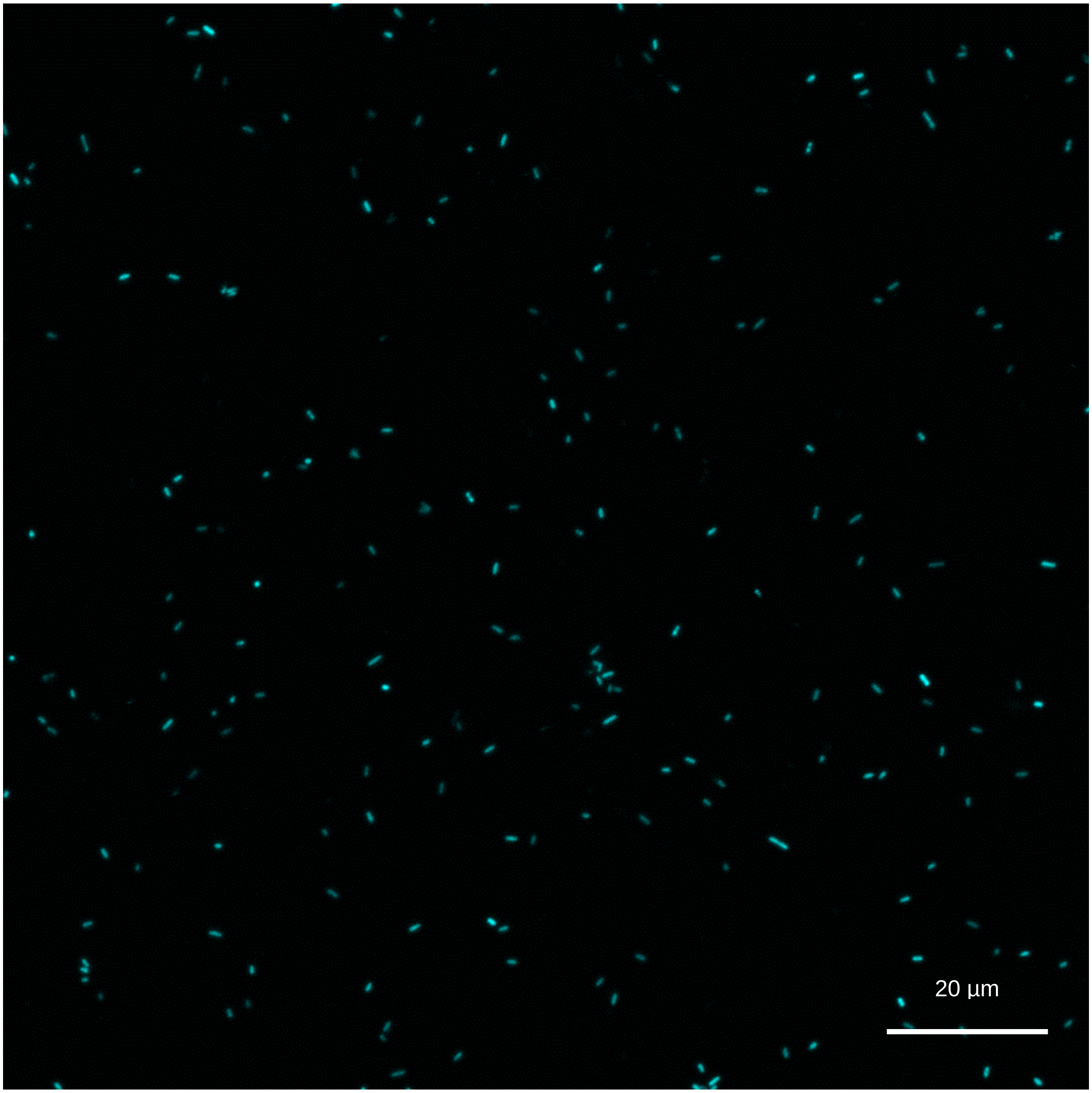


**Figure S37| Mature biofilm flagellated bacteria effluent.** Enlarged image of a dispersed cells out of a mature biofilm after 6 days of growth. Cells labeled in cyan. Channel: λ=488 nm.

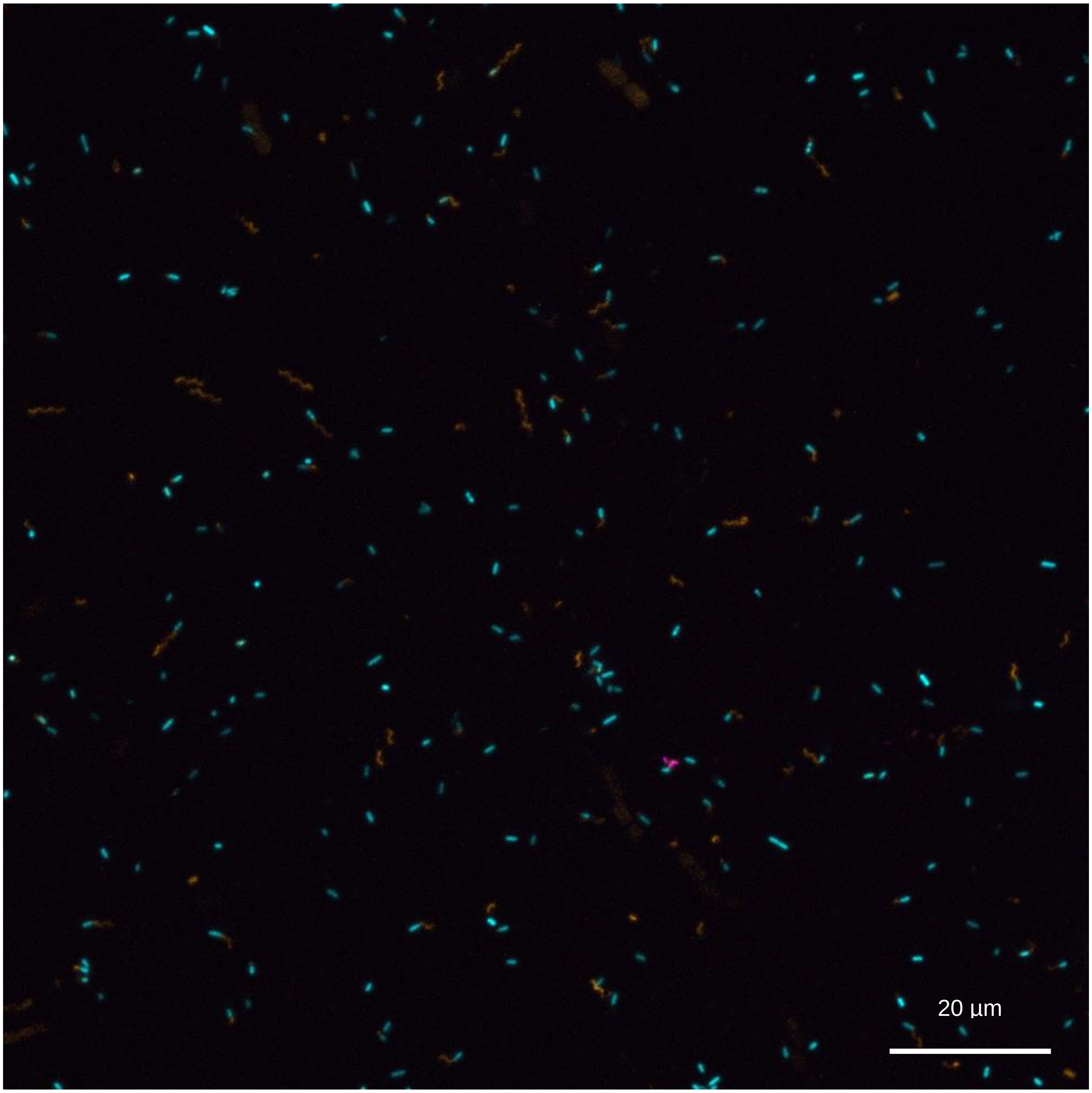


**Figure S38| Mature biofilm flagellated bacteria effluent.** Enlarged image of a dispersed cells out of a mature biofilm after 6 days of growth. Cells labeled in cyan, inoculated flagella labeled in magenta and newly synthesized flagella labeled in orange. Channels: λ=488 nm, λ=545 nm, λ=660 nm.

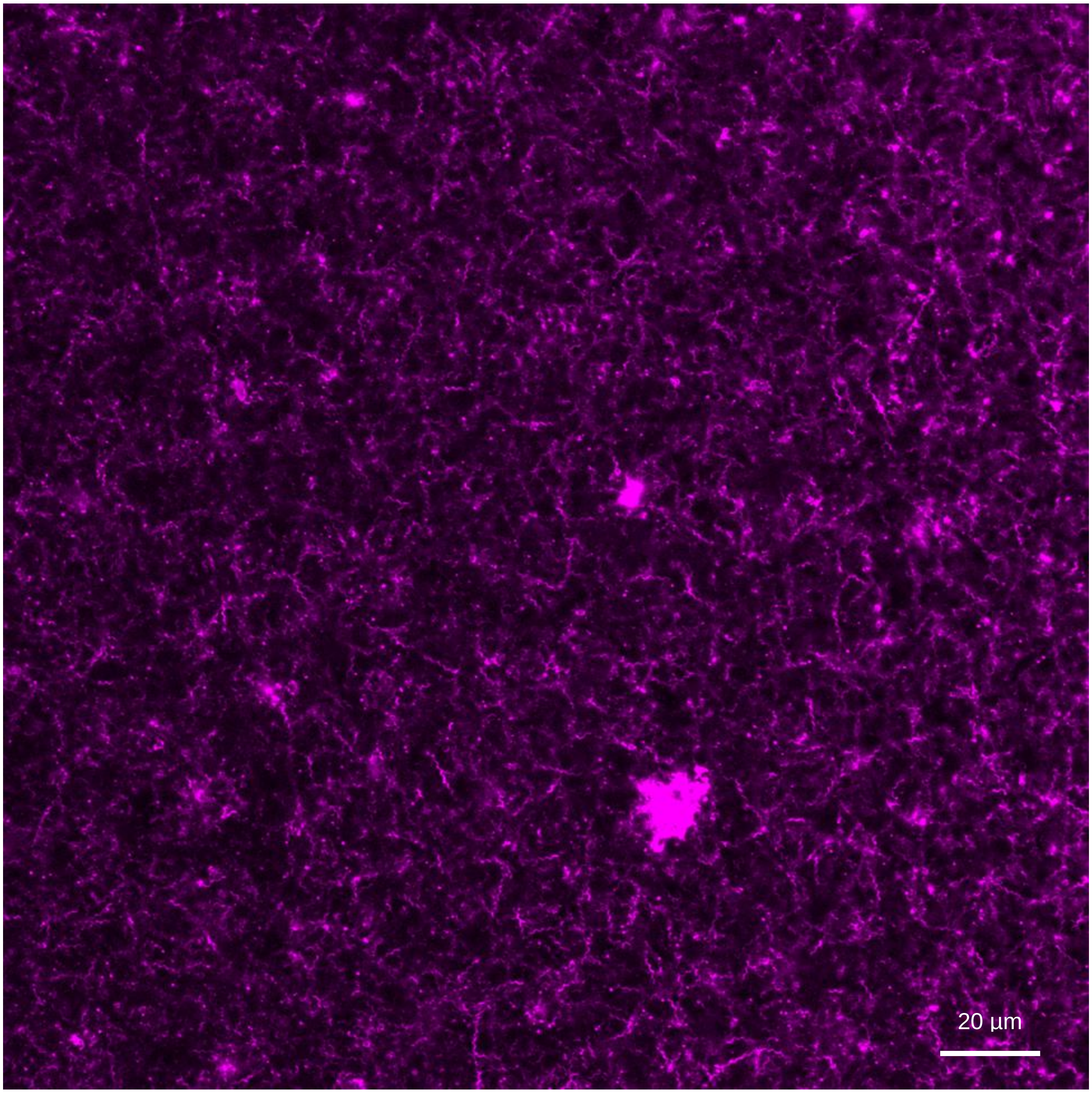


**Figure S39| Flagella in a grid-like structure in a growing biofilm.** CLSM of lower section of a growing biofilm with synthesized flagella demonstrating a grid structure. Channel: λ=545 nm

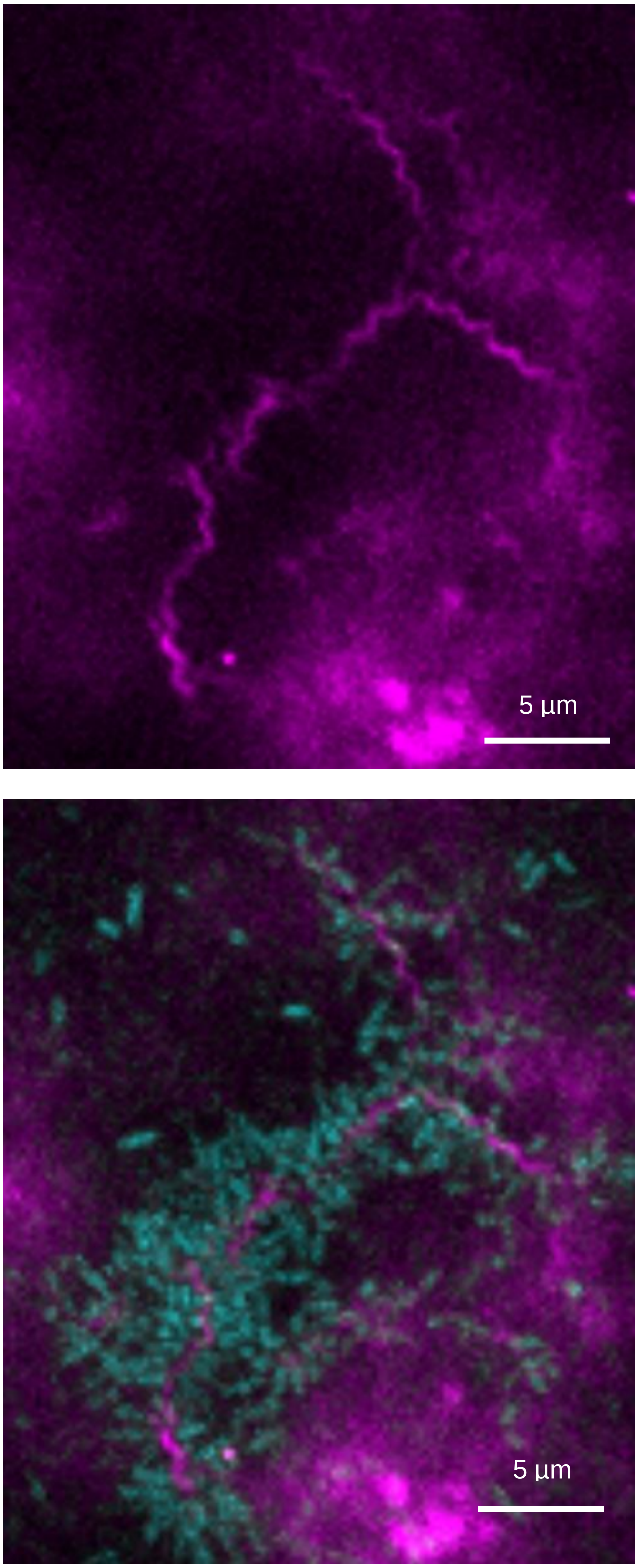


**Figure S40| Bacterial coagulation on flagella.** CLSM of bacterial coagulation on flagella within a growing biofilm.

Channels: λ=488 nm, λ=545 nm

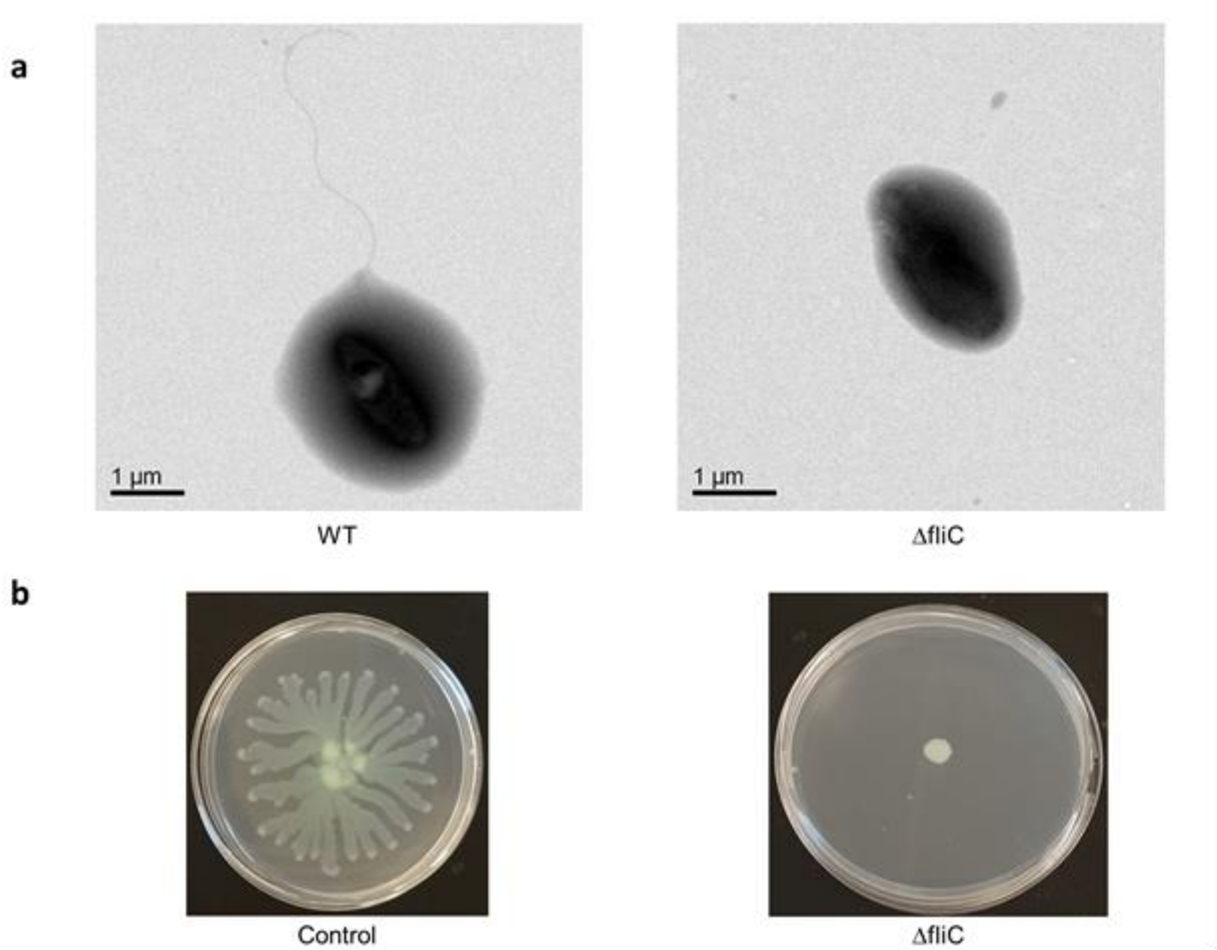


**Figure S41| Δ*fliC* validation for lacking flagella.** (a) TEM imaging of WT bacteria possessing flagella and Δ*fliC* bacteria lacking flagella. (b) Swarming assay demonstrating Δ*fliC* variant flagella absence.

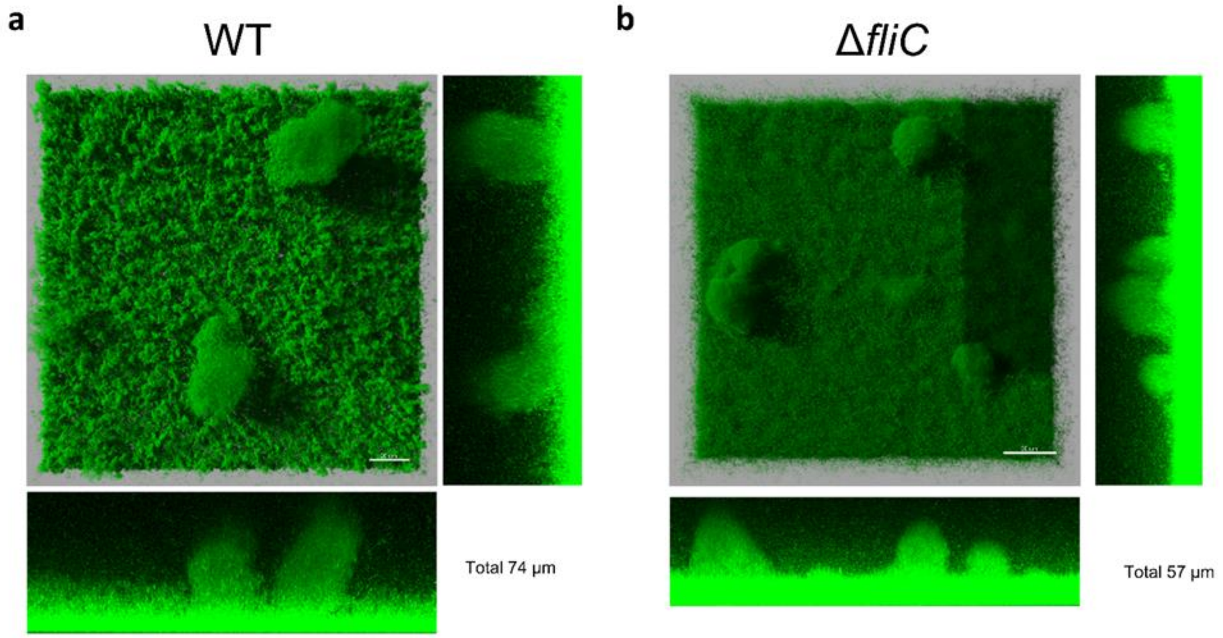


**Figure S42| Biofilm structure comparison through CLSM.** Comparison between mushroom-like structures inside biofilms generated with WT variant vs. Δ*fliC* variant.

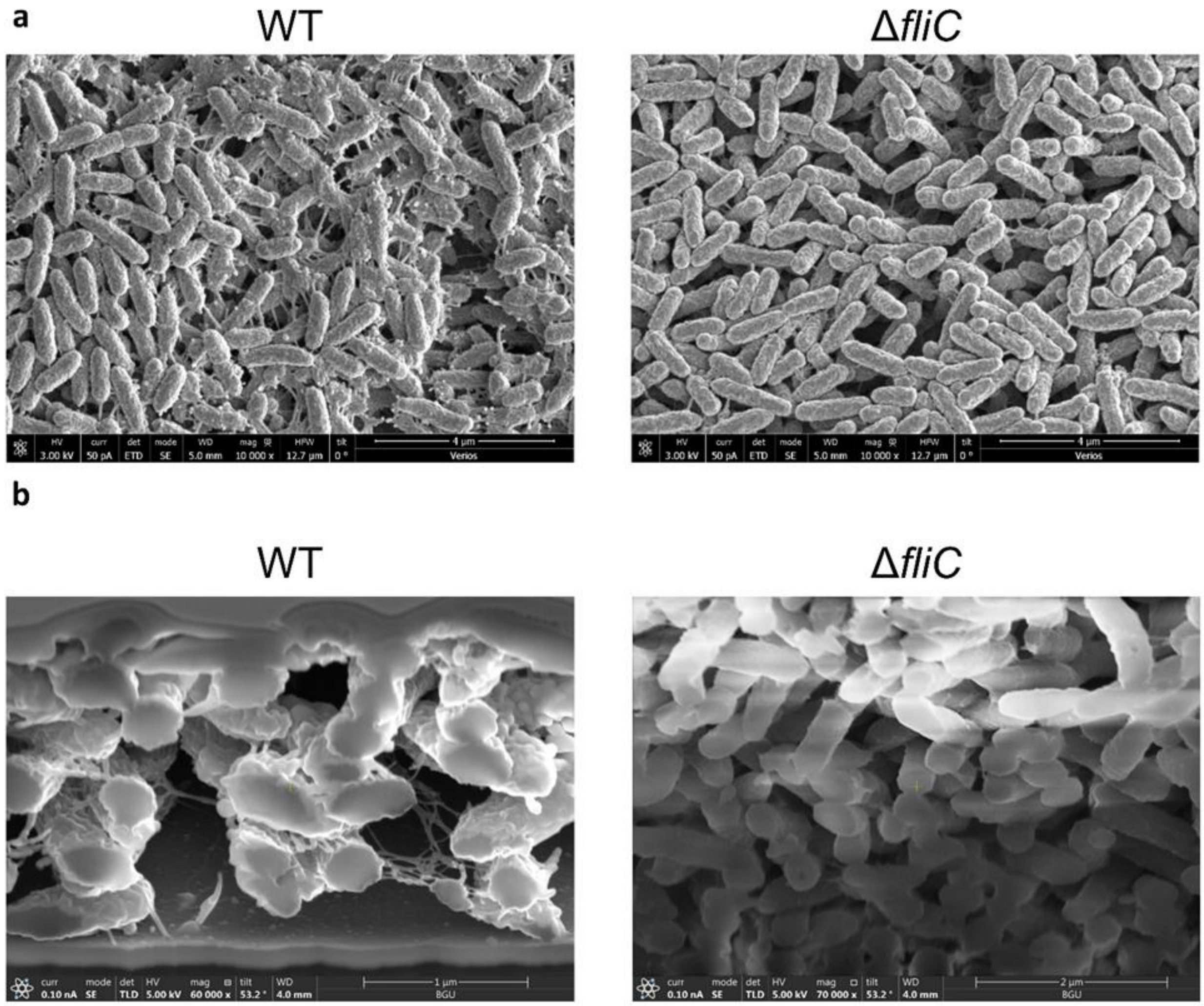


**Figure S43| Biofilm structure comparison trough HR-SEM.** Comparison between biofilms structures generated with WT variant vs. Δ*fliC* variant, through HR-SEM (a) or an inside look through Dual beam HR-SEM (b).

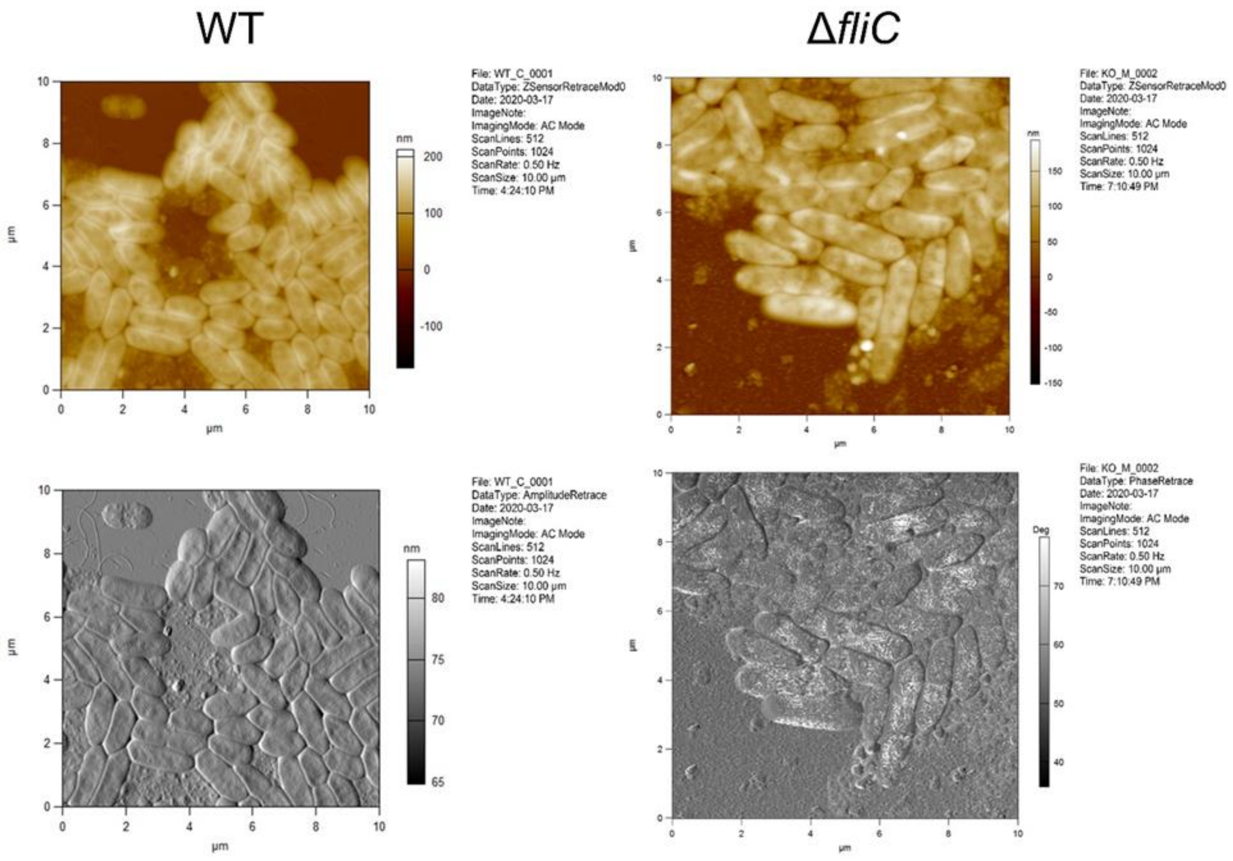


**Figure S44| Biofilm structure comparison trough AFM.** Comparison between biofilms generated with WT variant vs. Δ*fliC* variant, through AFM.

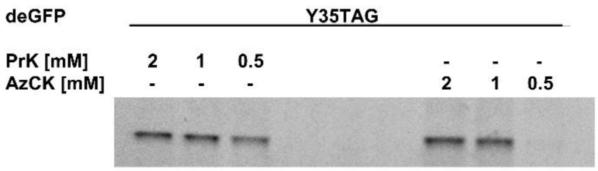


**Figure S45| Optimized Uaa concentration for incorporation.** Fluorescent SDS-PAGE examining PrK and AzCK incorporation into GFP. Samples were lysed and underwent CuAAC click reaction to an azide-containing (for PrK) or alkyne-containing (for AzCK) fluorescent dye.

**Plasmids and genes used in this study:**

<u>pPaGE Pyl TAG *fliC* prom deGFP WT NHis:</u>

TTATCCCCTGATTCTGTGGATAACCGTATTACCGCCTTTGAGTGAGCTGATACACTAGTAGTTTTGCTGGCCGCATCTTCAGAAATT
GACCAGCTGCGAAAGTGGTTTCAACTGGAAGGCCGGTATCCCTCGATCAAGGACTTCAAGTTGCGAGTGCTTGATCCAGCCGTGA
CGCAGATCAACGAGCACAGCCCGCTACAGGTGGAGTGGGCGCAGCGAAAGACCGGGCGCAAGGTCACACATCTGTTGTTCAGTT
TTGGACCGAAGAAGCCCGCCAAGGCGGTGGGTAAGGCCCCAGCGAAGCGCAAGGCCGGGAAGATTTCAGATGCTGAGATCGCGA
AACAGGCTCGCCCTGGTGAGACATGGGAAGCGGCCCGCGCTCGACTAACCCAGATGCCGCTGGATCTGGCCTAGAGGCCGTGGC
CACCACGGCCCGGCCTGCCTTTCGGATCCTAGAGGTAGGTCACCTTTTCTTCTTTAGCGTTTGCTGCCATTGCTCTATTGAAGCGCG
AAAACGCTTCGTCGGTCAGCTTCGGCAACAGAATAATTTTTATCGCATCAGGGGTTTACAAGCAGAATACGTACTCGTATAGTTCG
CGCCCACAACGTTGCTGAGACGTTGTAACACGGAAACGATGAGCGATCATCGACTCCAGGGAAACCTGATCATGTAGATCGAATG
GACTCTAAATCCGTTCAGCCGGGTTAGATTCCCGGGGTTTCCGCCAATATTTCAGACAAAGCCGGTCATTAGACCGTTTTTGTTTC
AGCAGTTCAGCCACTACCAGGCATTGAAACTGCGACAAGGTTTGCAACACCCAAGTCGATGAGCGATCATCGCACAATGCTGAAA
AGCACGAACACCTCTGAAGGTACCACTGGGCAGCCAGCCATCGACGGCCCATAGGGCAAGCTTAGCGTTCCCGTTCGGCTGAACC
GCCCCCGGCGACAAACGGCGCGGGCTATCGACAGGAGTGGGGGTCATGTATGCTACGAGGCTTTCGAACAGTCACGTCTAAGTA
GCGCCCCGCGGATGGATAAAAAACCACTAAACACTCTGATCTCTGCTACTGGTCTGTGGATGAGTCGTACCGGAACCATTCATAA
AATCAAACACCACGAGGTTAGCCGTTCGAAAATCTATATTGAGATGGCGTGTGGCGATCATCTGGTTGTGAACAATAGCCGCTCT
TCTCGTACAGCACGTGCACTGCGTCACCACAAATATCGTAAAACCTGTAAACGTTGCCGTGTGTCCGATGAGGATCTGAACAAAT
TCCTGACAAAAGCCAATGAGGACCAAACAAGCGTGAAAGTGAAAGTCGTTAGCGCTCCTACCCGTACTAAAAAAGCAATGCCGA
AATCCGTTGCTCGTGCCCCTAAACCACTGGAAAACACTGAAGCAGCACAGGCACAGCCGTCTGGAAGCAAATTCTCTCCGGCCAT
TCCTGTTTCTACCCAGGAGTCCGTTTCTGTTCCAGCAAGTGTGAGCACCAGCATTAGCAGTATTAGCACCGGTGCCACCGCTAGCG
CCCTGGTTAAAGGCAATACCAATCCGATTACAAGCATGTCTGCCCCGGTTCAAGCATCAGCTCCAGCACTGACAAAATCCCAAAC
CGATCGTCTGGAGGTTCTGCTGAATCCGAAAGACGAAATCAGCCTGAATTCCGGCAAACCGTTTCGTGAACTGGAGAGCGAACTG
CTGTCACGTCGTAAAAAAGACCTGCAACAAATCTATGCCGAAGAACGTGAGAACTATCTGGGGAAACTGGAACGTGAAATCACC
CGCTTTTTCGTGGATCGTGGCTTTCTGGAGATCAAATCCCCGATTCTGATTCCTCTGGAGTATATCGAGCGTATGGGCATCGACAA
TGATACCGAACTGAGCAAACAAATTTTCCGTGTGGATAAAAACTTCTGTCTGCGCCCTATGCTGGCACCAAATCTGTATAACTATC
TGCGCAAACTGGACCGTGCCCTGCCTGATCCTATCAAAATCTTCGAGATCGGCCCGTGTTATCGTAAAGAGTCCGACGGTAAAGA
ACATCTGGAGGAGTTTACCATGCTGAACTTTTGCCAAATGGGTTCAGGTTGTACTCGTGAGAACCTGGAAAGCATCATCACCGATT
TTCTGAACCACCTGGGCATTGACTTCAAAATTGTGGGCGACAGCTGTATGGTGTATGGCGACACCCTGGATGTCATGCACGGCGA
CCTGGAACTGTCTAGTGCCGTTGTTGGACCAATTCCGCTGGACCGTGAGTGGGGTATCGACAAACCGTGGATCGGAGCAGGATTC
GGTCTGGAACGCCTGCTGAAAGTGAAACACGACTTCAAAAACATCAAACGTGCCGCCCGTTCTGAATCGTATTATAACGGGATTT
CTACCAACCTGTAACCGCGGTACGCGATTCCATTGGACGCCCCCGGCCATCCGGGGGCGCTGATCAACAGGAGTGAGGGAAGCTT
GATCCGGGTTTTTCTCGAACGAGGCCGGTGGCGCAAGCCATCGGCCTTTTTTCATGCCCGGCGTGCCCTGTTGCACGGGAGGGCTA
AAGAAAATCGCCGGGGGGTCGATGCAATGGGTGTCGGAACTTCCACCCTCTGCCGGACCAACCGGGGGCGGTTCAGGACCGATA
TTGGCGAGTCCTCTTCGAAGCATGTAACCCACTGAAGAGGAAGAGAAAAAGAAAATGTTGATTTTTTCTCTAAAGCTCCGCCGGG

AAACGCCGATAAACACCATGAACGCGAATTCTTGGGGCACCTGAGCAAGCAGGCCGAGAGATCGCAAGCTCAGGTAACCGAAAT
AGGTCCTTTGGAGGAAATCACCATGGGCAGCAGCCATCATCATCATCATCACAGCTCTAGAGAGCTTTTCACTGGCGTTGTTCCCA
TCCTGGTCGAGCTGGACGGCGACGTAAACGGCCACAAGTTCAGCGTGTCCGGCGAGGGCGAGGGCGATGCCACCTACGGCAAGC
TGACCCTGAAGTTCATCTGCACCACCGGCAAGCTGCCCGTGCCCTGGCCCACCCTCGTGACCACCCTGACCTACGGCGTGCAGTGC
TTCAGCCGCTACCCCGACCACATGAAGCAGCACGACTTCTTCAAGTCCGCCATGCCCGAAGGCTACGTCCAGGAGCGCACCATCT
TCTTCAAGGACGACGGCAACTACAAGACCCGCGCCGAGGTGAAGTTCGAGGGCGACACCCTGGTGAACCGCATCGAGCTGAAGG
GCATCGACTTCAAGGAGGACGGCAACATCCTGGGGCACAAGCTGGAGTACAACTACAACAGCCACAACGTCTATATCATGGCCG
ACAAGCAGAAGAACGGCATCAAGGTGAACTTCAAGATCCGCCACAACATCGAGGACGGCAGCGTGCAGCTCGCCGACCACTACC
AGCAGAACACCCCCATCGGCGACGGCCCCGTGCTGCTGCCCGACAACCACTACCTGAGCACCCAGTCCGCCCTGAGCAAAGACCC
CAACGAGAAGCGCGATCACATGGTCCTGCTGGAGTTCGTGACCGCCGCCGGGATCTGACTCGAGCAAAGCCCGCCGAAAGGCGG
GCTTTTCTGTGGATCCGCGAATACCGCTTCCACAAACATTGCTCAAAAGTATCTCTTTGCTATATATCTCTGTGCTATATCCCTATA
TAACCTACCCATCCACCTTTCGCTCCTTGAACTTGCATCTAAACTCGACCTCTACATTTTTTATGTTTATCTCTAGTATTACTCTTTA
GACAAAAAAATTGTAGTAAGAACTATTCATAGAGTGAATCGAAAACAATACGAAAATGTAAACATTTCCTATACGTAGTATATAG
AGACAAAATAGAAGAAACCGTTCATAATTTTCTGACCAATGAAGAATCATCAACGCTATCACTTTCTGTTCACAAAGTATGCGCA
ATCCACATCGGTATAGAATATAATCGGGGATGCCTTTATCTTGAAAAAATGCACCCGCAGCTTCGCTAGTAATCAGTAAACGCGG
GAAGTGGAGTCAGGCTTTTTTTATGGAAGAGAAAATAGACACCAAAGTAGCCTTCTTCTAACCTTAACGGACCTACAGTGCAAAA
AGTTATCAAGAGACTGCATTATAGAGCGCACAAAGGAGAAAAAAAGTAATCTAAGATGCTTTGTTAGAAAAATAGCGCTCTCGG
GATGCATTTTTGTAGAACAAAAAAGAAGTATAGATTCTTTGTTGGTAAAATAGCGCTCTCGCGTTGCATTTCTGTTCTGTAAAAAT
GCAGCTCAGATTCTTTGTTTGAAAAATTAGCGCTCTCGTCGCGTTGCATTTTTGTTTTACAAAAATGAAGCACAGATTCTTCGTTGG
TAAAATAGCGCTTTCGCGTTGCATTTCTGTTCTGTAAAAATGCAGCTCAGATTCTTTGTTTGAAAAATTAGCGCTCTCGCGTTGCAT
TTTTGTTCTACAAAATGAAGCACAGATGCTTCGTTAACAAAGATATGCTATTGAAGTGCAAGATGGAAACGCAGAAAATGAACCG
GGGATGCGACGTGCAAGATTACCTATGCAATAGATGCAATAGTTTCTCCAGGAACCGAAATACATACATTGTCTTCCGTAAAGCG
CTAGACTATATATTATTATACAGGTTCAAATATACTATCTGTTTCAGGGAAAACTCCCAGGTTCGGATGTTCAAAATTCAATGATG
GGTAACAAGTACGATCGTAAATCTGTAAAACAGTTTGTCGGATATTAGGCTGTATCTCCTCAAAGCGTATTCGAATATCATTGAGA
AGCTGCAGCGTCACATCGGATAATAATGATGGCAGCCATTGTAGAAGTGCCTTTTGCATTTCTAGTCTCTTTCTCGGTCTAGCTAG
TTTTACTACATCGCGAAGATAGAATCTTAGATCACACTGCCTTTGCTGAGCTGGATCAATAGAGTAACAAAAGAGTGGTAAGGCC
TCGTTAAAGGACAAGGACCTGAGCGGAAGTGTATCGTACAGTAGACGGAGTATCTAGTATAGTCTATAGTCCGTGGAATTAATTC
TCATCTTTGACAGCTTATCATCGATAAGCTAGCTTTTCAATTCAATTCATCATTTTTTTTTTATTCTTTTTTTTGATTTCGGTTTCTTT
GAAATTTTTTTGATTCGGTAATCTCCGAACAGAAGGAAGAACGAAGGAAGGAGCACAGACTTAGATTGGTATATATACGCATATG
TAGTGTTGAAGAAACATGAAATTGCCCAGTATTCTTAACCCAACTGCACAGAACAAAAACCTGCAGGAAACGAAGATAAATCAT
GTCGAAAGCTACATATAAGGAACGTGCTGCTACTCATCCTAGTCCTGTTGCTGCCAAGCTATTTAATATCATGCACGAAAAGCAA
ACAAACTTGTGTGCTTCATTGGATGTTCGTACCACCAAGGAATTACTGGAGTTAGTTGAAGCATTAGGTCCCAAAATTTGTTTACT
AAAAACACATGTGGATATCTTGACTGATTTTTCCATGGAGGGCACAGTTAAGCCGCTAAAGGCATTATCCGCCAAGTACAATTTTT
TACTCTTCGAAGACAGAAAATTTGCTGACATTGGTAATACAGTCAAATTGCAGTACTCTGCGGGTGTATACAGAATAGCAGAATG
GGCAGACATTACGAATGCACACGGTGTGGTGGGCCCAGGTATTGTTAGCGGTTTGAAGCAGGCGGCAGAAGAAGTAACAAAGGA

ACCTAGAGGCCTTTTGATGTTAGCAGAATTGTCATGCAAGGGCTCCCTATCTACTGGAGAATATACTAAGGGTACTGTTGACATTG
CGAAGAGCGACAAAGATTTTGTTATCGGCTTTATTGCTCAAAGAGACATGGGTGGAAGAGATGAAGGTTACGATTGGTTGATTAT
GACACCCGGTGTGGGTTTAGATGACAAGGGAGACGCATTGGGTCAACAGTATAGAACCGTGGATGATGTGGTCTCTACAGGATCT
GACATTATTATTGTTGGAAGAGGACTATTTGCAAAGGGAAGGGATGCTAAGGTAGAGGGTGAACGTTACAGAAAAGCAGGCTGG
GAAGCATATTTGAGAAGATGCGGCCAGCAAAACTACTAGTGTATCAGCTCACTCAAAGGCGGTAATACGGTTATCCACAGAATCA
GGGGATAACGCAGGAAAGAACATGTGAGCAAAAGGCCAGCAAAAGGCCAGGAACCGTAAAAAGGCCGCGTTGCTGGCGTTTTTC
CATAGGCTCCGCCCCCCTGACGAGCATCACAAAAATCGACGCTCAAGTCAGAGGTGGCGAAACCCGACAGGACTATAAAGATAC
CAGGCGTTTCCCCCTGGAAGCTCCCTCGTGCGCTCTCCTGTTCCGACCCTGCCGCTTACCGGATACCTGTCCGCCTTTCTCCCTTCG
GGAAGCGTGGCGCTTTCTCATAGCTCACGCTGTAGGTATCTCAGTTCGGTGTAGGTCGTTCGCTCCAAGCTGGGCTGTGTGCACGA
ACCCCCCGTTCAGCCCGACCGCTGCGCCTTATCCGGTAACTATCGTCTTGAGTCCAACCCGGTAAGACACGACTTATCGCCACTGG
CAGCAGCCACTGGTAACAGGATTAGCAGAGCGAGGTATGTAGGCGGTGCTACAGAGTTCTTGAAGTGGTGGCCTAACTACGGCTA
CACTAGAAGGACAGTATTTGGTATCTGCGCTCTGCTGAAGCCAGTTACCTTCGGAAAAAGAGTTGGTAGCTCTTGATCCGGCAAA
CAAACCACCGCTGGTAGCGGTGGTTTTTTTGTTTGCAAGCAGCAGATTACGCGCAGAAAAAAAGGATCTCAAGAAGATCCTTTGA
TCTTTTCTACGGGGTCTGACGCTCAGTGGAACGAAAACTCACGTTAAGGGATTTTGGTCATGAGATTATCAAAAAGGATCTTCACC
TAGATCCTTTTAAATTAAAAATGAAGTTTTAAATCAATCTAAAGTATATATGAGTAAACTTGGTCTGACAGTTACCAATGCTTAAT
CAGTGAGGCACCTATCTCAGCGATCTGTCTATTTCGTTCATCCATAGTTGCCTGACTCCCCGTCGTGTAGATAACTACGATACGGG
AGGGCTTACCATCTGGCCCCAGTGCTGCAATGATACCGCGAGACCCACGCTCACCGGCTCCAGATTTATCAGCAATAAACCAGCC
AGCCGGAAGGGCCGAGCGCAGAAGTGGTCCTGCAACTTTATCCGCCTCCATCCAGTCTATTAATTGTTGCCGGGAAGCTAGAGTA
AGTAGTTCGCCAGTTAATAGTTTGCGCAACGTTGTTGCCATTGCTACAGGCATCGTGGTGTCACGCTCGTCGTTTGGTATGGCTTC
ATTCAGCTCCGGTTCCCAACGATCAAGGCGAGTTACATGATCCCCCATGTTGTGCAAAAAAGCGGTTAGCTCCTTCGGTCCTCCGA
TCGTTGTCAGAAGTAAGTTGGCCGCAGTGTTATCACTCATGGTTATGGCAGCACTGCATAATTCTCTTACTGTCATGCCATCCGTA
AGATGCTTTTCTGTGACTGGTGAGTACTCAACCAAGTCATTCTGAGAATAGTGTATGCGGCGACCGAGTTGCTCTTGCCCGGCGTC
AATACGGGATAATACCGCGCCACATAGCAGAACTTTAAAAGTGCTCATCATTGGAAAACGTTCTTCGGGGCGAAAACTCTCAAGG
ATCTTACCGCTGTTGAGATCCAGTTCGATGTAACCCACTCGTGCACCCAACTGATCTTCAGCATCTTTTACTTTCACCAGCGTTTCT
GGGTGAGCAAAAACAGGAAGGCAAAATGCCGCAAAAAAGGGAATAAGGGCGACACGGAAATGTTGAATACTCATACTCTTCCTT
TTTCAATATTATTGAAGCATTTATCAGGGTTATTGTCTCATGAGCGGATACATATTTGAATGTATTTAGAAAAATAAACAAATAGG
GGTTCCGCGCACATTTCCCCGAAAAGTGCCACCTGACGTCTAAGAAACCATTATTATCATGACATTAACCTATAAAAATAGGCGT
ATCACGAGGCCCTTTCGTCTCGCGCGTTTCGGTGATGACGGTGAAAACCTCTGACACATGCAGCTCCCGGAGACGGTCACAGCTT
GTCTGTAAGCGGATGCCGGGAGCAGACAAGCCCGTCAGGGCGCGTCAGCGGGTGTTGGCGGGTGTCGGGGCTGGCTTAACTATGC
GGCATCAGAGCAGATTGTACTGAGAGTGCACCATATGCGGTGTGAAATACCGCACAGATGCGTAAGGAGAAAATACCGCATCAG
GCGGCAATGGCAACAACGTTGCGCAAACTATTAACTGGCGAACTACTTACTCTAGCTTCCCGGCAACAATTAATAGACTGGATGG
AGGCGGATAAAGTTGCAGGACCACTTCTGCGCTCGGCCCTTCCGGCTGGCTGGTTTATTGCTGATAAATCTGGAGCCGGTGAGCGT
GGATCTCGCGGTATCATTGCAGCACTGGGGCCAGATGGTAAGCCCTCCCGTATCGTAGTTATCTACACGACGGGGAGTCAGGCAA
CTATGGATGAACGAAATAGACAGATCGCTGAGATAGGTGCCTCACTGATTAAGCATTGGTAACTGTCAGACCAAGTTTACTCATA
TATACTTTAGATTGATTTAAAACTTCATTTTTAATTTAAAAGGATCTAGGTGAAGATCCTTTTTGATAATCTCATGACCAAAATCCC

TTAACGTGAGTTTTCGTTCCACTGAGCGTCAGACCCCAATTACACGCCACTGGCTGTGCTTGCTGGGGTGACGGTGGCAACGGTGG
CGGCCTTGCTGGGCTATCGCGTTGGAAAGAAACGAGGGAAAGGGGACTGATAAACCGGTCTTAGCCCCTCCCCTTGGTGTCCAAC
CGCTCTGTAGGCCTCTCAGGCGCCGCTGGTGCCGCTGGTTGGACGCCAAGGGTGAATCCGCCTCGATACCCTGATTACTCGCTTCC
TGCGCCCTCTCAGGCGGCGATAGGGGACTGGTAAAACGGGGATTGCCCAGACGCCTCCCCCGCCCCTTCAGGGGCACAAATGCGG
CCCCAACGGGGCCACGTAGTGGTGCGTTTTTTGCGTTTCCACCCTTTTCTTCCTTTTCCCTTTTAAACCTTTTAGGACGTCTACAGGC
CACGTAATCCGTGGCCTGTAGAGTTTAAAAAGGGACGGATTTGTTGCCATTAAGGGACGGATTTGTTGTTAAGAAGGGACGGATT
TGTTGTTGTAAAGGGACGGATTTGTTGTATTGTGGGACGCAGATACAGTGTCCCCTTATACACAAGGAATGTCGAACGTGGCCTCA
CCCCCAATGGTTTACAAAAGCAATGCCCTGGTCGAGGCCGCGTATCGCCTCAGTGTTCAGGAACAGCGGATCGTTCTGGCCTGTA
TTAGCCAGGTGAAGAGGAGCGAGCCTGTCACCGATGAAGTGATGTATTCAGTGACGGCGGAGGACATAGCGACGATGGCGGGTG
TCCCTATCGAATCTTCCTACAACCAGCTCAAAGAAGCGGCCCTGCGCCTGAAACGGCGGGAAGTCCGGTTAACCCAAGAGCCCAA
TGGCAAGGGGAAAAGACCGAGTGTGATGATTACCGGCTGGGTGCAAACAATCATCTACCGGGAGGGTGAGGGCCGTGTAGAACT
CAGGTTCACCAAAGACATGCTGCCGTACCTGACGGAACTCACCAAACAGTTCACCAAATACGCCTTGGCTGACGTGGCCAAGATG
GACAGCACCCACGCGATCAGGCTTTACGAGCTGCTCATGCAATGGGACAGCATCGGCCAGCGCGAAAT

PylRS gene:

ATGGATAAAAAACCACTAAACACTCTGATCTCTGCTACTGGTCTGTGGATGAGTCGTACCGGAACCATTCATAAAATCAAACACC
ACGAGGTTAGCCGTTCGAAAATCTATATTGAGATGGCGTGTGGCGATCATCTGGTTGTGAACAATAGCCGCTCTTCTCGTACAGCA
CGTGCACTGCGTCACCACAAATATCGTAAAACCTGTAAACGTTGCCGTGTGTCCGATGAGGATCTGAACAAATTCCTGACAAAAG
CCAATGAGGACCAAACAAGCGTGAAAGTGAAAGTCGTTAGCGCTCCTACCCGTACTAAAAAAGCAATGCCGAAATCCGTTGCTC
GTGCCCCTAAACCACTGGAAAACACTGAAGCAGCACAGGCACAGCCGTCTGGAAGCAAATTCTCTCCGGCCATTCCTGTTTCTAC
CCAGGAGTCCGTTTCTGTTCCAGCAAGTGTGAGCACCAGCATTAGCAGTATTAGCACCGGTGCCACCGCTAGCGCCCTGGTTAAA
GGCAATACCAATCCGATTACAAGCATGTCTGCCCCGGTTCAAGCATCAGCTCCAGCACTGACAAAATCCCAAACCGATCGTCTGG
AGGTTCTGCTGAATCCGAAAGACGAAATCAGCCTGAATTCCGGCAAACCGTTTCGTGAACTGGAGAGCGAACTGCTGTCACGTCG
TAAAAAAGACCTGCAACAAATCTATGCCGAAGAACGTGAGAACTATCTGGGGAAACTGGAACGTGAAATCACCCGCTTTTTCGTG
GATCGTGGCTTTCTGGAGATCAAATCCCCGATTCTGATTCCTCTGGAGTATATCGAGCGTATGGGCATCGACAATGATACCGAACT
GAGCAAACAAATTTTCCGTGTGGATAAAAACTTCTGTCTGCGCCCTATGCTGGCACCAAATCTGTATAACTATCTGCGCAAACTGG
ACCGTGCCCTGCCTGATCCTATCAAAATCTTCGAGATCGGCCCGTGTTATCGTAAAGAGTCCGACGGTAAAGAACATCTGGAGGA
GTTTACCATGCTGAACTTTTGCCAAATGGGTTCAGGTTGTACTCGTGAGAACCTGGAAAGCATCATCACCGATTTTCTGAACCACC
TGGGCATTGACTTCAAAATTGTGGGCGACAGCTGTATGGTGTATGGCGACACCCTGGATGTCATGCACGGCGACCTGGAACTGTC
TAGTGCCGTTGTTGGACCAATTCCGCTGGACCGTGAGTGGGGTATCGACAAACCGTGGATCGGAGCAGGATTCGGTCTGGAACGC
CTGCTGAAAGTGAAACACGACTTCAAAAACATCAAACGTGCCGCCCGTTCTGAATCGTATTATAACGGGATTTCTACCAACCTGT
AA

Pyl tRNA:

GGAAACCTGATCATGTAGATCGAATGGACTCTAAATCCGTTCAGCCGGGTTAGATTCCCGGGGTTTCCGCCA

deGFP NHis gene:

ATGGGCAGCAGCCATCATCATCATCATCACAGCTCTAGAGAGCTTTTCACTGGCGTTGTTCCCATCCTGGTCGAGCTGGACGGCGA
CGTAAACGGCCACAAGTTCAGCGTGTCCGGCGAGGGCGAGGGCGATGCCACCTACGGCAAGCTGACCCTGAAGTTCATCTGCACC
ACCGGCAAGCTGCCCGTGCCCTGGCCCACCCTCGTGACCACCCTGACCTACGGCGTGCAGTGCTTCAGCCGCTACCCCGACCACA
TGAAGCAGCACGACTTCTTCAAGTCCGCCATGCCCGAAGGCTACGTCCAGGAGCGCACCATCTTCTTCAAGGACGACGGCAACTA
CAAGACCCGCGCCGAGGTGAAGTTCGAGGGCGACACCCTGGTGAACCGCATCGAGCTGAAGGGCATCGACTTCAAGGAGGACGG
CAACATCCTGGGGCACAAGCTGGAGTACAACTACAACAGCCACAACGTCTATATCATGGCCGACAAGCAGAAGAACGGCATCAA
GGTGAACTTCAAGATCCGCCACAACATCGAGGACGGCAGCGTGCAGCTCGCCGACCACTACCAGCAGAACACCCCCATCGGCGA
CGGCCCCGTGCTGCTGCCCGACAACCACTACCTGAGCACCCAGTCCGCCCTGAGCAAAGACCCCAACGAGAAGCGCGATCACATG
GTCCTGCTGGAGTTCGTGACCGCCGCCGGGATCTGA

pPaGE Pyl TAG *fliC* T248TAG:

TTATCCCCTGATTCTGTGGATAACCGTATTACCGCCTTTGAGTGAGCTGATACACTAGTAGTTTTGCTGGCCGCATCTTCAGAAATT
GACCAGCTGCGAAAGTGGTTTCAACTGGAAGGCCGGTATCCCTCGATCAAGGACTTCAAGTTGCGAGTGCTTGATCCAGCCGTGA
CGCAGATCAACGAGCACAGCCCGCTACAGGTGGAGTGGGCGCAGCGAAAGACCGGGCGCAAGGTCACACATCTGTTGTTCAGTT
TTGGACCGAAGAAGCCCGCCAAGGCGGTGGGTAAGGCCCCAGCGAAGCGCAAGGCCGGGAAGATTTCAGATGCTGAGATCGCGA
AACAGGCTCGCCCTGGTGAGACATGGGAAGCGGCCCGCGCTCGACTAACCCAGATGCCGCTGGATCTGGCCTAGAGGCCGTGGC
CACCACGGCCCGGCCTGCCTTTCGGATCCTAGAGGTAGGTCACCTTTTCTTCTTTAGCGTTTGCTGCCATTGCTCTATTGAAGCGCG
AAAACGCTTCGTCGGTCAGCTTCGGCAACAGAATAATTTTTATCGCATCAGGGGTTTACAAGCAGAATACGTACTCGTATAGTTCG
CGCCCACAACGTTGCTGAGACGTTGTAACACGGAAACGATGAGCGATCATCGACTCCAGGGAAACCTGATCATGTAGATCGAATG
GACTCTAAATCCGTTCAGCCGGGTTAGATTCCCGGGGTTTCCGCCAATATTTCAGACAAAGCCGGTCATTAGACCGTTTTTGTTTC
AGCAGTTCAGCCACTACCAGGCATTGAAACTGCGACAAGGTTTGCAACACCCAAGTCGATGAGCGATCATCGCACAATGCTGAAA
AGCACGAACACCTCTGAAGGTACCACTGGGCAGCCAGCCATCGACGGCCCATAGGGCAAGCTTAGCGTTCCCGTTCGGCTGAACC
GCCCCCGGCGACAAACGGCGCGGGCTATCGACAGGAGTGGGGGTCATGTATGCTACGAGGCTTTCGAACAGTCACGTCTAAGTA
GCGCCCCGCGGATGGATAAAAAACCACTAAACACTCTGATCTCTGCTACTGGTCTGTGGATGAGTCGTACCGGAACCATTCATAA
AATCAAACACCACGAGGTTAGCCGTTCGAAAATCTATATTGAGATGGCGTGTGGCGATCATCTGGTTGTGAACAATAGCCGCTCT
TCTCGTACAGCACGTGCACTGCGTCACCACAAATATCGTAAAACCTGTAAACGTTGCCGTGTGTCCGATGAGGATCTGAACAAAT
TCCTGACAAAAGCCAATGAGGACCAAACAAGCGTGAAAGTGAAAGTCGTTAGCGCTCCTACCCGTACTAAAAAAGCAATGCCGA
AATCCGTTGCTCGTGCCCCTAAACCACTGGAAAACACTGAAGCAGCACAGGCACAGCCGTCTGGAAGCAAATTCTCTCCGGCCAT
TCCTGTTTCTACCCAGGAGTCCGTTTCTGTTCCAGCAAGTGTGAGCACCAGCATTAGCAGTATTAGCACCGGTGCCACCGCTAGCG

CCCTGGTTAAAGGCAATACCAATCCGATTACAAGCATGTCTGCCCCGGTTCAAGCATCAGCTCCAGCACTGACAAAATCCCAAAC
CGATCGTCTGGAGGTTCTGCTGAATCCGAAAGACGAAATCAGCCTGAATTCCGGCAAACCGTTTCGTGAACTGGAGAGCGAACTG
CTGTCACGTCGTAAAAAAGACCTGCAACAAATCTATGCCGAAGAACGTGAGAACTATCTGGGGAAACTGGAACGTGAAATCACC
CGCTTTTTCGTGGATCGTGGCTTTCTGGAGATCAAATCCCCGATTCTGATTCCTCTGGAGTATATCGAGCGTATGGGCATCGACAA
TGATACCGAACTGAGCAAACAAATTTTCCGTGTGGATAAAAACTTCTGTCTGCGCCCTATGCTGGCACCAAATCTGTATAACTATC
TGCGCAAACTGGACCGTGCCCTGCCTGATCCTATCAAAATCTTCGAGATCGGCCCGTGTTATCGTAAAGAGTCCGACGGTAAAGA
ACATCTGGAGGAGTTTACCATGCTGAACTTTTGCCAAATGGGTTCAGGTTGTACTCGTGAGAACCTGGAAAGCATCATCACCGATT
TTCTGAACCACCTGGGCATTGACTTCAAAATTGTGGGCGACAGCTGTATGGTGTATGGCGACACCCTGGATGTCATGCACGGCGA
CCTGGAACTGTCTAGTGCCGTTGTTGGACCAATTCCGCTGGACCGTGAGTGGGGTATCGACAAACCGTGGATCGGAGCAGGATTC
GGTCTGGAACGCCTGCTGAAAGTGAAACACGACTTCAAAAACATCAAACGTGCCGCCCGTTCTGAATCGTATTATAACGGGATTT
CTACCAACCTGTAACCGCGGTACGCGATTCCATTGGACGCCCCCGGCCATCCGGGGGCGCTGATCAACAGGAGTGAGGGAAGCTT
GATCCGGGTTTTTCTCGAACGAGGCCGGTGGCGCAAGCCATCGGCCTTTTTTCATGCCCGGCGTGCCCTGTTGCACGGGAGGGCTA
AAGAAAATCGCCGGGGGGTCGATGCAATGGGTGTCGGAACTTCCACCCTCTGCCGGACCAACCGGGGGCGGTTCAGGACCGATA
TTGGCGAGTCCTCTTCGAAGCATGTAACCCACTGAAGAGGAAGAGAAAAAGAAAATGTTGATTTTTTCTCTAAAGCTCCGCCGGG
AAACGCCGATAAACACCATGAACGCGAATTCTTGGGGCACCTGAGCAAGCAGGCCGAGAGATCGCAAGCTCAGGTAACCGAAAT
AGGTCCTTTGGAGGAAATCACCATGGCCCTTACAGTCAACACGAACATTGCTTCCCTGAACACTCAGCGCAACCTGAATGCTTCTT
CCAACGACCTCAACACCTCGTTGCAGCGTCTGACCACCGGCTACCGCATCAACAGTGCCAAGGACGATGCTGCCGGCCTGCAGAT
CTCCAACCGCCTGTCCAACCAGATCAGCGGTCTGAACGTTGCCACCCGCAACGCCAACGACGGCATCTCCCTGGCGCAGACCGCT
GAAGGTGCCCTGCAGCAGTCCACCAATATCCTGCAGCGTATCCGCGACCTGGCCCTGCAATCCGCCAACGGCTCCAACAGCGACG
CCGACCGTGCCGCCCTGCAGAAAGAAGTCGCTGCGCAACAGGCCGAACTGACCCGTATCTCCGATACCACCACCTTCGGTGGCCG
CAAGCTGCTCGACGGCTCCTTCGGCACCACCAGCTTCCAGGTCGGTTCCAACGCCTACGAGACCATTGACATCAGCCTGCAGAAT
GCCTCTGCCAGCGCCATCGGTTCTTACCAGGTCGGCAGCAACGGCGCGGGTACCGTCGCCAGCGTAGCGGGCACCGCGACCGCTT
CGGGCATCGCCTCGGGCACCGTCAACCTGGTCGGTGGCGGTCAGGTGAAGAACATCGCCATCGCCGCCGGCGATAGCGCCAAGG
CCATCGCCGAGAAGATGGACGGTGCGATCCCGAACCTGTCGGCTCGTGCCCGTACCGTGTTCACCGCTGATGTCAGCGGCGTGTA
GGGTGGTTCGCTGAACTTCGACGTAACCGTTGGCAGCAACACCGTGAGCCTGGCAGGCGTGACCTCCACTCAGGATCTGGCCGAC
CAACTGAACTCCAACTCGTCGAAGCTGGGCATCACTGCCAGCATCAACGACAAGGGTGTACTGACCATCACCTCCGCTACCGGCG
AGAACGTCAAGTTCGGTGCGCAGACCGGTACCGCTACTGCCGGTCAGGTCGCAGTGAAGGTCCAGGGTTCCGACGGCAAGTTCGA
AGCGGCCGCCAAGAACGTGGTAGCTGCCGGTACTGCCGCTACCACCACCATCGTGACCGGCTACGTGCAACTGAACTCGCCGACC
GCCTACTCGGTCAGCGGTACCGGCACCCAGGCTTCGCAGGTCTTCGGCAACGCCAGCGCCGCGCAGAAGAGCAGCGTTGCCAGCG
TCGACATCTCCACTGCCGACGGCGCCCAGAACGCCATCGCGGTAGTCGATAACGCCCTGGCTGCGATCGACGCCCAGCGTGCTGA
CCTCGGTGCTGTTCAGAACCGCTTCAAGAACACTATCGACAACCTGACCAACATCTCGGAAAACGCTACCAACGCTCGTAGCCGC
ATCAAGGACACCGACTTCGCTGCCGAAACCGCGGCGCTGTCGAAGAACCAGGTGCTGCAACAGGCCGGTACCGCGATCCTGGCC
CAGGCCAACCAGCTGCCGCAGGCGGTCCTGAGCCTGCTGCGCTAAGCCCGGGAACGGTCACTCACGCGGTACTGGGAGGAAGGG
GTGACCCTTCCTCCCTTTTCCCCATGGAGGGCACAGTTAAGCCGCTAAAGGCATTATCCGCCAAGTACAATTTTTTACTCTTCGAA
GACAGAAAATTTGCTGACATTGGTAATACAGTCAAATTGCAGTACTCTGCGGGTGTATACAGAATAGCAGAATGGGCAGACATTA

CGAATGCACACGGTGTGGTGGGCCCAGGTATTGTTAGCGGTTTGAAGCAGGCGGCAGAAGAAGTAACAAAGGAACCTAGAGGCC
TTTTGATGTTAGCAGAATTGTCATGCAAGGGCTCCCTATCTACTGGAGAATATACTAAGGGTACTGTTGACATTGCGAAGAGCGAC
AAAGATTTTGTTATCGGCTTTATTGCTCAAAGAGACATGGGTGGAAGAGATGAAGGTTACGATTGGTTGATTATGACACCCGGTG
TGGGTTTAGATGACAAGGGAGACGCATTGGGTCAACAGTATAGAACCGTGGATGATGTGGTCTCTACAGGATCTGACATTATTAT
TGTTGGAAGAGGACTATTTGCAAAGGGAAGGGATGCTAAGGTAGAGGGTGAACGTTACAGAAAAGCAGGCTGGGAAGCATATTT
GAGAAGATGCGGCCAGCAAAACTACTAGTGTATCAGCTCACTCAAAGGCGGTAATACGGTTATCCACAGAATCAGGGGATAACG
CAGGAAAGAACATGTGAGCAAAAGGCCAGCAAAAGGCCAGGAACCGTAAAAAGGCCGCGTTGCTGGCGTTTTTCCATAGGCTCC
GCCCCCCTGACGAGCATCACAAAAATCGACGCTCAAGTCAGAGGTGGCGAAACCCGACAGGACTATAAAGATACCAGGCGTTTC
CCCCTGGAAGCTCCCTCGTGCGCTCTCCTGTTCCGACCCTGCCGCTTACCGGATACCTGTCCGCCTTTCTCCCTTCGGGAAGCGTGG
CGCTTTCTCATAGCTCACGCTGTAGGTATCTCAGTTCGGTGTAGGTCGTTCGCTCCAAGCTGGGCTGTGTGCACGAACCCCCCGTT
CAGCCCGACCGCTGCGCCTTATCCGGTAACTATCGTCTTGAGTCCAACCCGGTAAGACACGACTTATCGCCACTGGCAGCAGCCA
CTGGTAACAGGATTAGCAGAGCGAGGTATGTAGGCGGTGCTACAGAGTTCTTGAAGTGGTGGCCTAACTACGGCTACACTAGAAG
GACAGTATTTGGTATCTGCGCTCTGCTGAAGCCAGTTACCTTCGGAAAAAGAGTTGGTAGCTCTTGATCCGGCAAACAAACCACC
GCTGGTAGCGGTGGTTTTTTTGTTTGCAAGCAGCAGATTACGCGCAGAAAAAAAGGATCTCAAGAAGATCCTTTGATCTTTTCTAC
GGGGTCTGACGCTCAGTGGAACGAAAACTCACGTTAAGGGATTTTGGTCATGAGATTATCAAAAAGGATCTTCACCTAGATCCTT
TTAAATTAAAAATGAAGTTTTAAATCAATCTAAAGTATATATGAGTAAACTTGGTCTGACAGTTACCAATGCTTAATCAGTGAGGC
ACCTATCTCAGCGATCTGTCTATTTCGTTCATCCATAGTTGCCTGACTCCCCGTCGTGTAGATAACTACGATACGGGAGGGCTTAC
CATCTGGCCCCAGTGCTGCAATGATACCGCGAGACCCACGCTCACCGGCTCCAGATTTATCAGCAATAAACCAGCCAGCCGGAAG
GGCCGAGCGCAGAAGTGGTCCTGCAACTTTATCCGCCTCCATCCAGTCTATTAATTGTTGCCGGGAAGCTAGAGTAAGTAGTTCGC
CAGTTAATAGTTTGCGCAACGTTGTTGCCATTGCTACAGGCATCGTGGTGTCACGCTCGTCGTTTGGTATGGCTTCATTCAGCTCCG
GTTCCCAACGATCAAGGCGAGTTACATGATCCCCCATGTTGTGCAAAAAAGCGGTTAGCTCCTTCGGTCCTCCGATCGTTGTCAGA
AGTAAGTTGGCCGCAGTGTTATCACTCATGGTTATGGCAGCACTGCATAATTCTCTTACTGTCATGCCATCCGTAAGATGCTTTTCT
GTGACTGGTGAGTACTCAACCAAGTCATTCTGAGAATAGTGTATGCGGCGACCGAGTTGCTCTTGCCCGGCGTCAATACGGGATA
ATACCGCGCCACATAGCAGAACTTTAAAAGTGCTCATCATTGGAAAACGTTCTTCGGGGCGAAAACTCTCAAGGATCTTACCGCT
GTTGAGATCCAGTTCGATGTAACCCACTCGTGCACCCAACTGATCTTCAGCATCTTTTACTTTCACCAGCGTTTCTGGGTGAGCAA
AAACAGGAAGGCAAAATGCCGCAAAAAAGGGAATAAGGGCGACACGGAAATGTTGAATACTCATACTCTTCCTTTTTCAATATTA
TTGAAGCATTTATCAGGGTTATTGTCTCATGAGCGGATACATATTTGAATGTATTTAGAAAAATAAACAAATAGGGGTTCCGCGCA
CATTTCCCCGAAAAGTGCCACCTGACGTCTAAGAAACCATTATTATCATGACATTAACCTATAAAAATAGGCGTATCACGAGGCC
CTTTCGTCTCGCGCGTTTCGGTGATGACGGTGAAAACCTCTGACACATGCAGCTCCCGGAGACGGTCACAGCTTGTCTGTAAGCGG
ATGCCGGGAGCAGACAAGCCCGTCAGGGCGCGTCAGCGGGTGTTGGCGGGTGTCGGGGCTGGCTTAACTATGCGGCATCAGAGC
AGATTGTACTGAGAGTGCACCATATGCGGTGTGAAATACCGCACAGATGCGTAAGGAGAAAATACCGCATCAGGCGGCAATGGC
AACAACGTTGCGCAAACTATTAACTGGCGAACTACTTACTCTAGCTTCCCGGCAACAATTAATAGACTGGATGGAGGCGGATAAA
GTTGCAGGACCACTTCTGCGCTCGGCCCTTCCGGCTGGCTGGTTTATTGCTGATAAATCTGGAGCCGGTGAGCGTGGATCTCGCGG
TATCATTGCAGCACTGGGGCCAGATGGTAAGCCCTCCCGTATCGTAGTTATCTACACGACGGGGAGTCAGGCAACTATGGATGAA
CGAAATAGACAGATCGCTGAGATAGGTGCCTCACTGATTAAGCATTGGTAACTGTCAGACCAAGTTTACTCATATATACTTTAGAT

TGATTTAAAACTTCATTTTTAATTTAAAAGGATCTAGGTGAAGATCCTTTTTGATAATCTCATGACCAAAATCCCTTAACGTGAGTT
TTCGTTCCACTGAGCGTCAGACCCCAATTACACGCCACTGGCTGTGCTTGCTGGGGTGACGGTGGCAACGGTGGCGGCCTTGCTG
GGCTATCGCGTTGGAAAGAAACGAGGGAAAGGGGACTGATAAACCGGTCTTAGCCCCTCCCCTTGGTGTCCAACCGCTCTGTAGG
CCTCTCAGGCGCCGCTGGTGCCGCTGGTTGGACGCCAAGGGTGAATCCGCCTCGATACCCTGATTACTCGCTTCCTGCGCCCTCTC
AGGCGGCGATAGGGGACTGGTAAAACGGGGATTGCCCAGACGCCTCCCCCGCCCCTTCAGGGGCACAAATGCGGCCCCAACGGG
GCCACGTAGTGGTGCGTTTTTTGCGTTTCCACCCTTTTCTTCCTTTTCCCTTTTAAACCTTTTAGGACGTCTACAGGCCACGTAATCC
GTGGCCTGTAGAGTTTAAAAAGGGACGGATTTGTTGCCATTAAGGGACGGATTTGTTGTTAAGAAGGGACGGATTTGTTGTTGTA
AAGGGACGGATTTGTTGTATTGTGGGACGCAGATACAGTGTCCCCTTATACACAAGGAATGTCGAACGTGGCCTCACCCCCAATG
GTTTACAAAAGCAATGCCCTGGTCGAGGCCGCGTATCGCCTCAGTGTTCAGGAACAGCGGATCGTTCTGGCCTGTATTAGCCAGG
TGAAGAGGAGCGAGCCTGTCACCGATGAAGTGATGTATTCAGTGACGGCGGAGGACATAGCGACGATGGCGGGTGTCCCTATCG
AATCTTCCTACAACCAGCTCAAAGAAGCGGCCCTGCGCCTGAAACGGCGGGAAGTCCGGTTAACCCAAGAGCCCAATGGCAAGG
GGAAAAGACCGAGTGTGATGATTACCGGCTGGGTGCAAACAATCATCTACCGGGAGGGTGAGGGCCGTGTAGAACTCAGGTTCA
CCAAAGACATGCTGCCGTACCTGACGGAACTCACCAAACAGTTCACCAAATACGCCTTGGCTGACGTGGCCAAGATGGACAGCAC
CCACGCGATCAGGCTTTACGAGCTGCTCATGCAATGGGACAGCATCGGCCAGCGCGAAAT

*fliC* gene:

ATGGCCCTTACAGTCAACACGAACATTGCTTCCCTGAACACTCAGCGCAACCTGAATGCTTCTTCCAACGACCTCAACACCTCGTT
GCAGCGTCTGACCACCGGCTACCGCATCAACAGTGCCAAGGACGATGCTGCCGGCCTGCAGATCTCCAACCGCCTGTCCAACCAG
ATCAGCGGTCTGAACGTTGCCACCCGCAACGCCAACGACGGCATCTCCCTGGCGCAGACCGCTGAAGGTGCCCTGCAGCAGTCCA
CCAATATCCTGCAGCGTATCCGCGACCTGGCCCTGCAATCCGCCAACGGCTCCAACAGCGACGCCGACCGTGCCGCCCTGCAGAA
AGAAGTCGCTGCGCAACAGGCCGAACTGACCCGTATCTCCGATACCACCACCTTCGGTGGCCGCAAGCTGCTCGACGGCTCCTTC
GGCACCACCAGCTTCCAGGTCGGTTCCAACGCCTACGAGACCATTGACATCAGCCTGCAGAATGCCTCTGCCAGCGCCATCGGTT
CTTACCAGGTCGGCAGCAACGGCGCGGGTACCGTCGCCAGCGTAGCGGGCACCGCGACCGCTTCGGGCATCGCCTCGGGCACCGT
CAACCTGGTCGGTGGCGGTCAGGTGAAGAACATCGCCATCGCCGCCGGCGATAGCGCCAAGGCCATCGCCGAGAAGATGGACGG
TGCGATCCCGAACCTGTCGGCTCGTGCCCGTACCGTGTTCACCGCTGATGTCAGCGGCGTGACCGGTGGTTCGCTGAACTTCGACG
TAACCGTTGGCAGCAACACCGTGAGCCTGGCAGGCGTGACCTCCACTCAGGATCTGGCCGACCAACTGAACTCCAACTCGTCGAA
GCTGGGCATCACTGCCAGCATCAACGACAAGGGTGTACTGACCATCACCTCCGCTACCGGCGAGAACGTCAAGTTCGGTGCGCAG
ACCGGTACCGCTACTGCCGGTCAGGTCGCAGTGAAGGTCCAGGGTTCCGACGGCAAGTTCGAAGCGGCCGCCAAGAACGTGGTA
GCTGCCGGTACTGCCGCTACCACCACCATCGTGACCGGCTACGTGCAACTGAACTCGCCGACCGCCTACTCGGTCAGCGGTACCG
GCACCCAGGCTTCGCAGGTCTTCGGCAACGCCAGCGCCGCGCAGAAGAGCAGCGTTGCCAGCGTCGACATCTCCACTGCCGACGG
CGCCCAGAACGCCATCGCGGTAGTCGATAACGCCCTGGCTGCGATCGACGCCCAGCGTGCTGACCTCGGTGCTGTTCAGAACCGC
TTCAAGAACACTATCGACAACCTGACCAACATCTCGGAAAACGCTACCAACGCTCGTAGCCGCATCAAGGACACCGACTTCGCTG
CCGAAACCGCGGCGCTGTCGAAGAACCAGGTGCTGCAACAGGCCGGTACCGCGATCCTGGCCCAGGCCAACCAGCTGCCGCAGG
CGGTCCTGAGCCTGCTGCGCTAA

**Table S1.** Primers list:

| Number | Purpose | Sequence |
|---|---|---|
| 1 | pMRP9-1 backbone, 5' overhang of 2µ and URA3. Forward for yeast assembly. | TGCCGCTGGATCTGGCCTAGAGGCCGTGGCCACCA CGGCCCGGCCTGCCTTTCGGATCCGCGAATACCGCT TCCACAAAC |
| 2 | pMRP9-1 backbone, 5' overhang of 2µ URA3. Reverse for yeast assembly. | TTATCCCCTGATTCTGTGGATAACCGTATTACCGCCTT TGAGTGAGCTGATACACTAGTAGTTTGCTGGCCGCA TCTTC |
| 3 | Leu-tRNA promoter, 5' overhang of replicase gene from pMRP9-1. Forward for yeast assembly. | CCGCTGGATCTGGCCTAGAGGCCGTGGCCACCACGGC CCGGCCTGCCTTTC GGATCC TAGAGGTAGGTCACCTTTTCTTC |
| 4 | Leu-tRNA promoter, 5' overhang of Pyl- | CCGGGAATCTAACCCGGCTGAACGGATTTAGAGTCCA TTCGATCTACATGATCAGGTTTCC CTGGAGTCGATGATCGCTC |

| | | |
|---|---|---|
| | tRNA. Reverse for yeast assembly. | |
| 5 | Leu-tRNA terminator, 5' overhang of Pyl-tRNA. Forward for yeast assembly. | CATGTAGATCGAATGGACTCTAAATCCGTTCAGCCGG GTTAGATTCCCGG GGTTTCCGCCA ATATTTCAGACAAAGCCGG |
| 6 | Leu-tRNA terminator, 5' overhang of LeuRS promoter. Reverse for yeast assembly. | TTGCCCTATGGGCCGTCGATGGCTGGCTGCCCAGTGG TACCTTCAGAGGTGTTCGTGCTTTTCAGCATTG TGCG |
| 7 | LeuRS promoter. Forward for yeast assembly. | CGAACACCTCTGAAGGTACC |
| 8 | LeuRS promoter, 5' overhang of PylRS. Reverse for yeast assembly. | GACTCATCCACAGACCAGTAGCAGAGATCAGAGTGTT TAGTGGTTTTTATCCATCCGCGGGGCGCTACTTA GACGTGAC |
| 9 | PylRS. Forward for yeast assembly. | ATGGATAAAAAACCACTAAACACTC |

| | | |
|---|---|---|
| 10 | PylRS, 5' overhang of LeuRS terminator. Reverse for yeast assembly. | CACTCCTGTTGATCAGCGCCCCCGGATGGCCGGGGGCGTCCAATGGAATCGCGTACCGCGGTTACAGGTTGGTAGAAATCC |
| 11 | LeuRS terminator. Forward for yeast assembly. | TACGCGATTCCATTGGACGC |
| 12 | LeuRS terminator, 5' overhang of *fliC* promoter. Reverse for yeast assembly. | GCCGGGCATGAAAAAGGCCGATGGCTTGCGCCACCGGCCTCGTTCGAGAAAAACCCGGATCAAGCTTCCCTCACTCCTG |
| 13 | *fliC* promoter. Forward for yeast assembly. | ATCCGGGTTTTTCTCGAACG |
| 14 | *fliC* promoter, 5' overhang of N-terminal his-tag deGFP. Reverse for yeast assembly. | AACAACGCCAGTGAAAAGCTCTCTAGAGCTGTGATGATGATGATGATGGCTGCTGCCCATGGTGATTTCCTCCAAAGGAC |
| 15 | N-terminal his-tag deGFP. Forward for yeast assembly | ATGGGCAGCAGCCATCATCATC |

| | | |
|---|---|---|
| | (for both WT and Y35TAG). | |
| 16 | T500 terminator, 5’ overhang of 2µ origin. Reverse for yeast assembly (For both WT and Y35TAG). | GAGATATATAGCAAAGAGATACTTTTGAGCAATGTTTGTGGAAGCGGTATTCGCGGATCCACAGAAAAGCCCGCCTTTCG |
| 17 | *fliC* gene, 5’ overhang of *fliC* promoter. Forward for Gibson assembly. | TCAGGTAACCGAAATAGGTCCTTTGGAGGAAATCACCATGGCCCTTACAGTCAACACGAAC |
| 18 | *fliC* terminator, 5’ overhang of mid URA3. Reverse for Gibson assembly. | TCAGGTAACCGAAATAGGTCCTTTGGAGGAAATCACCATGGCCCTTACAGTCAACACGAAC |
| 19 | *fliC* T248TAG mutation installation. Forward for Gibson assembly. | GATGTCAGCGGCGTGTAGGGTGGTTCGCTGAAC |

| | | |
|---|---|---|
| 20 | *fliC* T248TAG mutation installation. Reverse for Gibson assembly. | GTTCAGCGAACCACCCTACACGCCGCTGACATC |
| 21 | pACRISPR, 5' overhang for gRNA installation for *fliC* deletion. Forward for Gibson assembly. | TCGTATAATGTGTGGGCCATCGGTTCTTACCAGGTGTT TTAGAGCTAGAAATAGC |
| 22 | pACRISPR template upstream to homology installation. Reverse. | CGTCTTCAAGAATTCTGGCAC |
| 23 | *fliC* upstream region, 5' overhang for pACRISPR plasmid. Forward for Gibson assembly. | GGGAAACCTGTCGTGCCAGAATTCTTGAAGACGTGGA GGCGATCCGCGCGCAC |

| | | |
|---|---|---|
| 24 | *fliC* upstream region, 5' overhang for *fliC* end-gene. Reverse for Gibson assembly. | TTAGCGCAGCAGGCTCAGGACCGCCTGCGGGGTGATTTCCTCCAAAGGAC |
| 25 | *fliC* downstream region, 5' overhang for *fliC* end-gene. Forward for Gibson assembly. | CCGCAGGCGGTCCTGAGCCTGCTGCGCTAAGCCCGGGAACGGTCACTCAC |
| 26 | *fliC* downstream region, 5' overhang for pACRISPR plasmid. Reverse for Gibson assembly. | CCTATAAAAATAGGCGTATCACGAGGCCCTTTCCTATCCCTTCGATAGGGCAAG |
| 27 | pACRISPR template downstream to homology | AAAGGGCCTCGTGATACGCC |

| | | |
|---|---|---|
| | installation. Forward. | |
| 28 | pACRISPR, 5' overhang for gRNA installation for *fliC* deletion. Reverse for Gibson assembly. | TTCTAGCTCTAAAACACCTGGTAAGAACCGATGGCCCACACATTATACGAGCCGG |
| 29 | Genome screening for *fliC* upstream. Forward. | CTGTTGCACGGGAGGGCTAAAGAAAATCG |
| 30 | Genome screening for *fliC* downstream. Reverse. | ACGAAAGACTGGATGTCGCTGACGGC |
| 31 | Genome sequencing for *fliC* deleted region. Reverse. | AAGTGATATTTCCGACGTCC |
